# Whole Organism Profiling of the Timp Gene Family

**DOI:** 10.1101/2021.10.15.464572

**Authors:** David Peeney, Yu Fan, Sadeechya Gurung, Carolyn Lazaroff, Shashikala Ratnayake, Andrew Warner, Baktiar Karim, Daoud Meerzaman, William G. Stetler-Stevenson

**Affiliations:** Extracellular Matrix Pathology Section, Laboratory of Pathology, National Cancer Institute, National Institute of Health, Bethesda, Maryland; Computational Genomics and Bioinformatics Branch, Center for Biomedical Informatics & Information Technology, National Cancer Institute, National Institute of Health, Rockville, Maryland; Molecular Histopathology Laboratory, Frederick National Laboratory, National Cancer Institute, Frederick, Maryland

**Keywords:** TIMP, Extracellular Matrix, Matrisome, scRNAseq, Spatial Transcriptomics

## Abstract

Tissue inhibitor of metalloproteinases (TIMPs/Timps) are an endogenous family of widely expressed matrisome-associated proteins that were initially identified as inhibitors of matrix metalloproteinase activity (Metzincin family proteases). Consequently, TIMPs are often considered simply as protease inhibitors by many investigators. However, an evolving list of new metalloproteinase-independent functions for TIMP family members suggests that this concept is outdated. These novel TIMP functions include direct agonism/antagonism of multiple transmembrane receptors, as well as functional interactions with matrisome targets. While the family was fully identified over two decades ago, there has yet to be an in-depth study describing the expression of TIMPs in normal tissues of adult mammals. An understanding of the tissues and cell-types that express TIMPs 1 through 4, in both normal and disease states are important to contextualize the growing functional capabilities of TIMP proteins, which are often dismissed as non-canonical. Using publicly available single cell RNA sequencing data from the Tabula Muris Consortium, we analyzed approximately 100,000 murine cells across eighteen tissues from non-diseased organs, representing seventy-three annotated cell types, to define the diversity in Timp gene expression across healthy tissues. We describe the unique expression profiles across tissues and organ-specific cell types that all four Timp genes display. Within annotated cell-types, we identify clear and discrete cluster-specific patterns of Timp expression, particularly in cells of stromal and endothelial origins. RNA in-situ hybridization across four organs expands on the scRNA sequencing analysis, revealing novel compartments associated with individual Timp expression. These analyses provide evidence of the biological significance of Timp expression in the identified cell sub-types, which are consistent with novel roles in normal tissue homeostasis and changing roles in disease progression. This understanding of the tissues, specific cell types and microenvironment conditions in which Timp genes are expressed adds important physiological context to the growing array of novel functions for TIMP proteins.

## Introduction

Tissue inhibitors of metalloproteinases (TIMPs/Timps) are named so due to their principally ascribed function as inhibitors of extracellular matrix proteolytic activity, namely Metzincin proteases of the Matrix Metalloproteinase (MMP), A Disintegrin and Metalloproteinase (ADAM) and ADAM with thrombospondin motifs (ADAMS-TS) families. The Metzincins are major regulators of extracellular matrix (ECM) structure and composition [1]. As a result, TIMPs are routinely designated solely as tissue inhibitors of metalloproteinases. Although their metalloproteinase inhibitory functions in many disease states are well recognized, TIMPs are also widely expressed in non-diseased tissues with little detectable Metzincin protein expression and/or activity [2–4]. TIMPs display a broad targeting of MMPs, excluding TIMP1 which shows poor affinity for the six membrane-associated MMPs [4]. Inhibition of ADAM and ADAM-TS proteinases is more selective between the TIMP family members, with TIMP3 displaying the broadest inhibitory capabilities in this regard [5, 6]. Since their original discovery additional functions have been identified in TIMPs that are independent of their metalloproteinase-inhibitory activities, including direct agonism/antagonism of cell-surface receptors and interactions with matrisome partners [7, 8]. Although it has been over two decades since all four members of the TIMP family were identified, there remains to be an in-depth analysis of TIMP gene expression in organs, tissues, and cells of normal adult mammals. An understanding of the tissues and cell types that express TIMPs is important to add context to the ever broadening and increasingly complex functions of TIMPs that are all too often dismissed as non-canonical.

The advent of single cell RNA (scRNA) sequencing and spatial transcriptomics have greatly expanded researcher’s abilities to profile gene expression at the cellular level in various tissues. This was perfectly illustrated by the Tabula Muris Consortium’s breakout manuscript presenting whole organism scRNA sequencing data in mice [9]. We have utilized this rich source of data, analyzing approximately 100,000 murine cells, isolated by fluorescence-activated cell sorting (FACS) or microfluidic partitioning (droplet-based), for the expression of the Timp family of genes. We describe the diversity in Timp gene expression across nineteen murine tissues and seventy-three annotated cell types, providing new insights into the proportion of expressing cells and the degree of expression in specific cell populations. We identify several discrete cell types, particularly of mesenchymal and endothelial origins, with unique gene expression profiles that correlate with distinct expression of specific Timp family genes. In addition, we performed RNA ISH across 4 normal tissues with high/distinct Timp expression patterns to reveal unique regional expression patterns of each Timp family member.

Our detailed analysis of the tissues, cell-types and microenvironmental conditions in which Timps are expressed, adds significant physiological context, and enhances our understanding of both protease inhibitor activity and metalloproteinase-independent functions of TIMP proteins in their resident tissues.

## Results

### Tissue expression of Timp family genes in murine organ types

We first set out to describe Timp gene expression across the 18 organs analyzed in the Tabula Muris dataset in a simple format. The original Tabula Muris publication describes 20 organs, however we combined “brain myeloid” with “brain non-myeloid” (brain), and “diaphragm” with “limb muscle” (skeletal muscle). Normalized expression of the 97,420 analyzed cells were subject to a low count cut-off expression value of 0.5, selected to ensure that at least 80% of the cells with detectable expression of Timps1-4 were included. Cells above the cut-off expression levels were marked as positive. Additionally, of these positive cells, the mean expression levels were calculated. These data were incorporated into a circle packing graph describing Timp family expression profiles in 18 whole organs (Figure 1A). Our analysis of all four Timp family members demonstrates Timp2 expression is widespread, expressed in all 18 organs studied, and detected in 46% of the analyzed cells. Timp3 is also well represented as it is expressed in 35 % of cells and detected in 16 of the 18 analyzed organs. Timp1 and Timp4 display restricted expression across the analyzed organs, present above threshold in only 12% and 5% of total analyzed cells, respectively (Figure 1A). Mean Timp expression values in organs, calculated by excluding cells below threshold expression, generally correlate with the proportion of Timp expressing cells. The complete tabulated data sets describing Figure 1A can be found in Table S1.

**Figure 1.**
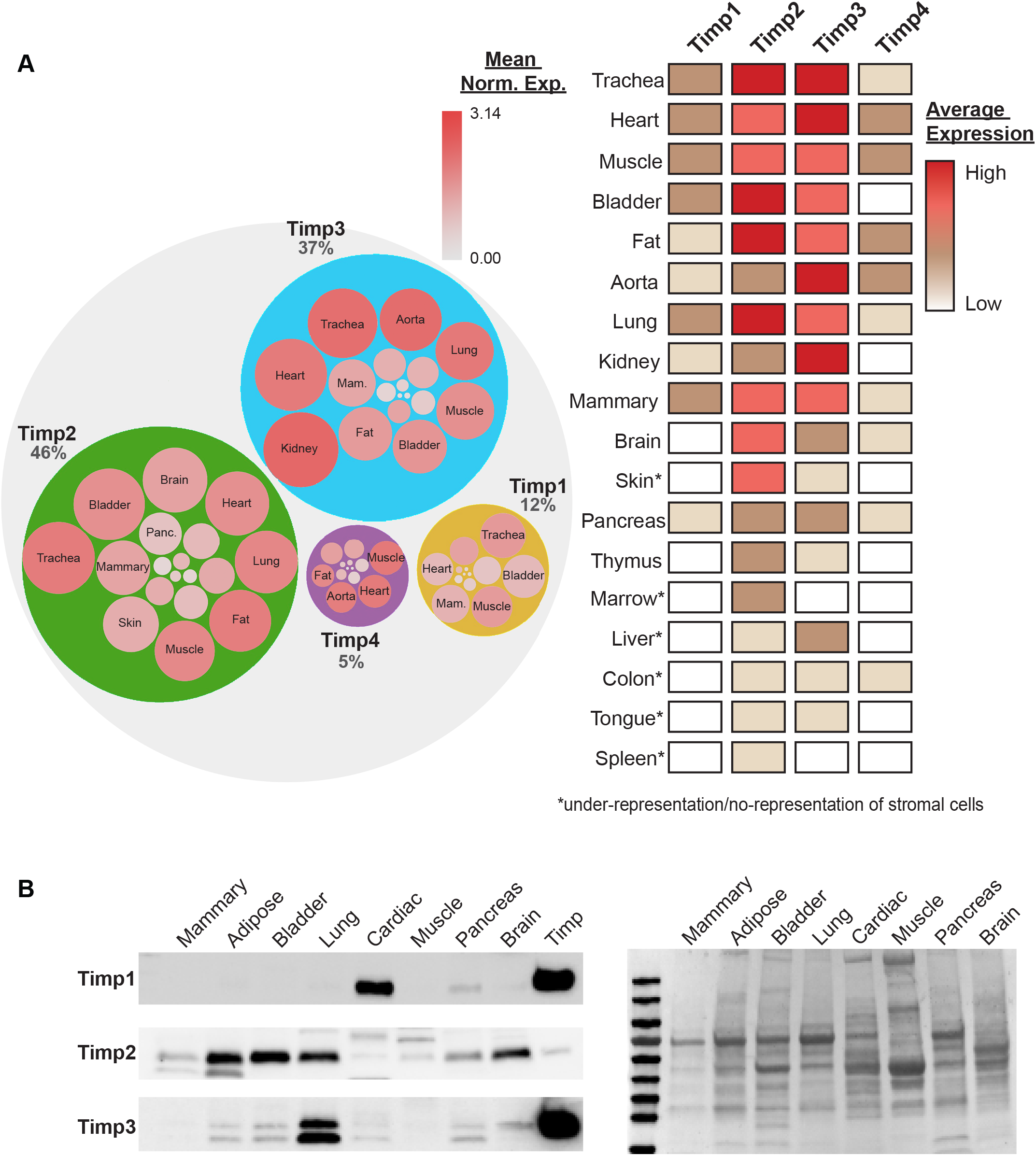
Comparison of Timp gene expression in healthy murine tissues. **(A)** Circle packing graph describing Timp gene expression in murine tissues. Indicated % values describe the total percentage of analyzed cells that are positive for Timp expression. Circle size is equivalent to percentage positive expression (positive expression = a normalized count above 0.5). Internal circles are also equivalent to percentage positive expression, with color describing the mean expression of positive cells (non-expressing cells are ignored in the mean expression calculation). **(B)** Immunoblotting of murine tissues plus 4ng (TIMP1/2) or 10ng (TIMP3) recombinant human TIMP1/2/3 describing Timp levels across various female C57/BL6 murine tissues. Ponceau red total protein staining is included as a reference. Abbreviations; Mam. (Mammary), Panc. (Pancreas).

It is important to appreciate that gene expression often poorly correlates with protein levels, which is a consequence of numerous levels of regulation that follow transcription (e.g., post transcriptional regulation by miRNAs, post-translational modifications). However, a recent study from the Genotype-Tissue Expression (GTEx) consortium reported that TIMPs 1-3 displayed good correlation, with Spearman’s correlation coefficients of between 0.68-0.7, all within the upper-quartile of over 12,000 genes/proteins analyzed (Table S2) [10]. We corroborate the observed organ expression profiles by immunoblotting of normal organs obtained from C57BL/6 mice for Timps1-3. Antibodies were pre-validated to confirm lack of cross-reactivity between the Timp family members using recombinant proteins produced in-house [11]. Unfortunately, despite great effort, we could not identify a suitable, commercial Timp4 antibody that would reliably detect Timp4 in murine tissues without excessive cross-reactivity with other Timps and/or unknown targets. Immunoblotting of murine tissues supports the findings that Timp2 is abundant across all tested samples (Figure 1B; full blots Figure S1). Timp1 protein expression is largely restricted in healthy organs, only detected at high amounts in cardiac tissue (Figure 1B; full blots Figure S1). The data in figure 1A describe significant mRNA levels of Timp3 found in many tissues, yet protein detection by immunoblotting is generally poor (Figure 1B). This could be attributed to poor correlation between mRNA levels and protein expression for Timp3, although data on human TIMPs challenges this idea [10]. The discrepancies between the transcriptome data and the observed protein expression levels may also be the result of Timp specific effects. For example, The Blood Atlas (from The Human Protein Atlas; https://www.proteinatlas.org/) quantified human TIMP1 protein levels in plasma at a concentration of 750ug/L (by mass spectrometry), which is almost an order of magnitude higher than that of TIMP2 in the circulation (83ug/L) [12]. It is unclear whether the same is true for circulating Timp1 concentrations in murine blood/plasma, but it may contribute to the high level of Timp1 protein observed in cardiac tissues. In contrast, the poor correlation between mRNA levels and protein expression for Timp3 may be attributable to significant interaction with glycosaminoglycans in the ECM, as described in several reports [13]. Tissue homogenization can generate a significant insoluble pellet that is discarded in processing. Thus, target loss during lysate homogenization and clearing may be exaggerated with Timp3 in the absence of chaotropic agents. We performed extensive testing with a range of Timp antibodies, in many cases observing a widespread lack of specificity in peptide detection. The poor performance of Timp antibodies in whole tissue lysates may also contribute to discrepancies between the RNA and protein levels that we observed. This suggest that the validation of new reagents for detecting Timp proteins in additional studies is necessary.

### Cell-type expression of Timp genes

After identifying these organ-specific patterns of Timp expression, we then characterized the specific cell-types expressing each Timp family member within each organ. For this, we retained the Tabula Muris Consortium’s original cell type designations (cell_ontology_class), as shown in Figure S2 & S3. However, to simplify our data presentation, we first reduced the Tabula Muris designations into a set of broader cell type designations based on the original cell_ontology_classification, which we refer to as CellTypeB subclassification (Table S3, Figure 2A). Subsequently, we arranged these CellTypeB subclasses into one of seven general cell classifications, which we list as CellTypeA (epithelial, hematopoietic, stromal, neuronal, myocytes, adipocytes and unknown) (Table S3 and Figure 2B). This results in a more manageable presentation of cell-type expression profiles within specific organs/tissues, with Figure 2B focusing on the 10 tissues with highest Timp expression.

**Figure 2.**
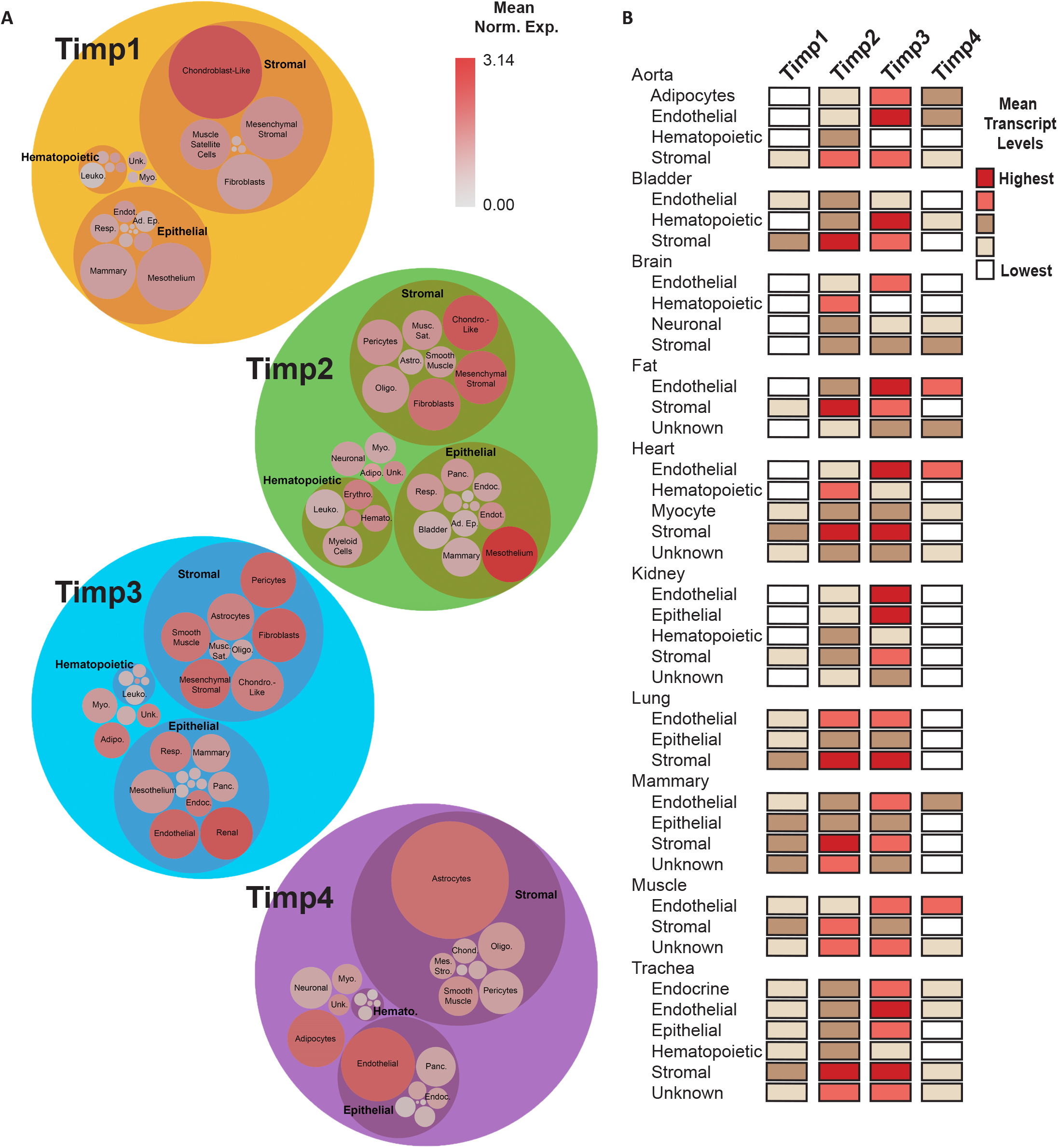
Cell-type expression of Timp genes. **(A)** Circle packing graphs describing Timp gene expression in murine cell-types. Cell-types are loosely grouped into epithelial, stromal, hematopoietic, and un-assigned groups, as described in the text. Circle size is equivalent to percentage positive expression (positive expression = a normalized count above 0.5). Internal circles are also equivalent to percentage positive expression, with color describing the mean expression of positive cells (non-expressing cells are ignored in the mean expression calculation). **(B)** Visualization depicting the broad cell-type specific expression of Timp family members from cells analyzed in the Tabula Muris dataset. Abbreviations; Unk. (Unknown), Myo. (Cardiomyocytes), Endot. (Endothelium), Ad. Ep. (Adipose Epithelium), Resp. (Respiratory Epithelium), Musc. Sat. (Muscle Satellite Cells), Astro. (Astrocytes), Oligo. (Oligodendrocytes), Adipo. (Epicardial adipocytes), Erythro. (Erythrocytes), Leuko. (Leukocytes), Hemato. (Hematopoietic), Panc. (Pancreatic Epithelium), Endoc. (Endocrine), Chondro-like/Chond. (Chondroblast like-cells), Mes. Stro. (Mesenchymal Stromal).

Timp1, often characterized as an “inducible” member of the Timp family [3], displays a restricted cell-type expression profile in normal organs that is consistent with this designation (Figure 1 & 2). Only chondroblast-like cells, identified in skeletal muscle, maintain high levels of Timp1 expression (Figures 2 & S2). Analysis of Timp4 expression, although low in most organs, identifies discrete cluster-specific expression in endothelial cells across several tissues (Figure 2). In addition, Timp4 transcripts can be found at appreciable levels in aorta-associated adipocytes and astrocytes of the brain, particularly Bergmann glial cells (an astrocyte sub-type found in the cerebellum) (Figures 2 & S2). Timp2 has an expansive expression profile across all organ types (Figure 1). This is also evident in the cell-type analysis, with broad expression of Timp2 identified across all tissue types in cells of mesodermal lineages (Figure 2). On the contrary, Timp3 displays highly enriched expression in stromal, endothelial, and other vascular-associated cell types, such as aorta-associated adipocytes, brain pericytes and cardiac fibroblasts (Figures 2, S2 & S3). In general, Timp expression is not associated with cells of epithelial origin, although the highest expressing cell types for Timp2 and Timp3 are mesothelial cells (of the plural cavity, identified from lung tissue samples) and renal tubule cells, respectively, which are both included as epithelia in the original Tabular Muris scheme. However, it is important to note that only 24 mesothelial cells were identified in the Tabula Muris dataset, which may not reflect sufficient sampling of this cell population. A detailed tabulation of cell type annotations and their Timp expression can be found in Tables S4 and S5.

### Mammary gland Timp expression

As is common with most organs, only Timp1-3 are significantly expressed in murine mammary tissue, with little or no detection of Timp4. Also, consistent with other organs, stromal and endothelial cells are the predominant sources of Timp expression within the mammary tissues, although basal and luminal epithelial cells also contribute (Figures 3A & 3B). Closer inspection of stromal cells alone reveals multiple superclusters (designated Superclusters 1, 2 & 3) associated with unique expression of Timps1-3 (Figures 3C & S2). Supercluster 1 is associated with low expression of all three Timps, and differential expression (DE) analysis of which reveals an increase in oxidative phosphorylation and a significant drop in ECM remodeling. This finding of diminished ECM remodeling is evidenced by an overwhelming downregulation of matrisome genes (Figures 3C, 3D, Table S6). Although pathway analysis does not describe any obvious inflammatory role for Timp low mammary stromal cells (supercluster 1), DE identifies a strong up-regulation in the proinflammatory cytokine/chemokine Il1b (+1.9 log2 FC) and Ccl5 (+7.5 log2 FC) (Table S6), which may allude to a potential immune regulatory role. Supercluster 2 is characterized by high Timp2/3 expression, with pathway analysis describing an expression profile associated with acute phase inflammation (14 gene hits, giving a positive activation score: Table S7). Manual assessment of the DE gene list highlights that this cluster is Pi16+, Ly6c1+ and Dpp4+, a group of fibroblast progenitor markers [14]. Supercluster 3 displays a notable decrease in Timp2/3 levels and, inversely, an increase in Timp1 expression. The increase in Timp1 coincides with an interesting increase in Cd63, a well-recognized Timp1 receptor. Although pathway analysis suggests that this cluster has a limited involvement in acute phase inflammation, there is an increase in Mmp3 and Mmp19 levels that are suggestive of a role in the modulating inflammation [15] (Table S8).

**Figure 3.**
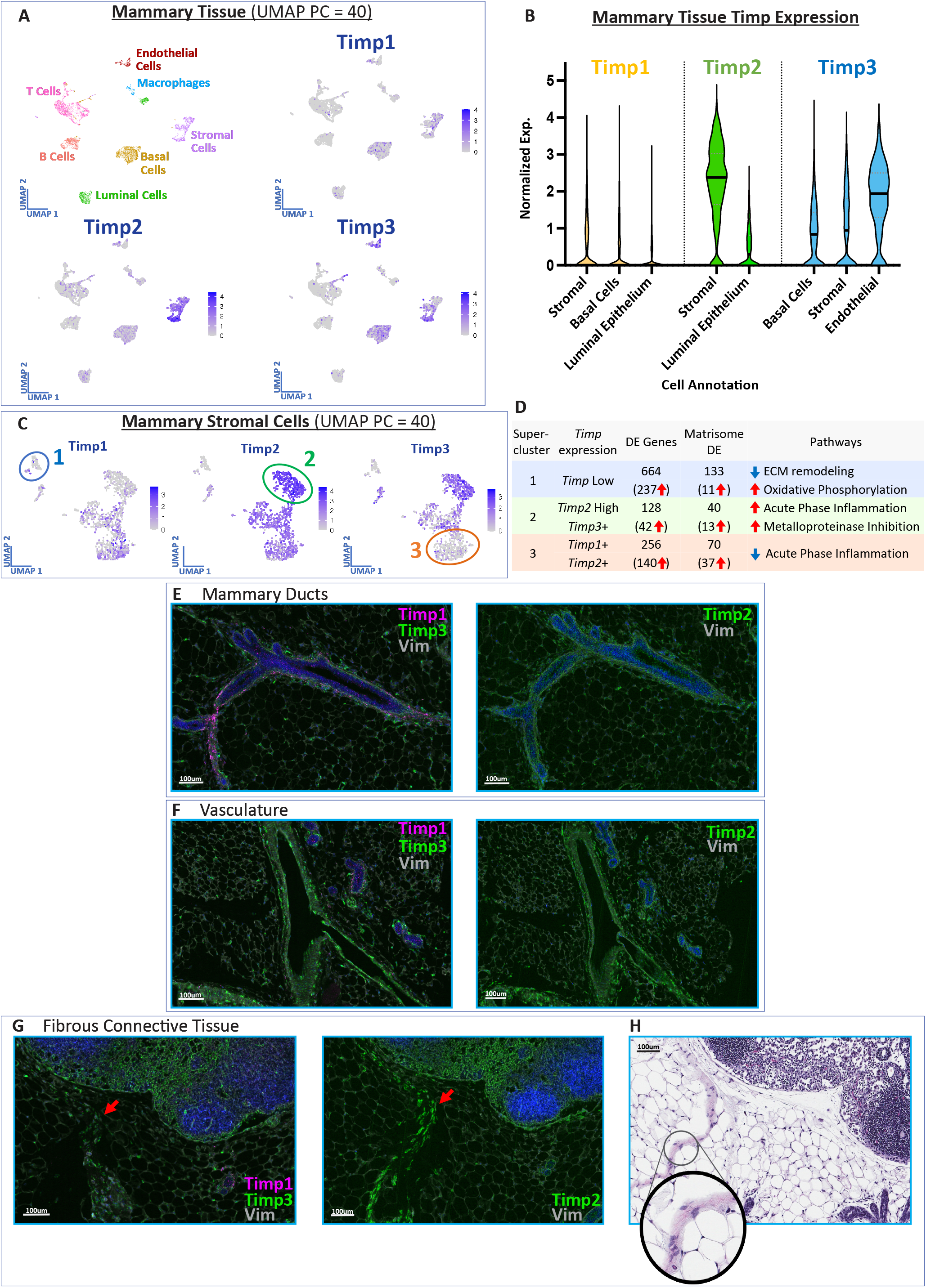
Mammary tissue expression of Timp family members. **(A)** UMAP clustering of murine mammary tissue transcriptomes illustrating the tissue-wide expression patterns of Timps1-3. **(B)** Violin plots describing several key cell-types with appreciable Timp expression levels in mammary tissue. **(C)** Visual interpretation of stromal cell UMAPs for Timp1-3 in mammary tissue, with 3 highlighted superclusters. **(D)** Comparison of the differentially expressed (DE) genes, DE matrisome genes, and altered pathways across the 3 identified superclusters. DE gene lists are filtered with adjusted p-values < 0.05 and a minimum 2-fold change in expression. RNA in-situ hybridization (RNA ISH) and immunohistochemistry reveals the spatial distribution of Timp transcription and vimentin (Vim) expression across the **(E)** ducts/acini, **(F)** vasculature and **(G)** fibrous connective tissue (red arrows). DAPI staining is represented as blue nuclei and vimentin staining is grey scaled for a high contrast image. **(H)** H+E image of an adjacent section to the fibrous connective tissue RNA ISH images, revealing a fibrous mesh-like collagenous structure.

RNA ISH for Timp1/2/3 was performed in C57BL6/J mammary tissues to corroborate the scRNA sequencing data from Tabula Muris. Timp3, and to a lesser extent Timp2, can be found throughout the mammary parenchyma, likely representing mammary adipocytes, endothelial cells, and various immune cell subtypes. Timp3 is also noticeably expressed throughout periductal regions of the mammary gland (Figure 3E). In more limited and highly localized periductal regions, Timp1 expression is readily detected (Figure 3E). In addition, low levels of Timp1 & 2 are noted in the luminal epithelium of ducts and acini (Figure 3E). Our analysis of the Tabula Muris data demonstrates thatTimp3 is abundantly expressed in endothelial cells and RNA ISH supports these conclusions, illustrating that Timp3 is expressed in all regions across the vasculature (Figure 3F). Timp2 expression is also detected in the perivascular region (Figure 3F). The strongest Timp2 signal arises within bands of fibrous connective tissue observed throughout the mammary tissue (Figures 3G & S4), which are clearly visualized with H+E staining (Figure 3H). It is also noted that Timp2 & 3 expression permeates through the non-hematopoietic, lymphatic endothelial cells of the lymph nodes (Figure 3G). A surprising additional observation from the RNA ISH experiments is that Timp2 and Timp3 were clearly and uniformly expressed in the peripheral nerve tissue (Figure S5). Closer inspection of the RNA ISH data suggests that Timp3 positive cells are contained within the peripheral nerve sheath and may represent either neuron or Schwann cell expression. In contrast, Timp2 positive cells are located within the peripheral nerve sheath and surrounding connective tissue.

### Timp expression in Pulmonary tissues

As observed in mammary tissue, Timp4 is not found within the lung. Timp2 & 3 are the predominant Timps expressed in pulmonary tissue (Figures 4A & 4B), both of which display a strong correlation in expression across the whole tissue (Pearson correlation coefficient = 0.64; 6928 cells total) and expressed across multiple cell types. The principal source of Timp expression in the lung is found in stromal cells, and an in-depth analysis of stromal Timp expression patterns reveal 4 major superclusters, one of which is Timp High (Figure 4C; green. circles) and another Timp low (Figure 4C; orange circles). Comparable to Timp high stromal cells in mammary tissue, the Timp high pulmonary stromal cells are Pi16+ Ly6c1+, and pathway analysis highlights acute phase inflammation as a key discriminatory pathway (10 gene hits, giving a positive activation score: Table S9). Additionally, pathway analysis suggests that these Timp high pulmonary stromal cells display enhanced extracellular matrix deposition and remodeling (Figure 4D). In contrast, pathway analysis of the Timp low stromal cluster reveals a range of activation scores across various pathways, notably reduction in acute phase inflammation and ephrin receptor signaling, along with an increase in cell stress signaling via the Eif2 pathway (Figure 4D & Table S10). Inspection of the gene changes and pathway analysis also reveals a unique expression pattern of ECM genes, suggesting that these cells display unique remodeling capabilities associated with high expression of collagen IV & VIII, osteopontin, periostin and vitronectin (Table S10).

**Figure 4.**
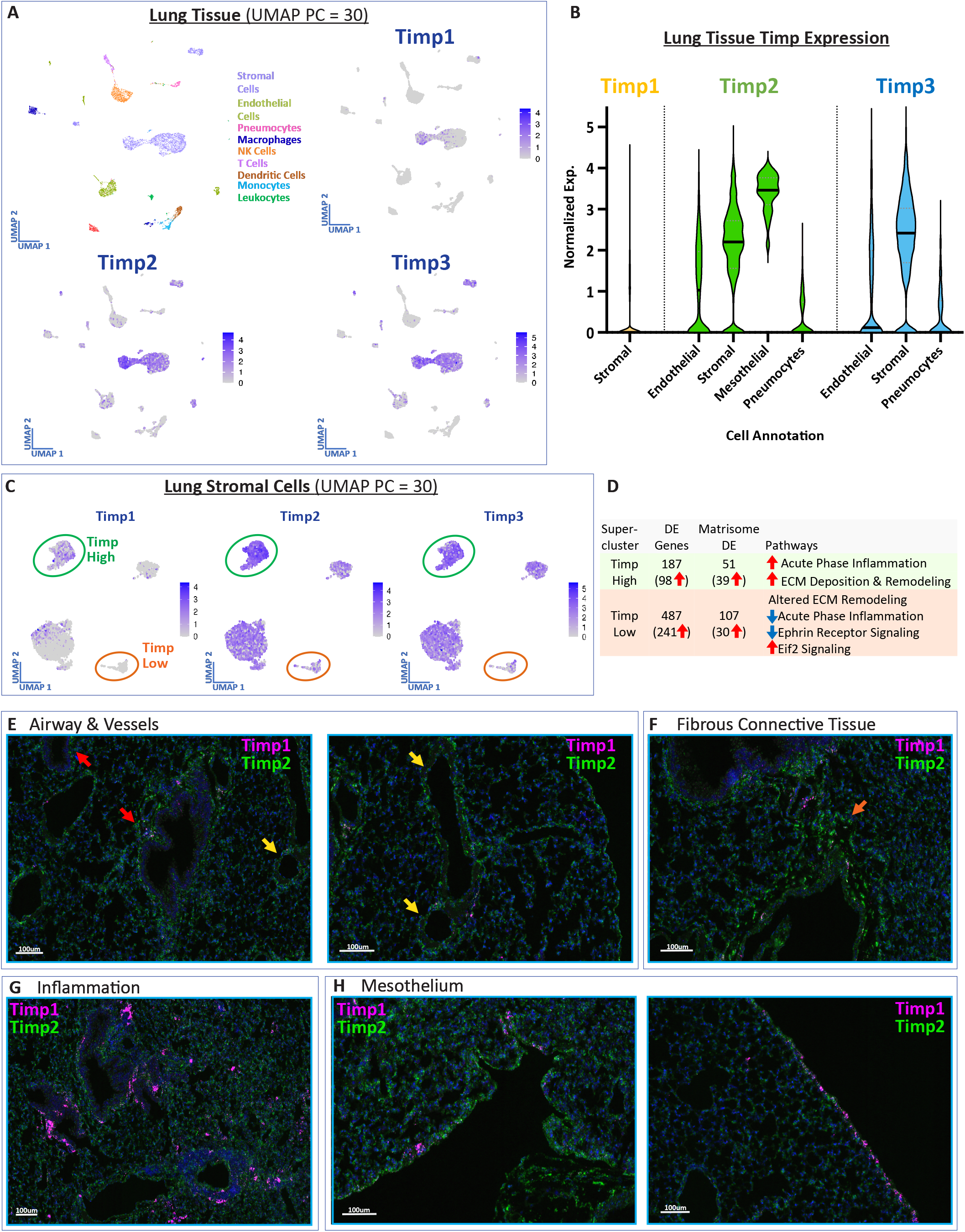
Lung tissue expression of Timp family members. **(A)** UMAP clustering of murine lung tissue transcriptomes illustrating the tissue-wide expression patterns of Timps1-3. **(B)** Violin plots describing several key cell-types with appreciable Timp expression levels in lung tissue. **(C)** Visual interpretation of stromal cell UMAPs for Timp1-3 in lung tissue, with 2 highlighted superclusters. **(D)** Comparison of the differentially expressed (DE) genes, DE matrisome genes, and altered pathways across the identified Timp high and Timp low superclusters. DE gene lists are filtered with adjusted p-values < 0.05 and a minimum 2-fold change in expression. RNA in-situ hybridization (RNA ISH) reveals the spatial distribution of Timp transcription across the **(E)** airways (red arrows) and vessels (yellow arrows), **(F)** fibrous connective tissue, **(G)** regions of immune infiltration and expansion, and **(H)** the pleural mesothelium. DAPI staining is represented as blue nuclei.

Since there was a strong correlation between Timp2 & 3 expression in scRNA sequencing data, RNA ISH was performed using Timp1 & 2 (with the assumption that the strong correlation between Timp2 & 3 would prevail). Timp2 expression is distinctly universal in lung tissue, with enhanced expression illustrated in peribronchiolar/perivascular compartments and fibrous connective tissue throughout the lung (Figure 4E). The Tabula Muris data describes how Timp1 expression is minor within the lung, however histological analysis of the C57BL6/J lung sections reveal focal areas of atelectasis with chronic inflammation, characterized by infiltration of macrophages and clusters of mononuclear cells within the alveolar space, as well as hyaline membrane formation (Figure S6). This focal region of chronic (resolving) inflammation reveals a dense population of Timp1+ cells, consistent with macrophage/monocyte infiltration. Additionally, we observed punctate cells subjacent to the pleural surface that are strongly positive for Timp1 expression (Figure 4H). These may again reflect tissue macrophage surveillance of the mesothelial surface.

### Cardiac Timp expression

Cardiac tissues are differentiated from mammary and pulmonary tissues by significant transcriptional activity for all four TIMPs (Figures 1A, 1B, & 1C). Stromal cell types are the dominant suppliers of Timp2 & 3, although these cells do not display any cluster-specific patterns of expression (Figures 5B, 5C, & S3). Endothelial cells also represent a major source of cardiac Timp expression, representing one of the few sources of detectable Timp4 expression. Unlike stromal cell populations in the mammary gland and lung, cardiac endothelial cells do not form distinct superclusters with vast differentially expressed gene-sets, but the defined clusters do display unique expression of Timp4 (Figure 5D). The Timp4+ cluster displays only 10 genes with >2-fold changes in gene expression, one of which is Timp4. Removal of the fold-change threshold reveals 165 significant gene changes that are suggestive of decreases in hypoxia and eNos signaling pathways, as well as enhanced tight junction formation (Figure 5D & Table S11).

**Figure 5.**
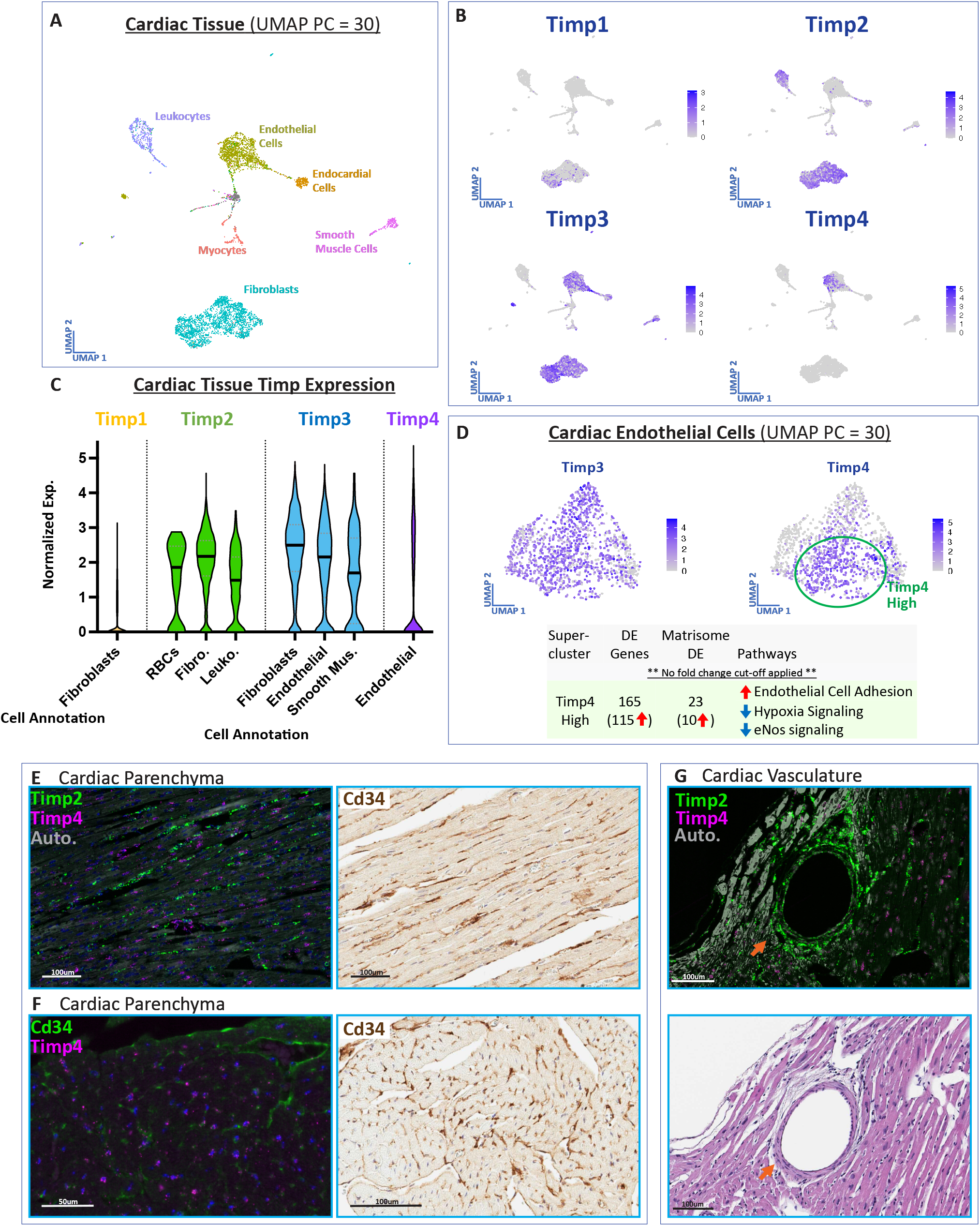
Cardiac tissue expression of Timp family members. **(A)** UMAP clustering of murine cardiac tissue transcriptomes illustrating and **(B)** the tissue-wide expression patterns of Timps1-3. **(C)**. Violin plots describing several key cell-types with appreciable Timp expression levels in cardiac tissue. **(D)** Visual interpretation of endothelial cell UMAPs for Timp3 and Timp4 in heart tissue, with the highlighted Timp4+ supercluster and comparison of the differentially expressed (DE) genes, DE matrisome genes, and altered pathways across the identified Timp high and Timp low superclusters. DE gene lists are filtered with adjusted p-values < 0.05, with no fold change threshold applied. **(E)** RNA in-situ hybridization (RNA ISH) for Timp2 (Cy3 – Green) and Timp4 (Cy5 – Magenta) reveals the spatial distribution of Timp2/4 expression across the cardiac parenchyma. Grey-scaled regions of autofluorescence (FITC channel) are indicated. Cd34 expression was also assessed via IHC in an adjacent tissue section. **(F)** Due to high autofluorescence in the first animal, a new heart was collected for RNA ISH and IHC for the assessment of Timp4 RNA (Cy5 – Magenta) and Cd34 protein expression (Alexa Fluor 488 – Green), with an analogous region analyzed by IHC for Cd34 expression. **(G)** RNA ISH for Timp2 and Timp4 in cardiac vasculature, with a corresponding region analyzed via hematoxylin and eosin staining. In fluorescence images, DAPI staining is represented as blue nuclei.

To determine if there are localized regions of Timp4+ endothelial cells that could explain their subtle transcriptomic differences, we performed RNA ISH to assess Timp4 transcripts in cardiac tissue. Timp4 expression is found solely in the Cd34+ cells that localize in parallel with cardiac myocytes and are distinct from Timp2 expressing cells within the same region (Figure 5E/F). Visual comparison of Timp2 and Timp4+ cells in parallel formation to cardiac myocytes reveals no correlation in expression, with the Timp2+ foci likely representing cardiac fibroblasts (Figure 5C). In contrast with the Tabula Muris data, RNA ISH reveals visible TIMP2 expression within the coronary arteries, with notable expression in both the endothelial and perivascular cells (pericytes) clearly separated by the subendothelial basement membrane (Figures 5F, S7).

### Skeletal muscle Timp Expression

Like cardiac muscle, skeletal muscle of the limb exhibits expression of all four members of the Timp family across a range of cell types including endothelia, various stromal cells, and macrophages (Figure 6A, 6B, & 6C). Single cell analysis also included a significant proportion of chondrocyte-like cells, described by the Tabula Muris Consortium as cells that express the chondrocyte markers Chodl and Acta2, which may represent a subset of muscle-resident mesenchymal cells with chondrogenic potential [9]. These cells are rich in Timp1, 2, & 3 expression, and present as a uniform population (Figure S2). Similar to cardiac endothelial cells, skeletal muscle endothelia are universally Timp3+. Timp4+ endothelia of skeletal muscle represent a large proportion of cells that, unlike cardiac endothelia, form a distinct supercluster (Figure 6D; green circle). DE identified 146 gene changes in Timp4+ skeletal muscle endothelia, 32 of which are matrisome genes that includes an increase in the basement membrane associated Collagen IV (Figure 6D, Table S12). Pathway analysis of this gene set highlights a likely reduction in hypoxia and eNos signaling, analogous to the Timp4+ population of cardiac endothelial cells. This Timp4+ cluster also displays a reduction acute phase response signaling factors such as Interleukin-6, von Willebrand factor and fibronectin (Figure 6D, Table S12). Interestingly, single cell profiling also reveals two separate clusters of Timp4-cells are Timp2+ (Figure 6D; blue and orange circles). Pathway analysis of the two Timp2+ clusters (versus the Timp4+/Timp2-cluster) reveals a number of significant genes changes associated with enhanced fibrotic responses and hypoxic signaling in supercluster 1, or an increase in acute phase inflammation associated with interleukin-6 signaling in supercluster 3 (Figure 6D, Tables S13/S14).

**Figure 6.**
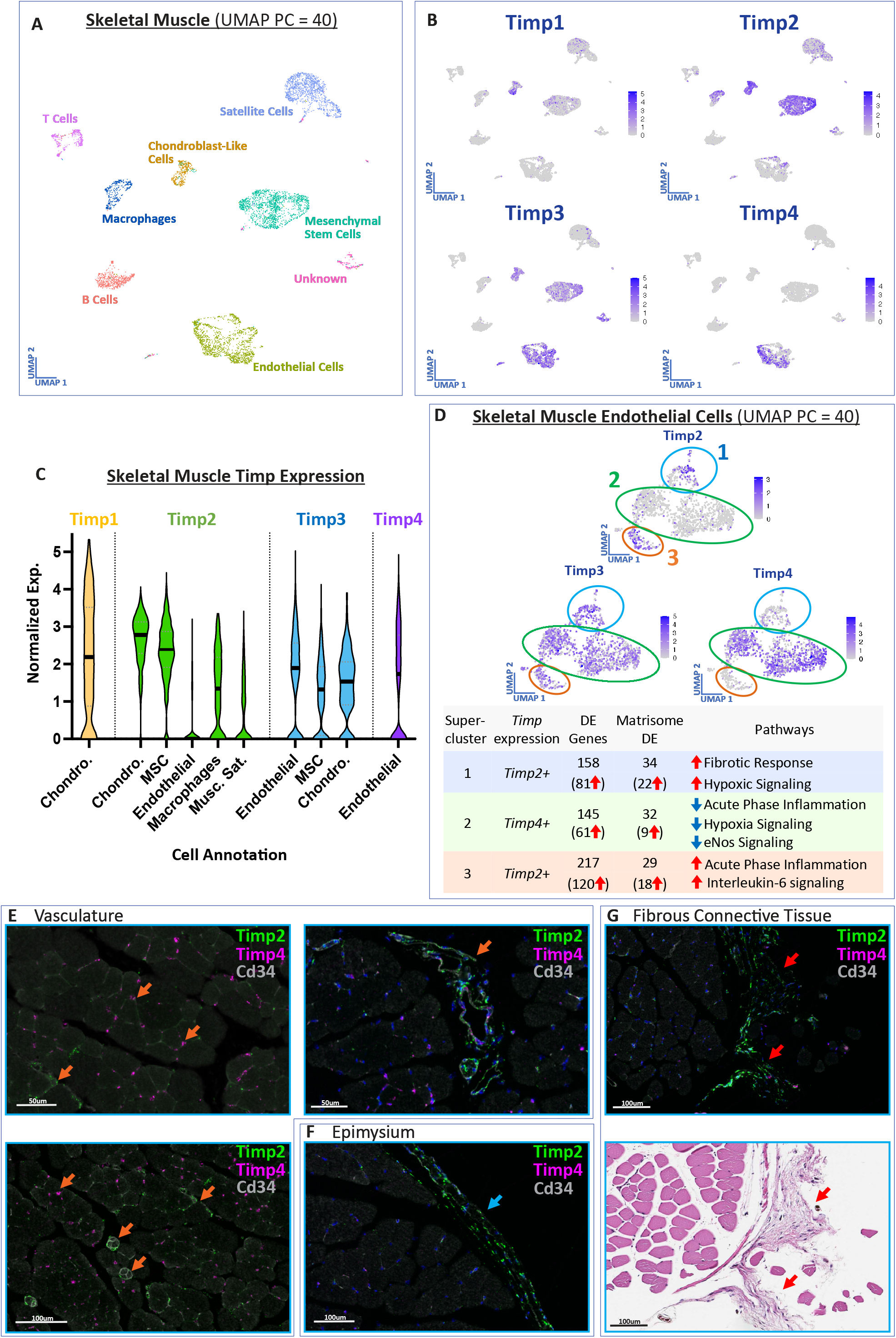
Skeletal muscle tissue expression of Timp family members. **(A)** UMAP clustering of murine skeletal muscle tissue transcriptomes illustrating and **(B)** the tissue-wide expression patterns of Timps1-3. **(C)**. Violin plots describing several key cell-types with appreciable Timp expression levels in skeletal muscle tissue. **(C)** Visual interpretation of endothelial cell UMAPs for Timp2-4 in skeletal muscle, with 3 highlighted superclusters and comparison of the differentially expressed (DE) genes, DE matrisome genes, and altered pathways across the 3 identified superclusters. DE gene lists are filtered with adjusted p-values < 0.05 and a minimum 2-fold change in expression. RNA in-situ hybridization (RNA ISH) and immunohistochemistry reveals the spatial distribution of Timp2 (Cy3 – Green) and Timp4 (Cy5 – Magenta) transcription and Cd34 expression across the **(E)** vasculature (orange arrows), **(F)** epimysium (blue arrow) and **(G)** fibrous connective tissue (red arrows). DAPI staining is represented as blue nuclei and Cd34 staining is grey scaled for a high contrast image. **(G)** H+E image of an adjacent section to the fibrous connective tissue RNA ISH images, revealing a fibrous mesh-like collagenous structure.

Comparison of RNA-ISH signals for Timp2 and Timp4 reveals that, like the Timp4+ endothelial cells of cardiac tissue, Timp4+ cells of skeletal muscle constitute a distinct endothelial population that is closely associated with the endomysium that surrounds skeletal muscle fibers (Figure 6E). In addition, occasional cells within the endomysium display limited Timp2 transcript expression that may represent one of the Timp2+ clusters identified in the scRNA sequencing analysis but may also be identified as Timp2+ muscle satellite cells (Figure 6C). A definitive Timp2+ population of endothelial cells is represented within the intermediate- and large-sized vasculature of perimysium, with some larger vessels also exhibiting Timp4 transcripts (Figure 6E). Strong Timp2 expression was observed within the epimysium and fibrous connective tissue surrounding the bulk of the muscle, which may be constitute mesenchymal stem cells or chondroblast-like cells identified in the Tabula Muris data (Figures 6F & 6G).

## Discussion

Despite their key functional attributes and expanding biological roles, Timps are an understudied family of endogenous proteins. In this study, we describe Timp gene expression across eighteen different organ systems utilizing the publicly available Tabula Muris scRNA sequencing dataset of C57BL/6 mice. We then focus our attention on four tissues that display high levels of Timp expression, as well as unique cluster-specific patterns of Timp expression. Timp1 and 3 have been referred to as inducible Timps [3]. We show that Timp1 expression, although present, is low in many normal tissues, supporting the idea that Timp1 is generally inducible in response to tissue damage or inflammation. Despite the broadly low Timp1 expression, there appears to be distinct, dense populations on Timp1 expressing cells within the periductal regions of mammary tissue, and within the mesothelium and parenchyma of the lung (Figures 3 & 4). These cells may represent immune surveillance cells, such as tissue resident monocytes or macrophages, that would be consistent with the limited Timp1 expression in bone marrow monocyte and stem cell population (Figure S2). In addition, there are numerous reports describing an increase in Timp1 expression in response to local tissue inflammation associated with pathologies such as fibrosis and cancer [16–20]. Further supporting the generally accepted association between Timp1 and inflammation, the lungs used in our RNA ISH analysis displayed regions of chronic inflammation that corresponded with a distinct increase in Timp1 expressing cells (Figures 4G & S6). Comparison of the differential expression (DE) profiles of Timp1+ stromal cells in breast and lung tissue reveal a list of 98 genes that display similar DE (without a fold change threshold). Included in this list are the Timp1 receptor Cd63 [18], and the pro-inflammatory metalloproteinase MMP3 [15, 21] (Table S15). The correlation between Timp1 and Cd63 expression may function to mediate autocrine cell survival/proliferation under inflammatory conditions.

In contradistinction to the restricted, inducible expression of Timp1, Timp2 is almost universally expressed in stromal cells, and is often correlated with that of Timp3. Timp2 & 3 high stromal cells exhibit specific patterns of expression, displaying a shared upregulation of markers associated with a universal fibroblast subtype that presents as a Pi16+ population (Table S16). Additional markers associated with Timp2+/Timp3+/Pi16+ fibroblasts include Ly6c1, Col14a1, Scara5 and progenitor-associated genes Cd34 and Ly6a/SCA1 [22, 23], suggesting that these populations of fibroblasts may represent a pool of undifferentiated adventitial fibroblasts with broad lineage potential, as described by Buechler et. al. [14, 24–26]. RNA ISH reveals that Timp2 expression is generally strongest within fibrous connective tissue, organ boundaries (lung pleura, skeletal muscle epimysium) and perivascular regions, implying that the proposed fibroblastic progenitor cells may reside within these domains in tissues. However, it remains to be determined if there is a direct functional link between Timps and the aforementioned stem cell-like characteristics. Interestingly, the side population in six NSCLC cell lines, which includes the cancer stem cell population, was inversely correlated with their Timp2 expression [27]. Further investigation of the role of Timps in regulation of tissue stem cell behavior are warranted.

The stroma is a broad term that is sometimes used to encapsulate all non-epithelial cells in tissues. In more specific-terms, stroma (or stromal cells) is used to represent cells of mesodermal lineages. In the case of the Tabula Muris dataset, this includes fibroblasts, mesenchymal cells, smooth muscle cells, muscle satellite cells, chondroblast-like cells, pericytes, oligodendrocytes (and precursors) and astrocytes. One of the richest sources of Timps can be identified in adipose mesenchymal stem cells (aMSCs) (Figure S2). These multipotent stromal cells cluster into numerous visibly distinct populations/sub-types. We observed that aMSCs are universally high in Timp2 expression, followed by somewhat lower but diffuse expression of Timp3 and, to a lesser extent, Timp1 (Figure S2). Similarly, the brain is another tissue of notable Timp expression that presents with no clearly defined cluster-specific expression patterns, an observation that also extends to skeletal muscle mesenchymal stem cells (Figures S2 & S3). For this reason, Timp expression was not assessed through RNA ISH in these tissues.

Within the stromal cells analyzed in mammary and lung tissue, there were clusters of cells with low/reduced Timp expression. Between the Timp low stromal cells of breast and lung tissue, there were 115 shared DE genes (with a fold change cut-off of 2, Table S17), 54 of which were downregulation in matrisome genes. This finding implies that these Timp low stromal cells have a limited role in the regulation of ECM composition and function. It is uncertain what the Timp low stromal populations represent within tissues, but they may represent a terminally differentiated stromal population with reduced matrix remodeling capabilities, as suggested by the downregulation of matrisome genes and fibroblastic progenitor markers Pi16 and Cd34.

Despite Timp2’s strong association with stromal cell types, its highest expression was identified in mesothelial cells from lung tissue. Mesothelial cells are descriptively referred to as a simple squamous epithelial lining of the pleural and abdominal cavities. However, they are derived from the embryonic mesoderm and have been ascribed a wide variety of functions, including fluid transport, ECM modulation, as well as cytokine and growth factor secretion [28]. There have been limited reports of Timp2 expression in the mesothelium [29], and the Tabula Muris dataset only annotated 24 mesothelial cells in their lung tissue. RNA ISH analysis supports the limited findings from RNA sequencing regarding mesothelial Timp2 expression. RNA ISH also revealed dense, localized regions of Timp1 expression within the lung pleura that may be associated with local inflammatory responses or immune surveillance. Given their mesodermal origin it is not surprising that mesothelial cells show significant similarities with stromal/mesenchymal cells with regards to their Timp expression levels.

At a transcriptional level Timp3 is highly expressed in many normal tissues, particularly within the vascular and renal systems [30, 31]. Timp3 expression is strongly associated with the vasculature and is widely expressed across all structural components of the cardiovascular system. Vascular and renal tubular cells are exposed to shear stress from the flow of fluid [32, 33]. Whether Timp3 expression in these cells is lineage dependent or a result of outside-in signaling due to luminal flow is an important question of interest, with shear stress having been shown to enhance Timp3 expression in pericytes [34]. A clear advantage for expression of Timp3 in these situations, over that of other Timp family members, is the fact that Timp3 is matrix-associated through specific interactions via glycosaminoglycans. This is partially supported by an N-glycosylation site near the C-terminus of Timp3 that helps anchor Timp3 to proteoglycans [13], thereby maintaining a presence of MMP-inhibitory capabilities that may not be maintained with readily diffusible Timps [5]. This would contribute to ECM stability in microenvironments that maintain active flux, such as those found in the vascular and renal systems.

Despite Timp3 being the dominant vascular associated Timp, Timp4 is co-expressed in subsets of endothelial cells from both cardiac and skeletal muscle tissue, with that latter also characterized by a Timp2+ population (Figure 5 & 6). In addition, Timp2 & 4 can be also detected in clusters of endothelial cells from adipose tissue and trachea (Figure S3). Timp2 and Timp4 display an inverse correlation of expression in skeletal muscle (Figure 6D; echoed within trachea and adipose endothelial cell populations, Figure S3), with skeletal muscle endothelia displaying a prominent inverse correlation that consists of two opposing Timp2+ populations that are separated by a Timp4+ population. Gene set pathway analysis strongly suggests that Ppar-γ signaling is responsible for Timp4 expression in Timp4+ cardiac, skeletal, adipose and trachea endothelial cells (Tables S18-S21, “Upstream Analysis”). Despite being poorly characterized, Timp4 has been shown to be induced through Ppar-γ signaling [35].

Why do distinct cell type clusters express different Timps, and is differential expression of Timp family members of real physiological consequence? In some cases, cluster specific Timp expression is associated with a host of transcript changes, including both matrisome and non-matrisome associated genes. In these instances, it is possible that there is collateral transcript expression due to overlap of broad and divergent signaling pathways. In other cases, cluster definition and significant transcript changes are minimal, as observed with Timp4+ cardiac endothelial cells. Interestingly, despite minor alterations in transcript expression, Timp4+ endothelial cells are predominantly exclusive to the smallest cardiac microvessels that are closely associated with myocardiocytes (Figure 5E). The utility of a more targeted increase in Timp expression is yet to be determined, as enhanced Timp protein levels can have multiple outcomes that include MMP-dependent and -independent biological activities [36].

We acknowledge that this analysis of the Tabula Muris dataset has its limitations. The techniques that are used to isolate and extract single cell RNA displays internal biases towards complicit cell types, leading to a misrepresentation of the cell type proportions. These limitations are exampled by the lack of skeletal muscle myocytes, no stromal cell representation in several tissues (colon, liver, skin, bone marrow), and the disproportionately low number of neurons. A distinct omission from the scRNA sequencing data is the lack of Timp2+ cardiac endothelial cells, which were prominently revealed using RNA ISH (Figure 5F). Ideally, data from parallel single cell RNA sequencing and bulk RNA sequencing experiments would be used to address these issues with cell type biases. Regardless, cell type proportions can vary between regions within individual organs, as exampled by the distinct chronic inflammatory cell infiltration within two of the minor lung lobes from the otherwise healthy murine lungs used in our RNA ISH experiment (Figure S6). The latter point can be addressed with concurrent RNA ISH analyses. An intriguing additional observation from our RNA ISH studies was the prominent identification of Timp3 expressing peripheral neurons within mammary tissue (Figure S5). The relationship between Timp3 and neuronal function is poorly understood, although it has been reported that Timp3 can promote Schwann cell myelination [37].

The GTEx consortium recently reported that, although transcript-protein level correlation is generally poor, human TIMPs 1-3 protein expression correlates well with transcript levels [10]. Although murine and human Timps/TIMPs display very high sequence identity, it is unknown whether this corresponds with a similar tissue/cell-type expression profile. At the transcriptional level, the heart is awash with Timp2, 3, & 4 expression. However, our immunoblot analysis of cardiac tissue revealed poor detection of Timp2 & 3, yet strong detection of Timp1 that may be represent contamination from circulating plasma Timp1 [12]. Despite these findings, the general poor performance of Timp antibodies with tissue samples, or sub-optimal tissue processing for Timp detection, warrants caution when interpreting immunoblot data, as evidenced by our extensive antibody testing.

We describe the tissues and cell-types that express members of the Timp family in murine tissues. Tissue expression of Timps is drastically altered in disease states [3, 38–40], and in situations where Timp expression is relatively unaffected many matrisome/matrisome-associated genes that can modulate Timp function are affected [41]. Understanding how these cell clusters and their matrisome gene expression patterns change in disease may reveal mechanistic insights that can be targeted for novel therapeutic interventions. Our study identifies unique Timp expressing cells that are principally of mesodermal origin and distributed across a broad range of healthy tissues. Considering the potential therapeutic utilities of Timp family genes, we propose that targeted expansion of these tissue-resident cell subtypes may represent viable treatment alternatives for a range of pathologic conditions. Furthermore, understanding the behavior of the Timp expressing cellular subtypes in response to pathologic conditions, such as cancer, neurodegenerative disease, and myocardial infarction, may reveal appropriate therapeutic conditions for utilizing/targeting Timp family members.

## Experimental Procedures

### Tissue collection and protein extraction for immunoblotting

Organs for immunoblotting were harvested from wild type female C57BL/6 mice (4-month-old) immediately following euthanasia by CO_2_ narcosis with aortic transection. Whole bladder, pancreas, gastrocnemius, left inguinal mammary fat pad and anterior subcutaneous white adipose tissue were collected. A ~5mm^2^ section of the left & right cardiac ventricle, lower left lung and sagittal section of the brain were collected and rinsed in ice cold phosphate buffered saline, before being manually diced using two scalpels in a scissor-action. Diced organs were then placed into 1mL of ice-cold lysis buffer (1X RIPA buffer, 1% Phosphatase Inhibitor, 1% Protease Inhibitor, 1% Anti-foam Y-30) in M-tubes (gentleMACS™ Tubes, Miltenyi Biotec). Organs were then mechanically homogenized using a gentleMACS Dissociator (Miltenyi Biotec) using program Protein_01. Tissues needing further homogenization were placed on ice for 5 minutes before running the program again, to avoid overheating. The M-tubes were then spun at 200G for 2 minutes at 4°C and incubated on ice for 5 minutes. The homogenized tissue was then sonicated at 60% power for 1 minute in 10 second intervals (10s run, 10s rest, 110s total). The debris was pelleted via centrifugation at 10,000G for 15 minutes at 4°C. The supernatant was then placed in QIAshredder tubes (Qiagen) and run at full speed for 2 minutes at 4°C. The flowthrough was aliquoted and placed in −80°C for storage.

### Immunoblotting

Protein concentration was quantified using Pierce BCA Protein Assay (Thermo Scientific, USA) and normalized to 40 μg of total protein for each tissue sample. After heat denaturation, reduced samples were resolved by SDS-PAGE on 4-20% gels and transferred on to nitrocellulose membranes. Blots were blocked in 2.5% milk in TBS with 0.1% Tween-20 (TBST) for 30 min at room temperature and then incubated with primary antibody overnight at 4°C. Blots were washed with TBST, incubated with HRP-conjugated secondary antibody for 1 hour at room temperature, and then visualized using Bio-Rad ChemiDoc™ Imaging System after detection with chemiluminescent substrate (SuperSignal™ West Pico, Thermo Scientific; or Radiance Plus, Azure Biosystems). Primary antibodies were extensively optimized. The following primary antibodies were used for detection: TIMP1 (Cell Signaling Technology, USA) at 1:500 dilution, TIMP2 (R&D Systems; AF971) 1:500, and TIMP3 (Cell Signaling Technology, USA) 1:750. Secondary anti-rabbit HRP-linked antibody (Cell Signaling Technology, USA) or anti-goat HRP-linked antibody (Invitrogen, USA) was used at 1:5000 dilution. Protein loading was determined by Ponceau Stain (Sigma Aldrich, USA). Immunoblot images were processed using Image Lab Software Version 6.1 (Bio-Rad). In-house produced recombinant human TIMP1/2/3 were included in the immunoblots at 4ng (TIMP1/2) or 10ng (TIMP3) per lane. Human TIMPs display high sequence identity with murine Timps (74.3%, 97.3%, 96.2% for Timp1/2/3, respectively).

### Multiplex RNA ISH staining of Mouse Timp1, Timp2, Timp3, and Timp4 RNA

Mouse lung, mammary, heart, and femoral skeletal muscle tissue were all harvested within 5 min and immediately placed into 10% neutral buffer formalin. Tissue was fixed for 72 hours before being processed into paraffin using a Tissue-Tek® Prisma™ (Sakura Finetek USA, Torrance, CA). One 5um section from each tissue type was stained with hematoxylin and Eosin (H&E) and scanned at 20X magnification using an Aperio AT2 (Leica Biosystems, Buffalo Grove, IL) into whole slide digital images.

Mouse Timp1, Timp2, Timp3, and Timp4 transcripts were detected by manually staining 5um FFPE tissue sections with RNAscope® Probe-Mm-Timp1-C3 (Cat No. 316841-C3), RNAscope® Probe-Mm-Timp2 (Cat No. 567831), RNAscope® Probe-Mm-Timp3-C2 (Cat No. 471311-C2), RNAscope® Probe-Mm-Timp4-C3 (Cat No. 1090531-C3), respectively, using the RNAscope™ Multiplex Fluorescent V2 Assay (Cat. No. 323270) with tissue pretreatment of 15 minutes at 95°C using Bond Epitope Retrieval Solution 2 (Leica Biosystems), 15 minutes of Protease III (ACD) at 40°C using the Bond RX auto-stainer (Leica Biosystems), and 1:750 dilution of TSA-Cyanine 5 Plus (AKOYA, SKU NEL745001KT) and TSA-Cyanine 3 Plus (AKOYA, SKU NEL744001KT). RNAscope® 3-plex Negative Control Probe (Bacillus subtilis dihydrodipicolinate reductase (*dapB*) gene in channels C1, C2, and C3, Cat# 320871) was used as a negative control. RNAscope® 3-plex Positive Control Probe-Mm (Cat# 320881) was used as a technical control to ensure the RNA quality of tissue sections were suitable for staining.

RNA ISH staining was followed by immunohistochemical staining of lung and mammary tissue with a 1:200 dilution of Vimentin (D21H3) XP® Rabbit mAb (Cell Signaling Technology, Ref# 5741) with a Bond RX auto-stainer using the Bond Polymer Refine Kit (Leica Biosystems, Cat# DS9800) minus the peroxidase block, post primary reagent, DAB and Hematoxylin. Cd34 protein was detected in heart and skeletal muscle by staining sections post RNAscope with a 1:100 dilution of Cd34(RAM34) Monoclonal Antibody (eBioscience™, Ref# 14-0341-82) using the Bond Polymer Refine Kit minus the peroxidase block, post primary reagent, DAB and Hematoxylin followed by Biotinylated Rabbit a/Rat (Vector Laboratories, cat# BA-4001) secondary antibody. The binding of antibodies were detected with a 1:150 dilution of OPAL 520 Reagent (AKOYA, SKU FP1487001KT). Slides were digitally imaged using an Aperio ScanScope FL Scanner (Leica Biosystems).

Image analysis was performed using Halo imaging analysis software (v3.4.2986.246, Indica Labs, Corrales, NM), and image annotations were performed independently by two pathologists (WGSS & BK). Halo algorithm (FISH Amplification/Deletion v1.4) were used to quantify Timp expression.

### Data harvest and processing

Single-cell transcriptomic data generated through FACS (fluorescence-activated cell sorting) or microfluidic-droplet methods, as well as individual cell annotations, were downloaded from Tabula Muris Figshare (https://figshare.com/articles/dataset/Single-cell_RNA-seq_data_from_Smart-seq2_sequencing_of_FACS_sorted_cells_v2_/5829687)(https://figshare.com/articles/dataset/Single-cell_RNA-seq_data_from_microfluidic_emulsion_v2_/5968960). Brain, colon, fat, pancreas, and skin tissues only have FACS data. Bladder, heart, aorta, kidney, liver, lung, mammary, marrow, muscle, spleen, thymus, tongue, and trachea organs have data from both methods. In our data analyses the Timp expression data from aorta was separated from heart, and limb muscle (which we refer to as skeletal muscle) was separated from the diaphragm. Additionally, brain myeloid and non-myeloid organs were combined into one organ. As a result, there are 17 organs analyzed (see Table S1) compared to the original 20 organs in the Tabula Muris dataset. Seurat (V4) R package (https://satijalab.org/seurat/) was used for data quality control (QC), normalization, integration, clustering, visualization as well as differential expression (DE) analyses.

For organs that only have FACS data, SCTransform [42] was applied for normalization before clustering. For tissues with datasets from both methods, due to the observed technical batch effect between the two methods, Seurat integration pipeline [43] was applied along with SCTransform normalization (https://satijalab.org/seurat/articles/integration_introduction.html).

When studying specific cell types, based on annotations, gene raw counts data were retrieved for those cell types followed by the same normalization or integration workflow. For tissue-specific analyses of cell-types, cells obtained from the aorta and diaphragm tissue were separated from heart and skeletal muscle datasets, respectively. Expression visualizations were generated manually using Adobe Photoshop and the circlepackeR R package.

### Clustering

Principal components (PCs), explaining most variance, were selected based on their rankings displayed by Seurat’s Elbowplot() function. Selected PCs for the different datasets ranged between 30-40. FindNeighbors() function was run to generate a nearest neighbor graph with the identified PCs as dimensions of reductions with the default settings. Clusters of cells were identified by a shared nearest neighbor (SNN) modularity optimization-based algorithm by utilizing the FindClusters() function at the default 0.8 resolution. Clusters are visualized with Uniform Manifold Approximation and Projection (UMAP) dimensional reduction technique with the selected PCs.

### Differential expression (DE) and pathway analysis

Based on the UMAP clusters, different clusters and/or groups of clusters were selected for differential expression analysis. Differentially expressed genes were identified for groups and/or clusters with the FindMarkers(?) function. For samples that were integrated to remove the batch effect, a logistic regression framework was used with batch information as a latent variable. For samples without multiple batches, default Wilcoxon Rank Sum test was used with SCT assay data. Identified markers were filtered for Bonferroni corrected adjusted p-values <0.05. Pathway analysis was performed using Ingenuity Pathway Analysis (Qiagen) and MetaCore (Clarivate).

## Abbreviations

TIMPs/Timps: Tissue inhibitors of metalloproteinases
MMP: Metzincin proteases of the Matrix Metalloproteinase/Metallopeptidase
ADAM: A Disintegrin and Metalloproteinase/Metallopeptidase
ADAMS-TS: A Disintegrin and Metalloproteinase/Metallopeptidase with thrombospondin motifs
scRNA sequencing: single cell RNA sequencing
RNA ISH: RNA in-situ hybridization

## Acknowledgements

This research was supported by the Intramural Research Program of the NIH (W.G.S.-S. Project ID ZIA SC 009179) and with Federal funds from the National Cancer Institute, National Institutes of Health, under Contract No. HHSN261201500003I. The content of this publication does not necessarily reflect the views or policies of the Department of Health and Human Services, nor does mention of trade names, commercial products, or organizations imply endorsement by the U.S. Government (BK, AW).

## Animal Studies

Animals were maintained under specific pathogen-free conditions at NCI Bethesda. All animal procedures reported in this study that were performed by NCI-CCR affiliated staff were approved by the NCI Animal Care and Use Committee (ACUC) and in accordance with federal regulatory requirements and standards. All components of the intramural NIH ACU program are accredited by AAALAC International.

## Contributions

D.P. and W.G.S.S. conceived the study. D.P., Y.F. and S.R. performed data harvest and analysis. S.G. and C.L. performed tissue harvest and immunoblots. A.W. and B.K. performed tissue processing, staining and imaging for RNA ISH experiments. All authors assisted in interpretation of data. The manuscript was written by D.P. and W.G.S.S. All authors reviewed the manuscript.

## Ethics Declarations

The authors declare no competing interests.

## Data Availability

The authors confirm that the data supporting the findings of this study are available within the article [and/or] its supplementary materials. Example code for tissue-specific and cell-type specific data collection can be found in supplementary Files S1 & S2.

---

**Figure S1. Full immunoblots.**

**Figure S2. Whole tissue UMAPs depicting cell clusters and Timp expression in the Tabula Muris dataset.**

**Figure S3. UMAPs depicting clusters and Timp expression in select tissues from the Tabula Muris dataset.**

**Figure S4. RNA ISH of mammary tissue depicting Timp2 expression throughout the fibrous connective tissue.** Red arrows indicate fibrous connective tissue, blue arrows indicate large capillaries.

**Figure S5. RNA ISH of mammary tissue depicting Timp2/3 expression throughout a peripheral nerve and skeletal muscle co-harvested with the mammary tissue.** Red arrows indicate nervous tissue, blue arrows indicate skeletal muscle, orange arrows indicate vasculature.

**Figure S6. Inflammation of the murine lungs.** (A-C) Hematoxylin and eosin staining of the murine used in RNA ISH studies. (D) 10X RNA ISH image depicting enhanced Timp1 RNA expression throughout inflamed region of murine lung.

**Figure S7. RNA ISH and H+E the cardiac vessel within the interventricular septum.**

**Figure S8. RNA ISH/IHC negative control images**

