## Supplemental Figure 1 for "Whole Organism Profiling of the Timp Gene Family"

**Figure S1. Full immunoblots.**

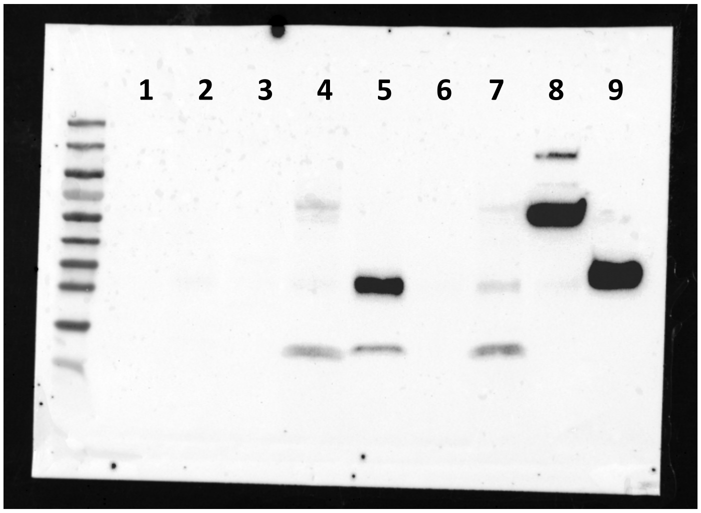


Timp1

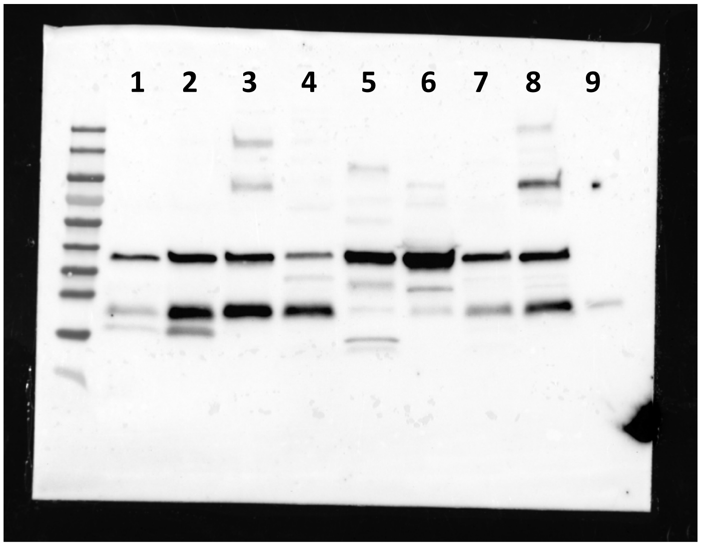


Timp2

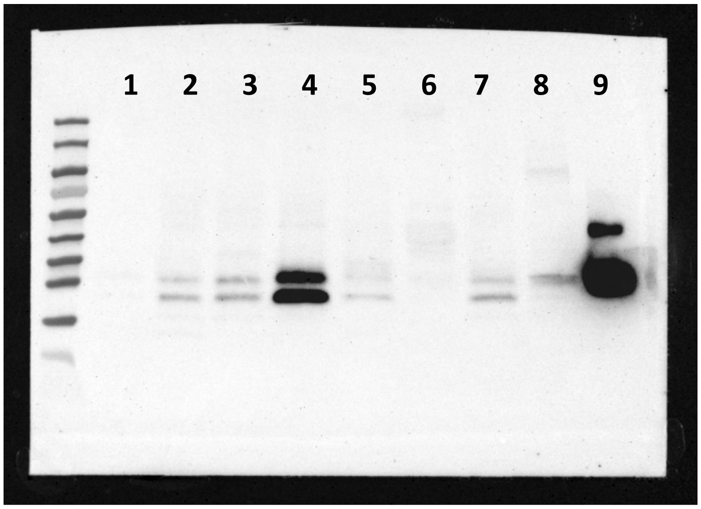


Timp3

1: Mammary fat pad
2: Adipose tissue
3: Bladder
4: Lung
5: Cardiac
6: Skeletal muscle
7: Pancreas
8: Brain
9: Timp
