## Supplemental Figure 2 for "Whole Organism Profiling of the Timp Gene Family"

Adipose Tissue_UMAP Cluster (PC = 30)

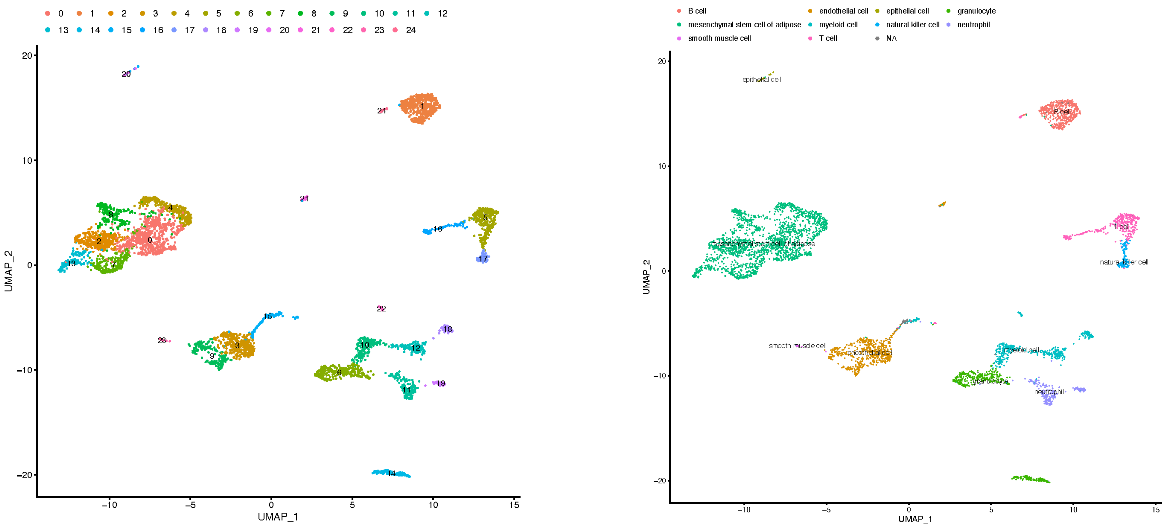


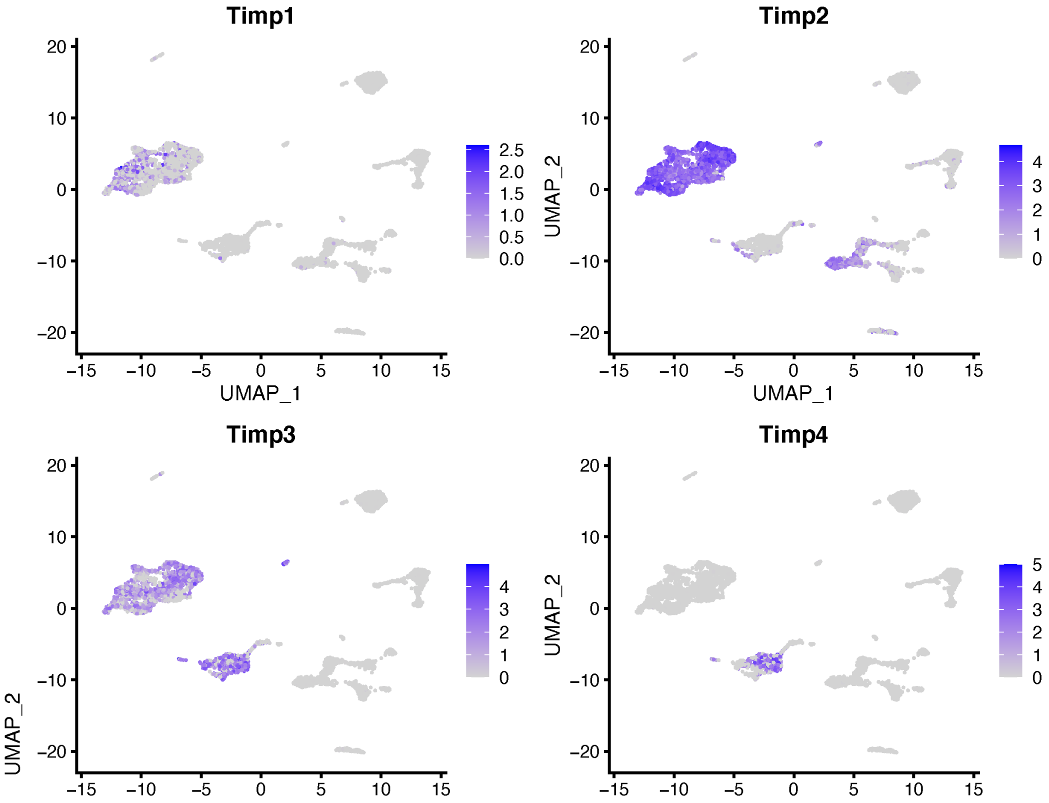

### Bladder Tissue_UMAP Cluster (PC = 30)

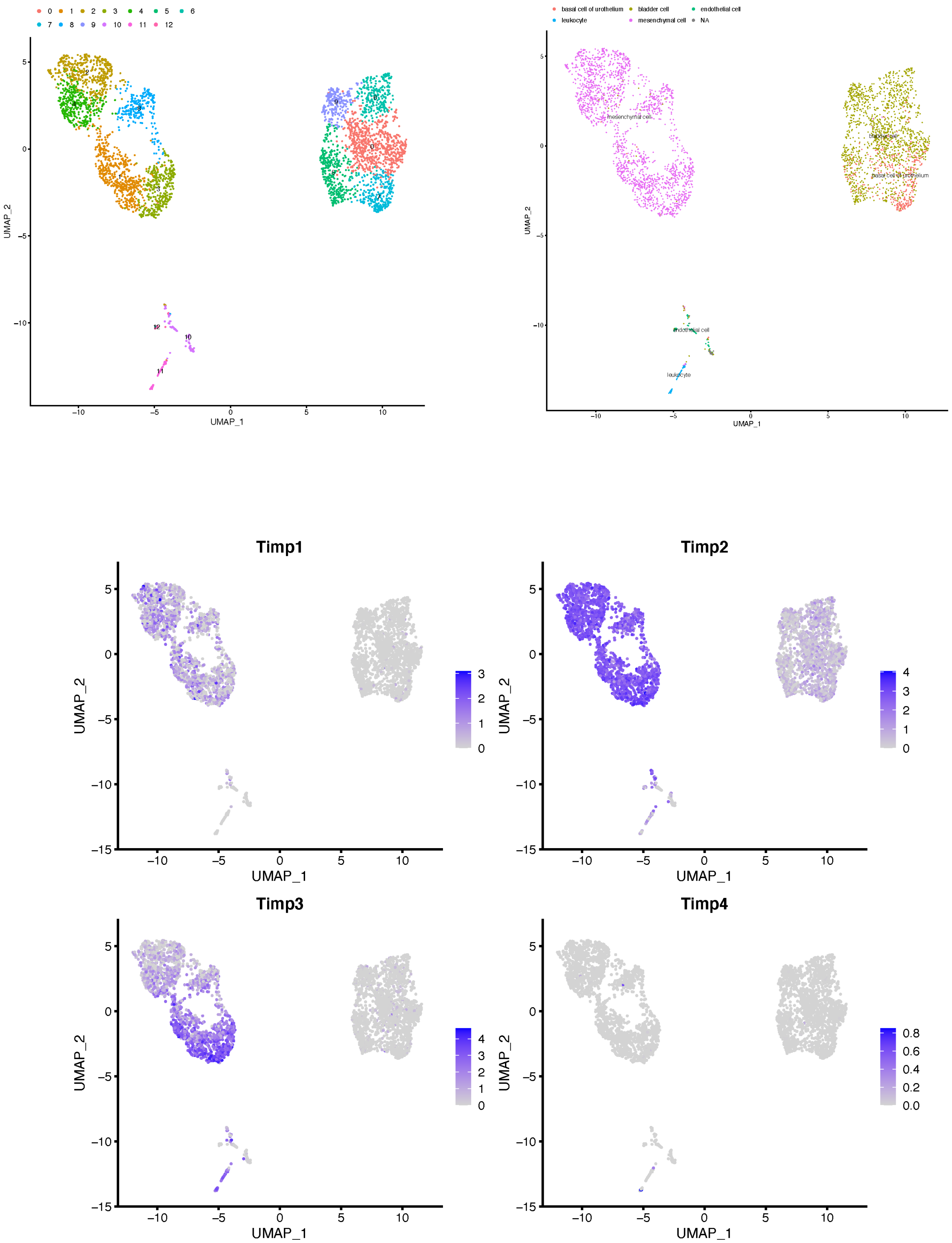

### Bone Marrow Tissue_UMAP Cluster (PC = 40)

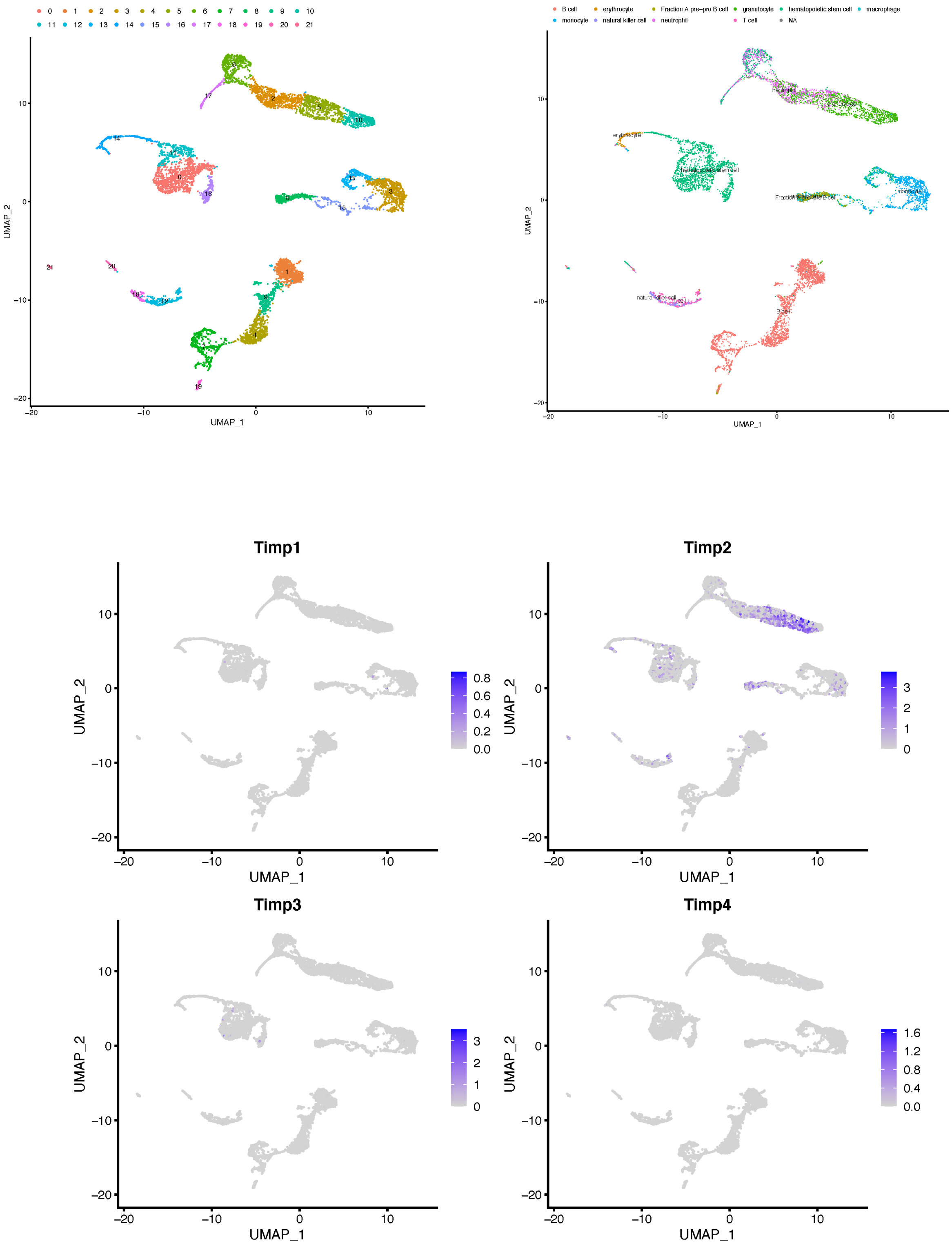

Brain Tissue_UMAP Cluster (PC = 30)

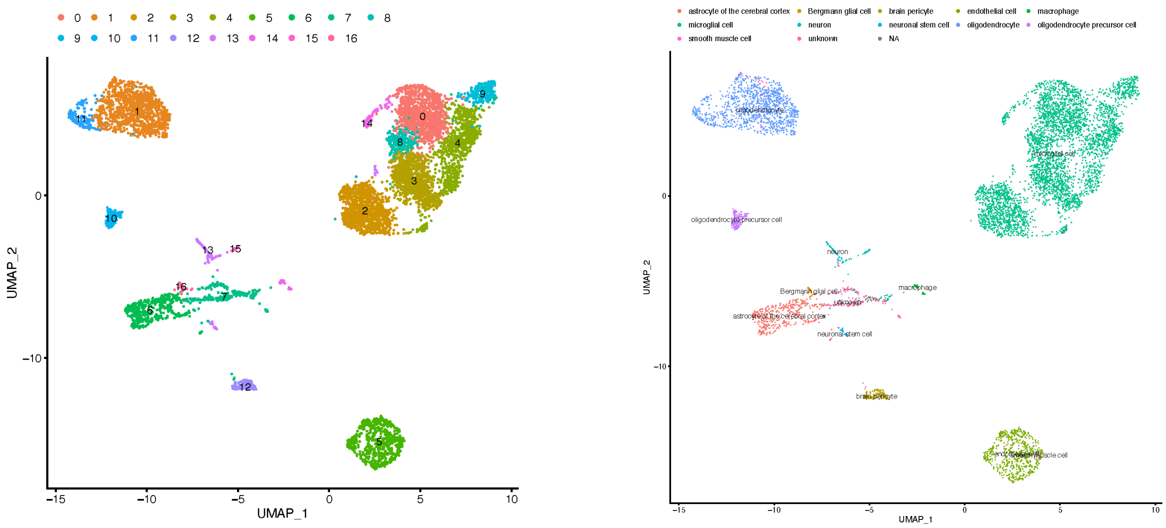


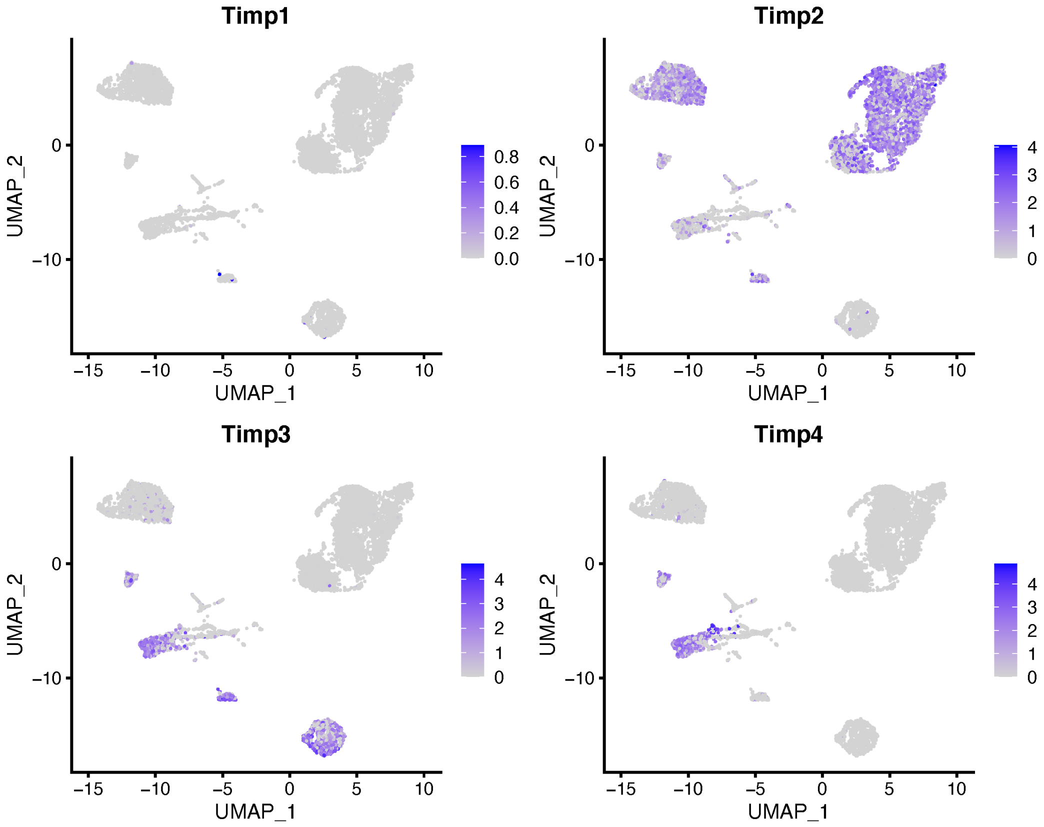

Colon Tissue_UMAP Cluster (PC = 30)

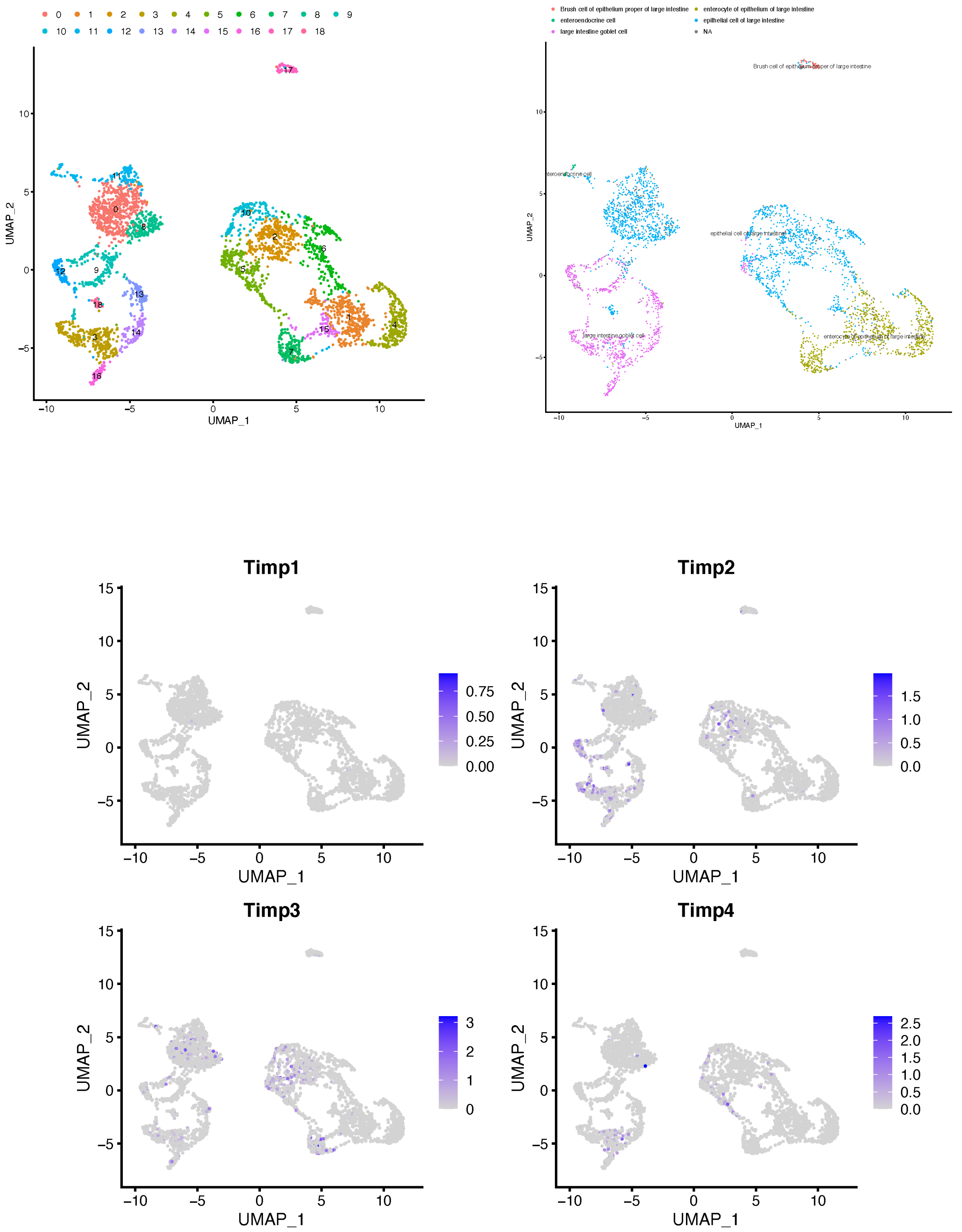

### Heart Tissue_UMAP Cluster (PC = 40)

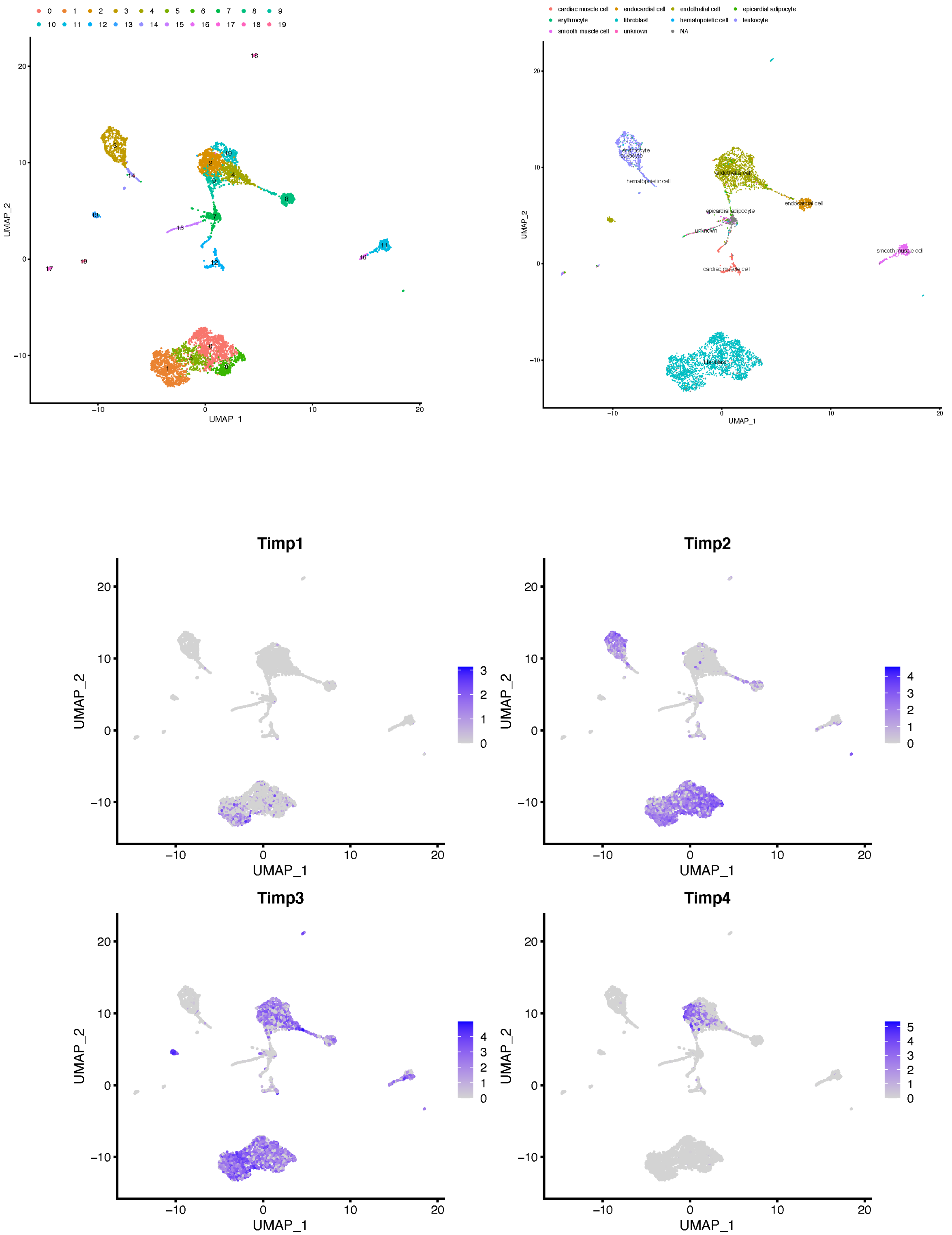

Kidney Tissue_UMAP Cluster (PC = 40)

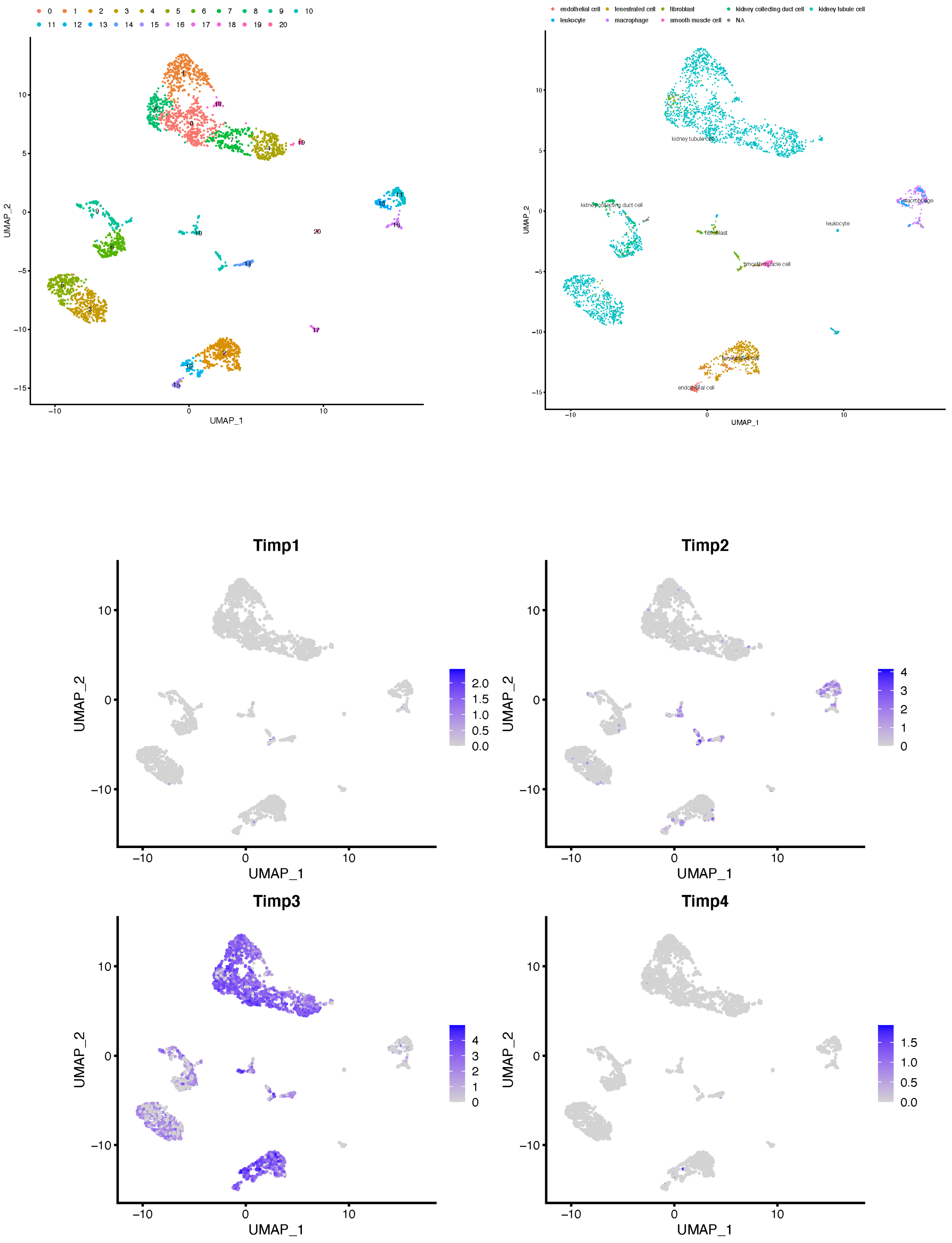

### Liver Tissue_UMAP Cluster (PC = 30)

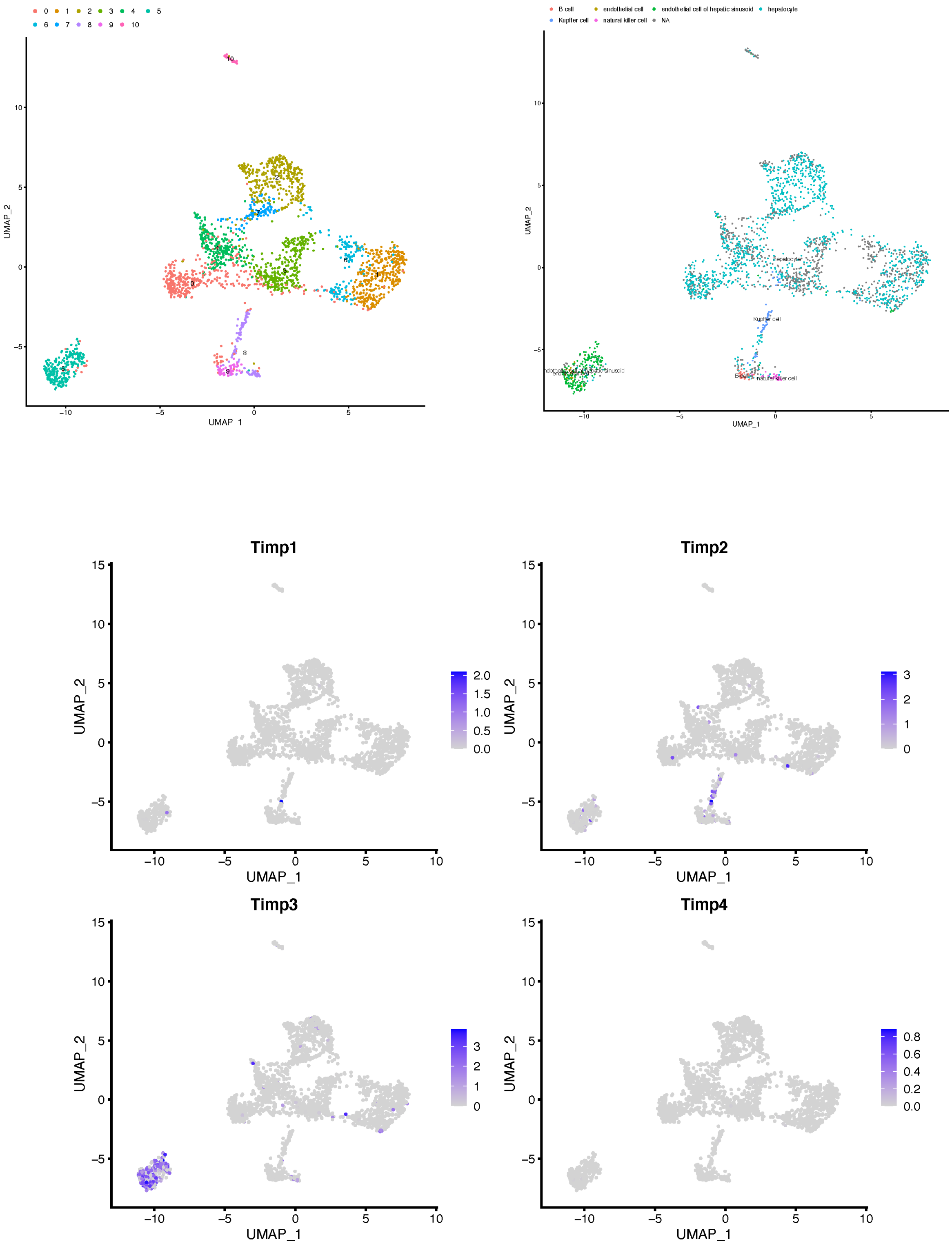

### Lung Tissue_UMAP Cluster (PC = 30)

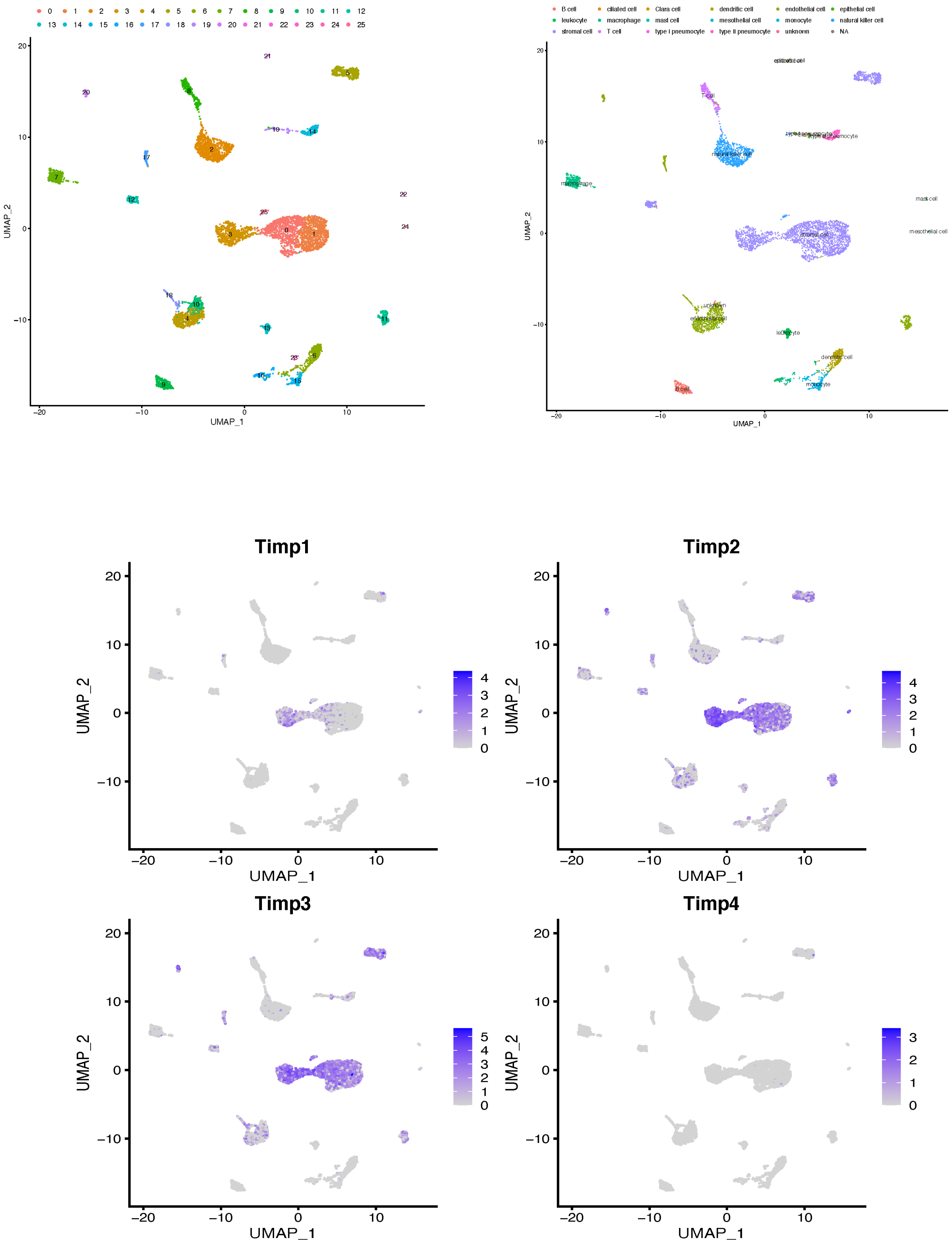

Mammary Tissue_UMAP Cluster (PC = 40)

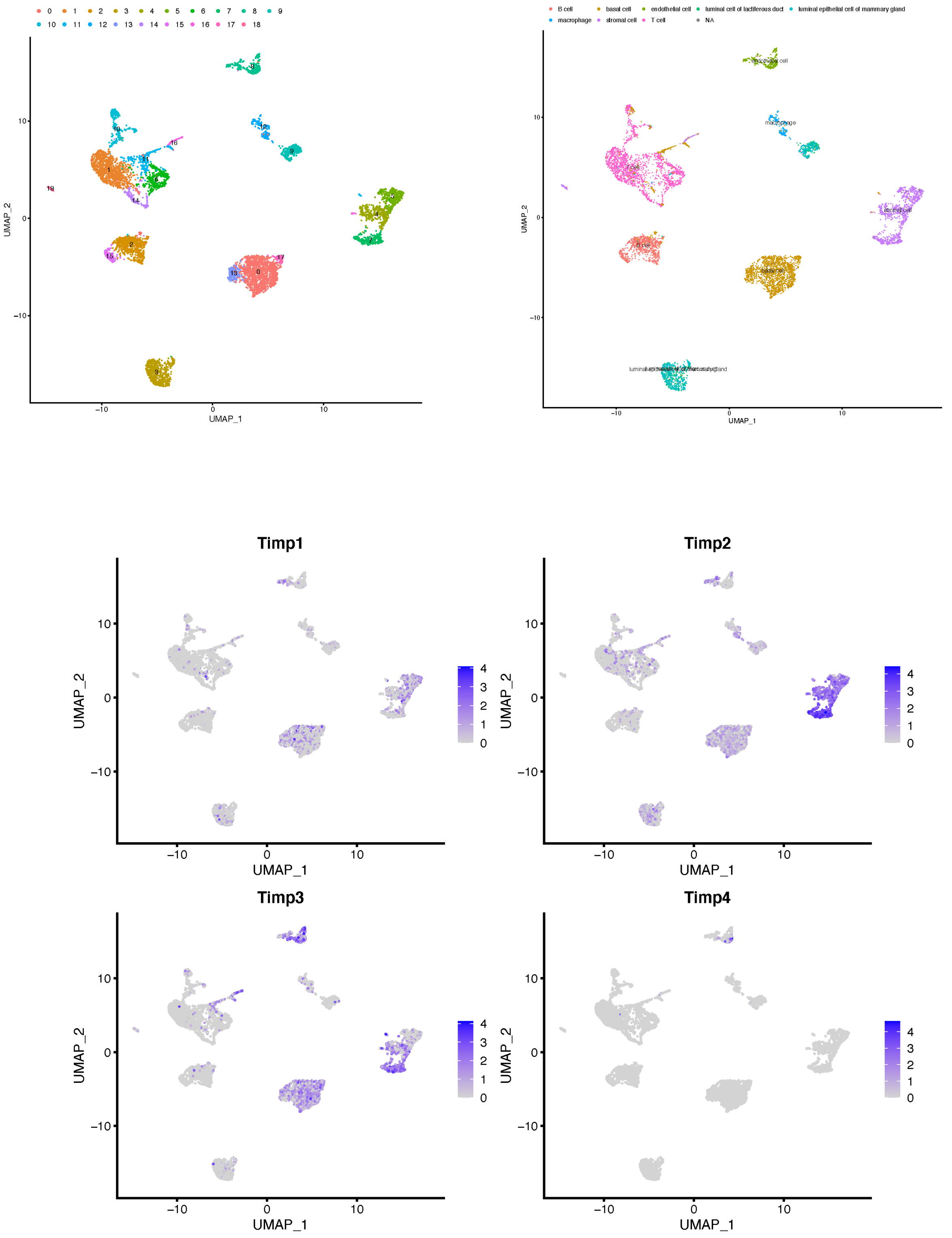

### Pancreas Tissue_UMAP Cluster (PC = 30)

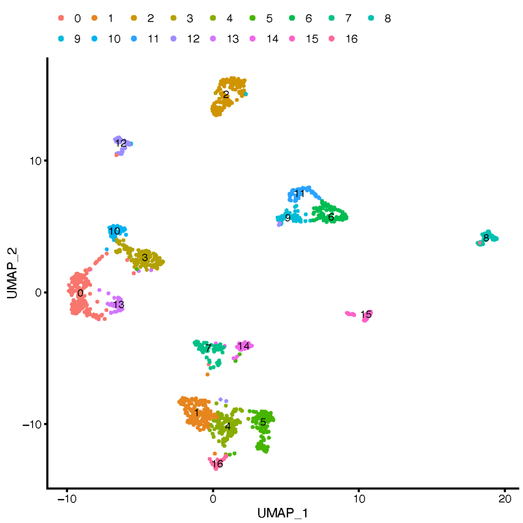


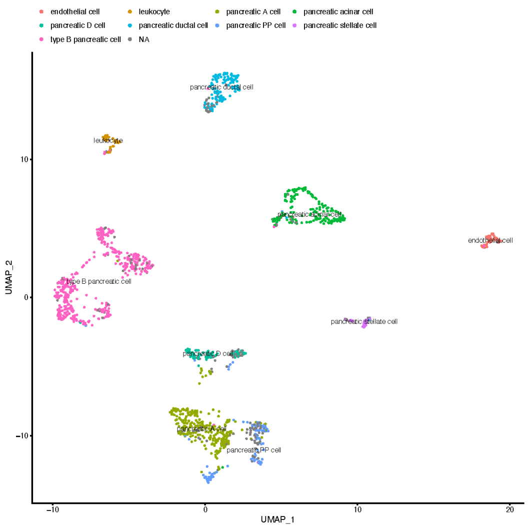


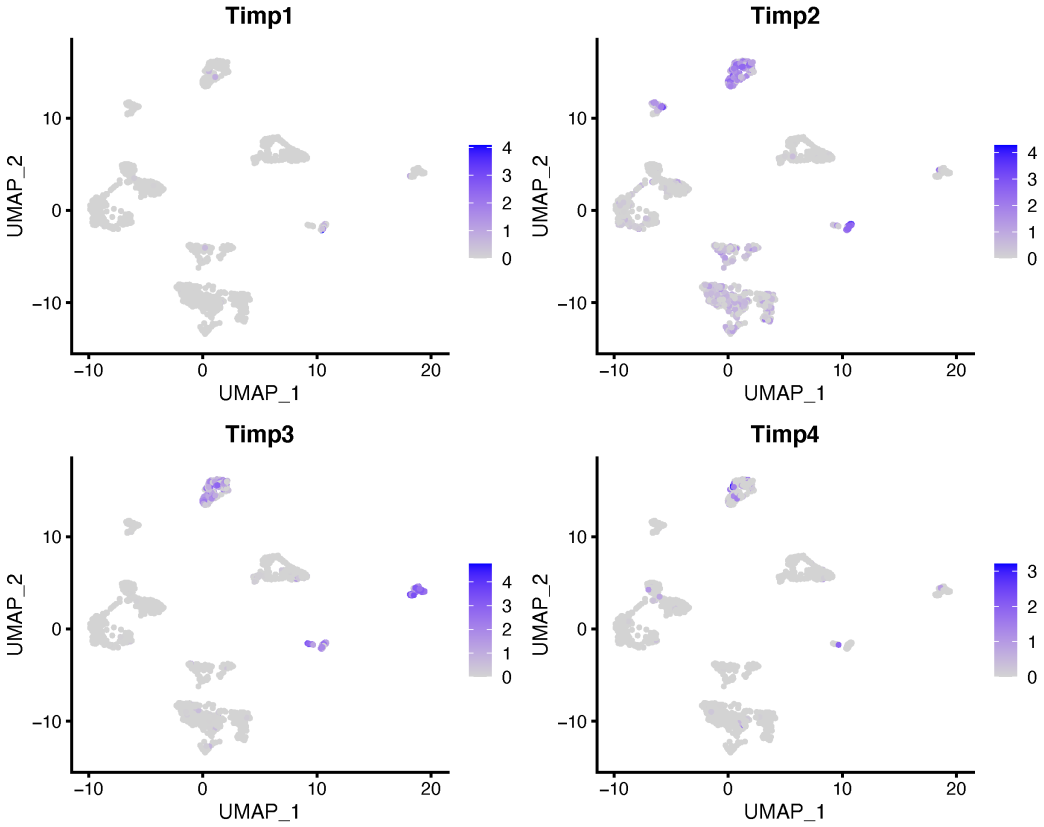

### Skeletal Muscle Tissue_UMAP Cluster (PC = 40)

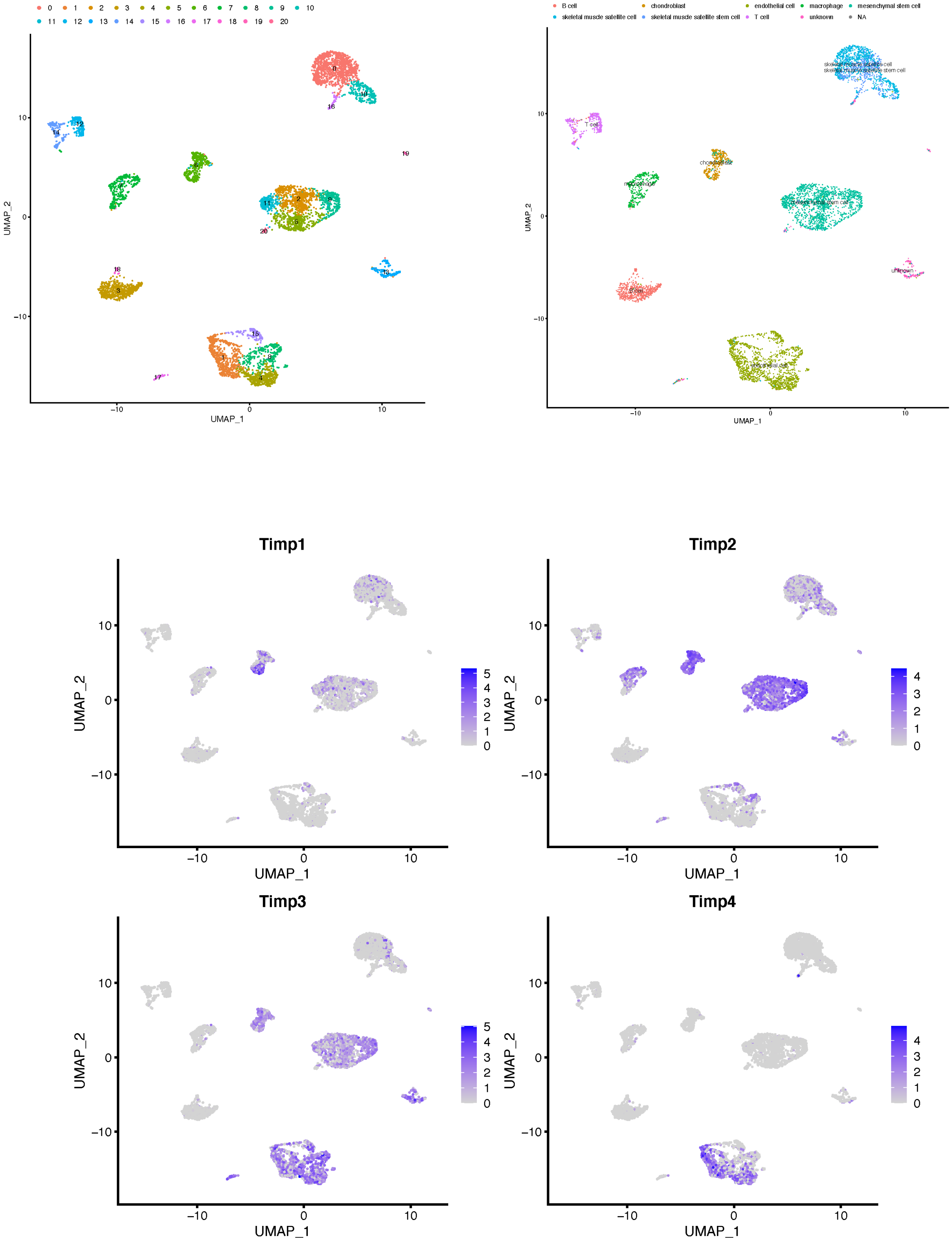

### Skin Tissue_UMAP Cluster (PC = 20)

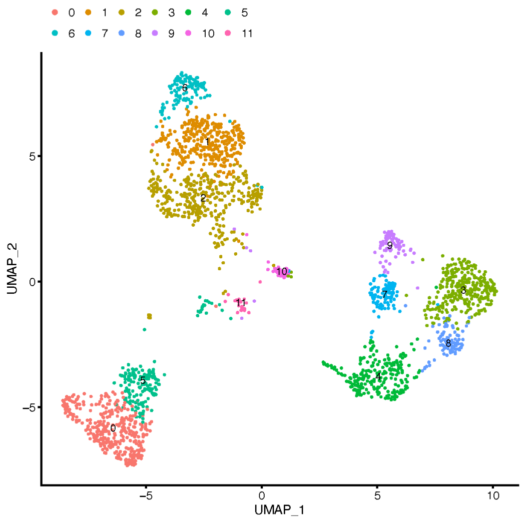


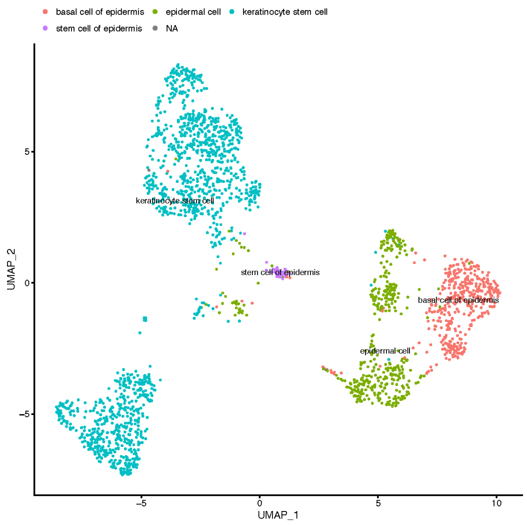


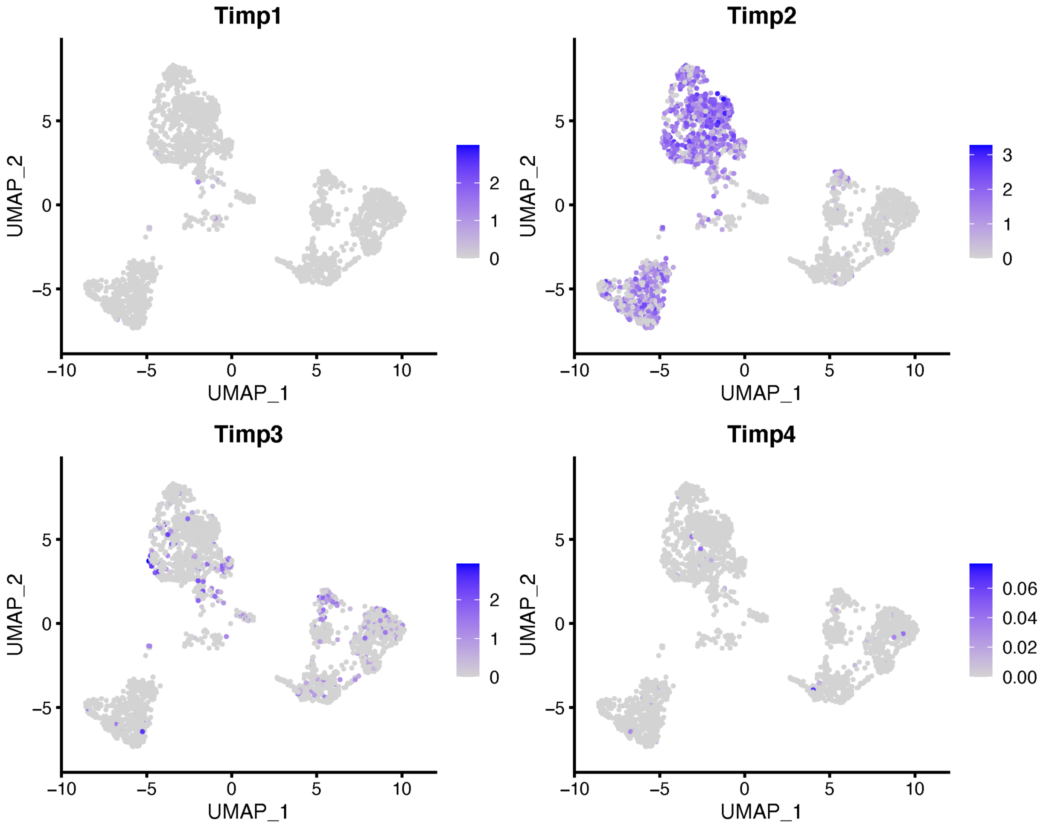

### Spleen Tissue_UMAP Cluster (PC = 40)

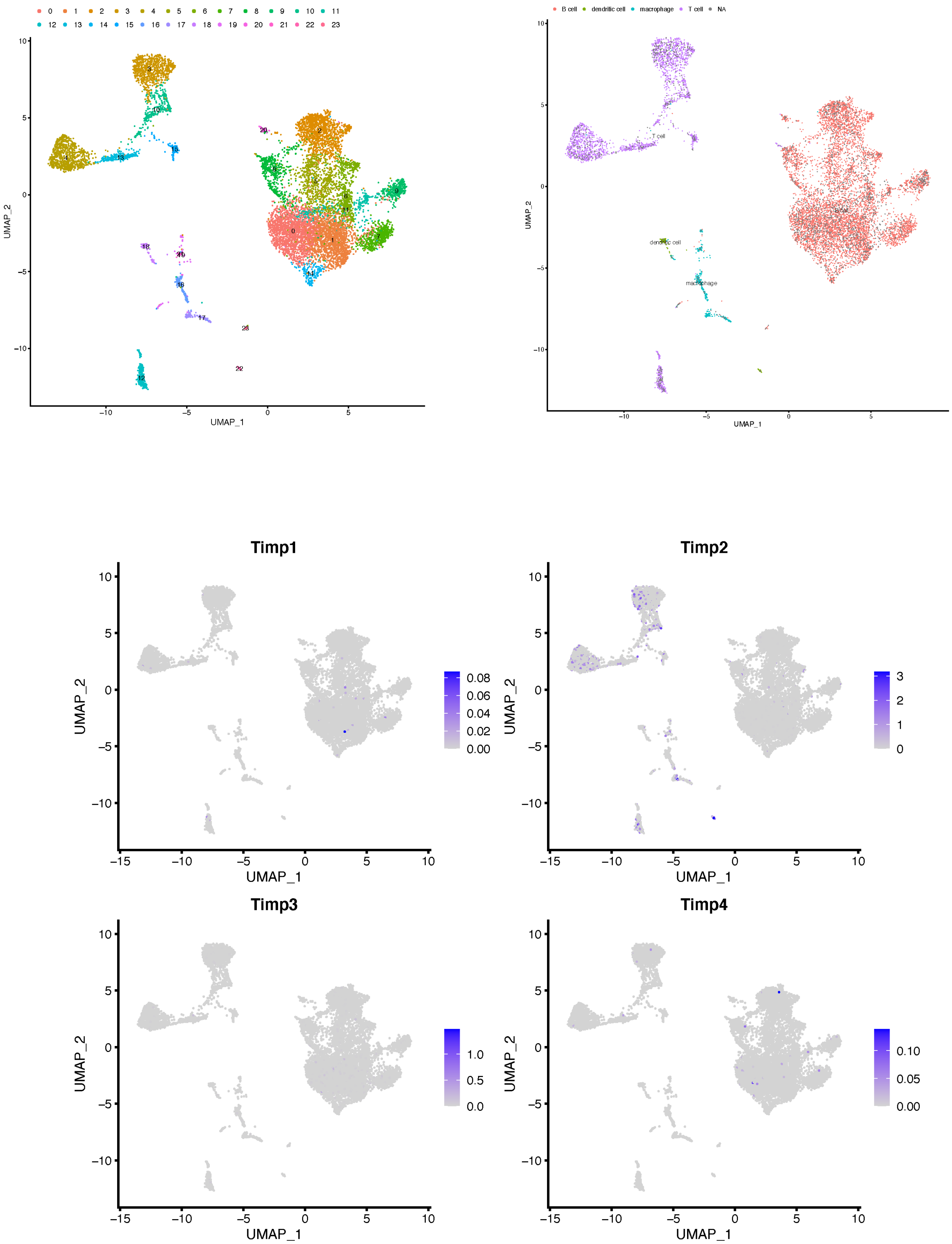

### Thymus Tissue_UMAP Cluster (PC = 40)

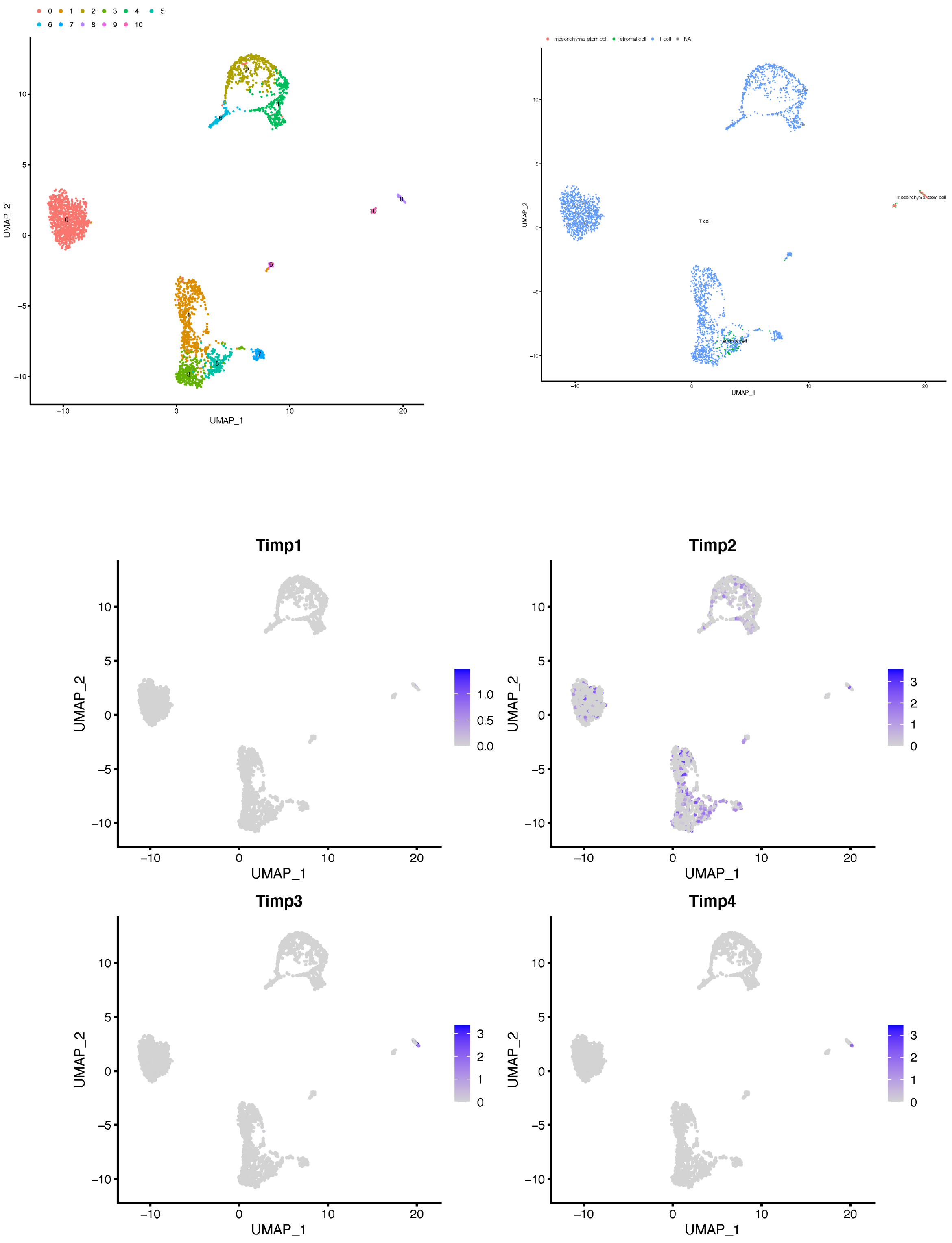

### Tongue Tissue_UMAP Cluster (PC = 30)

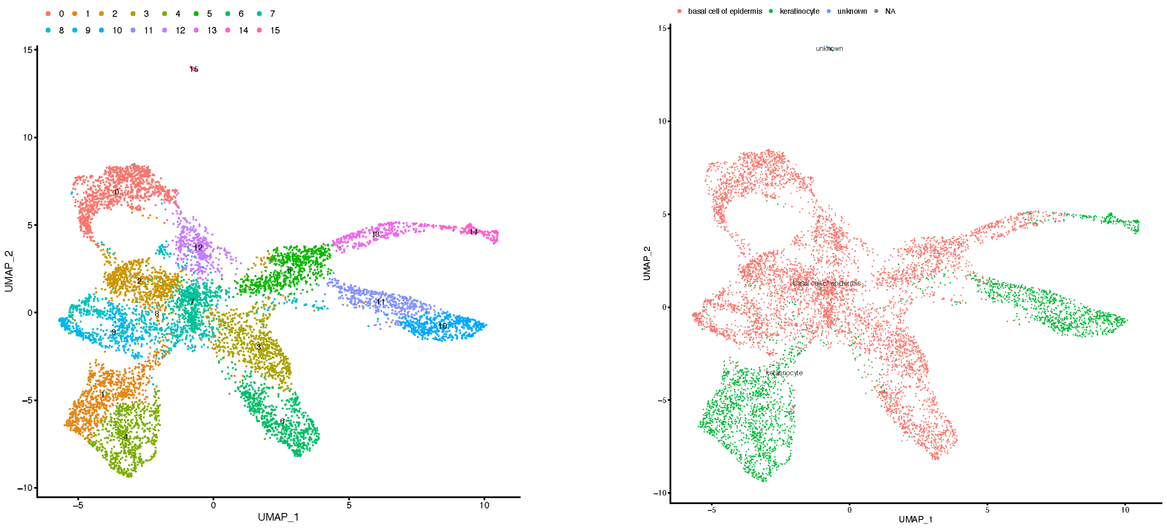


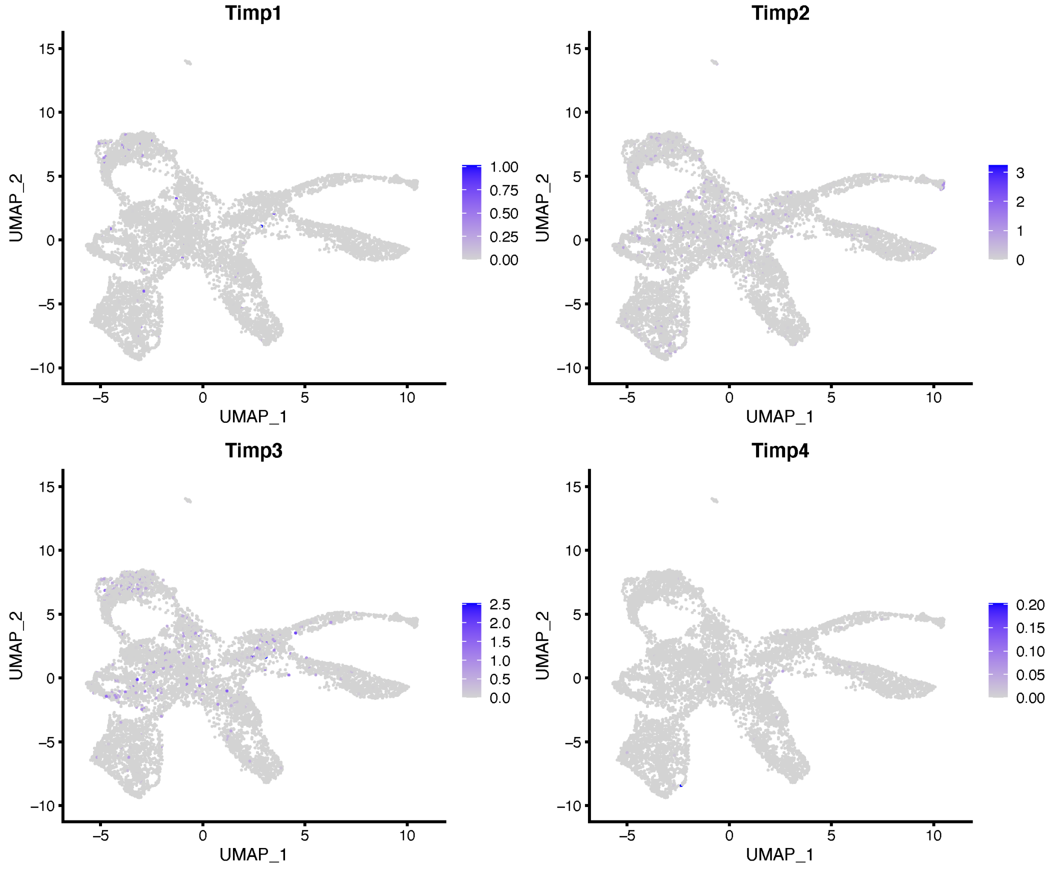

### Trachea Tissue_UMAP Cluster (PC = 40)

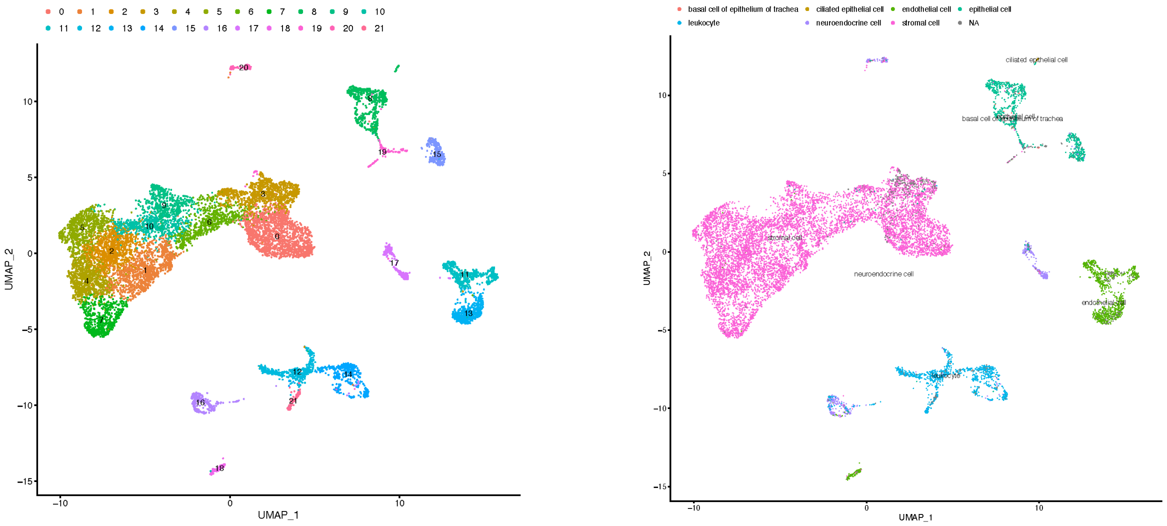


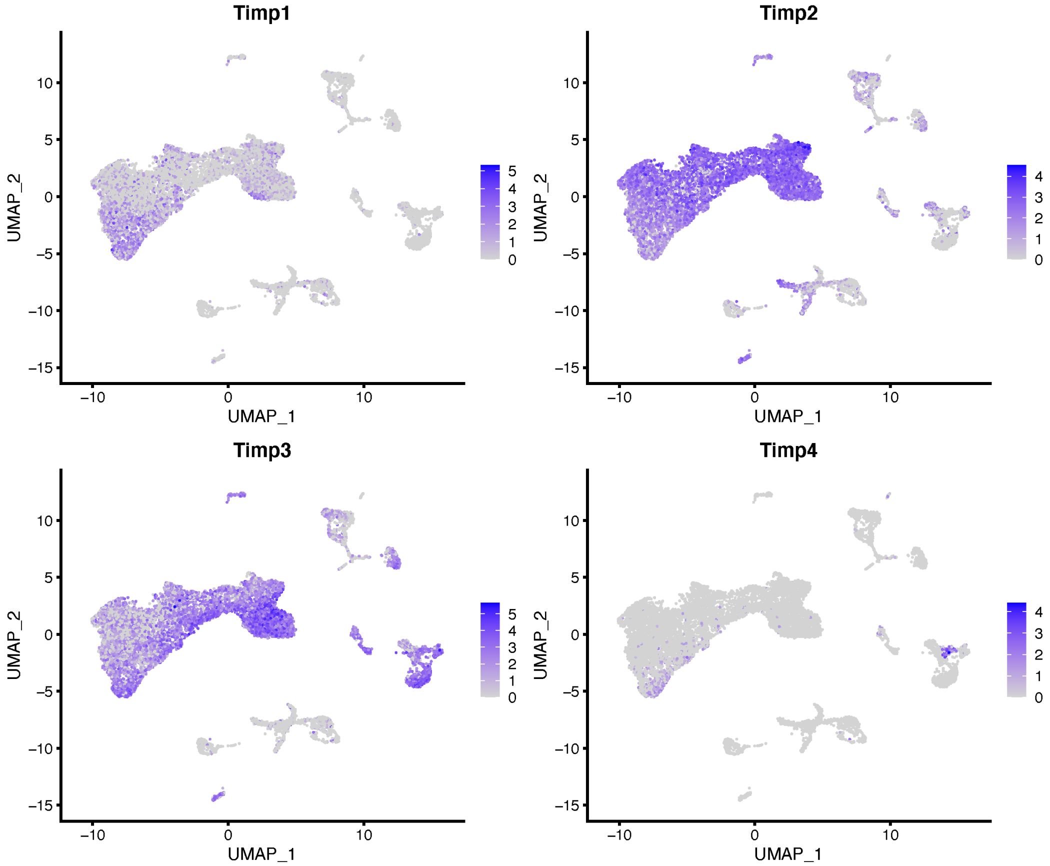
