## Supplemental Figure 3 for "Whole Organism Profiling of the Timp Gene Family"

### Adipose Endothelial Cells_UMAP Cluster (PC = 30)

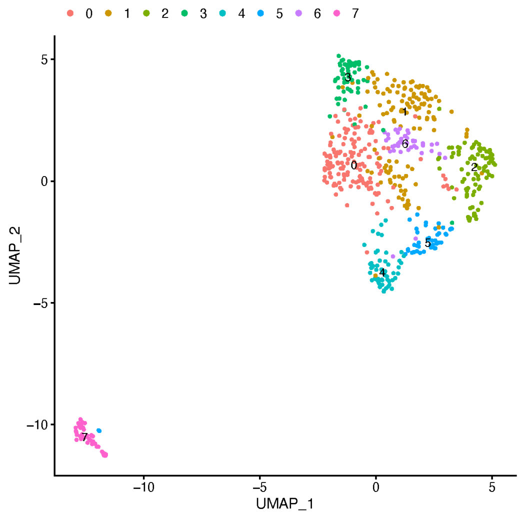


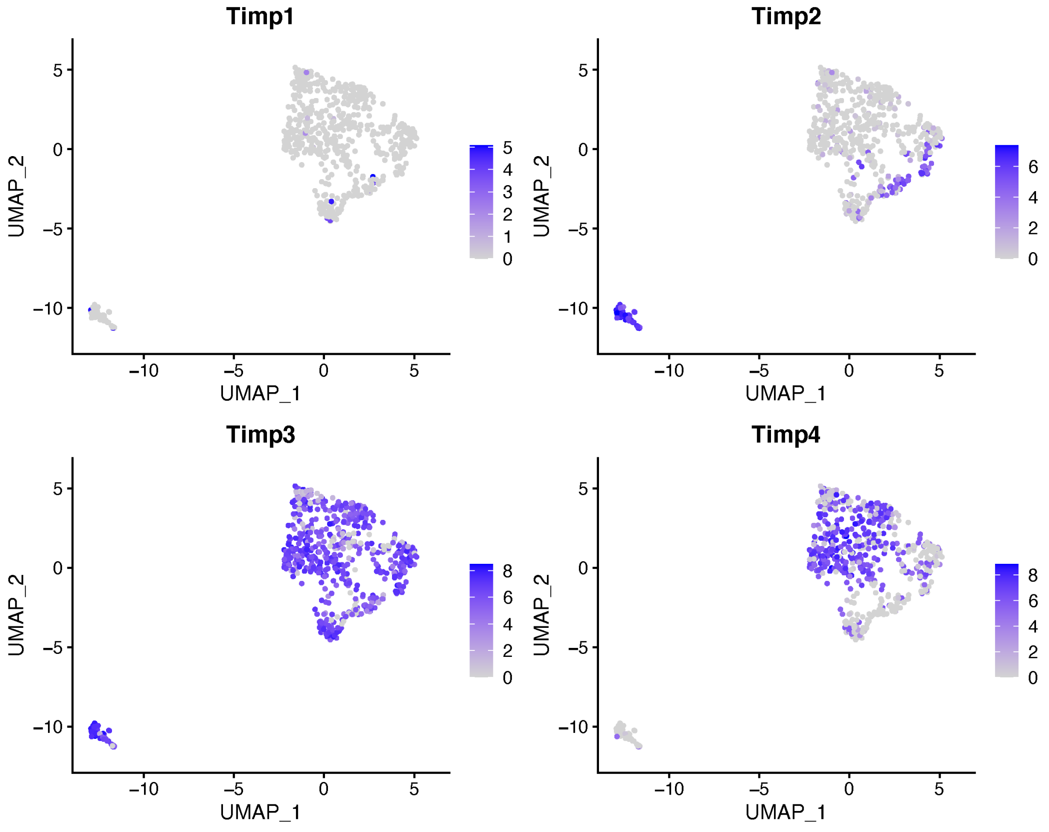

Adipose Granulocytes_UMAP Cluster (PC = 30)

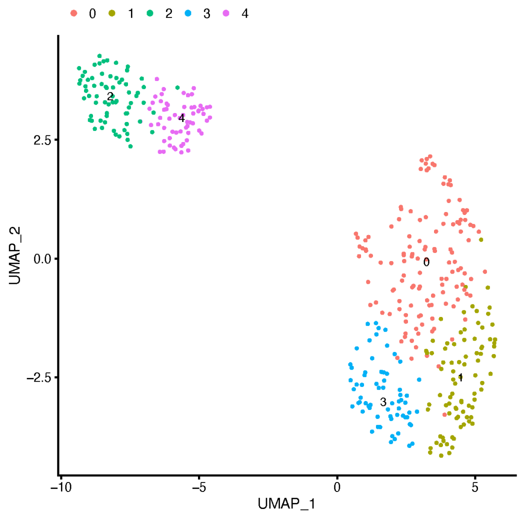


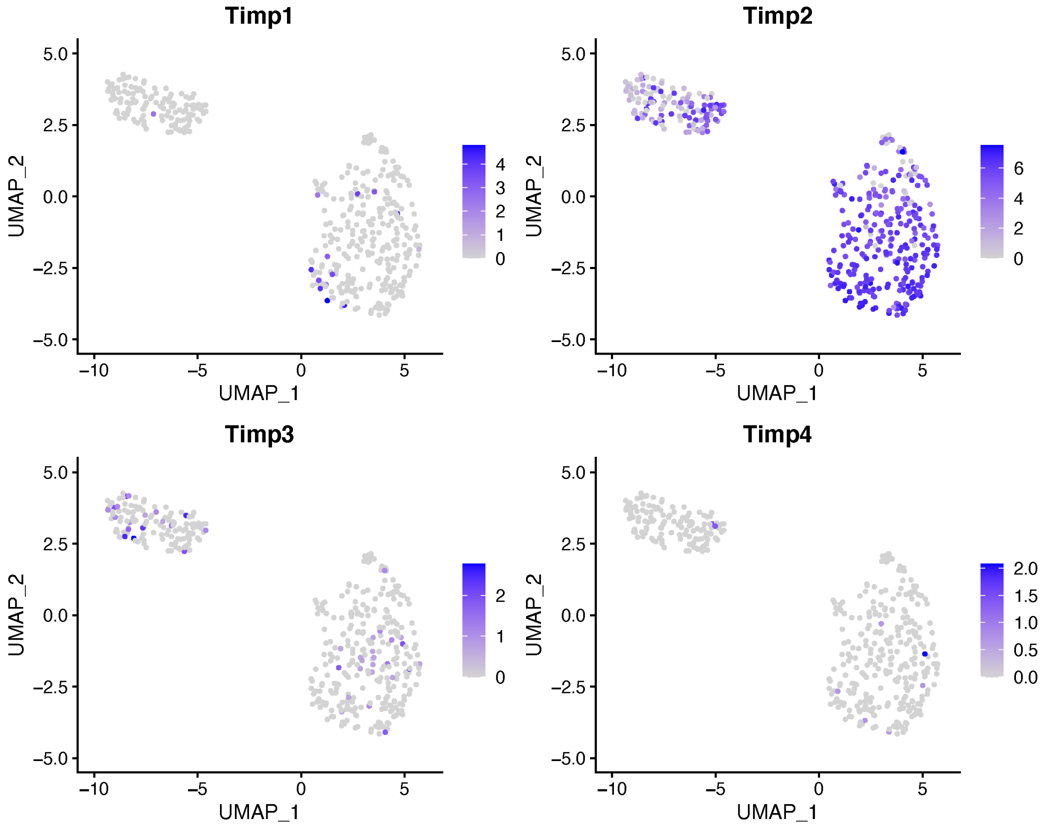

### Adipose Mesenchymal Stem Cells_UMAP Cluster (PC = 30)

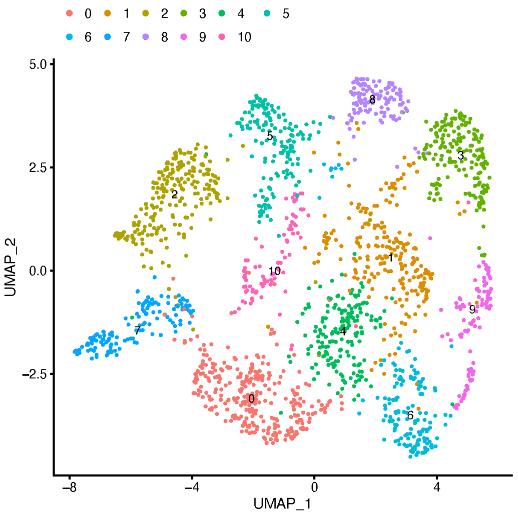


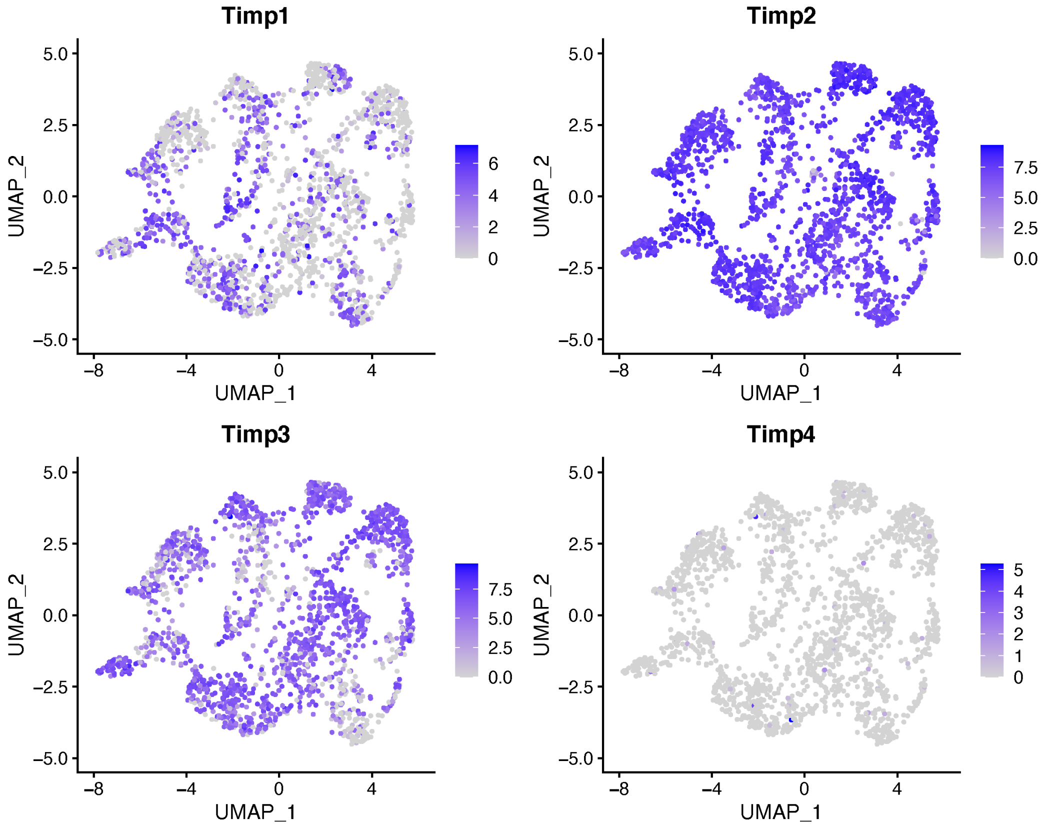

### Bladder Mesenchymal Cells_UMAP Cluster (PC = 30)

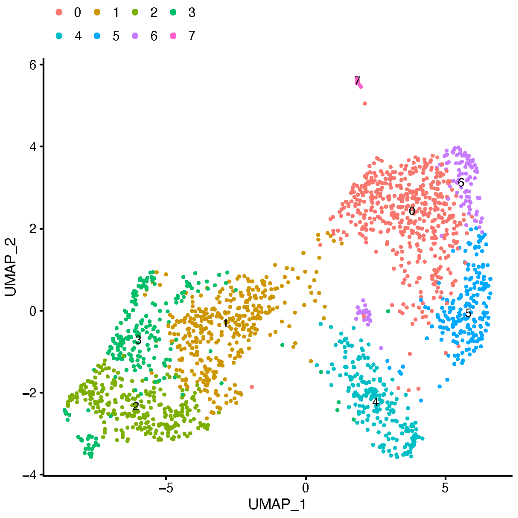


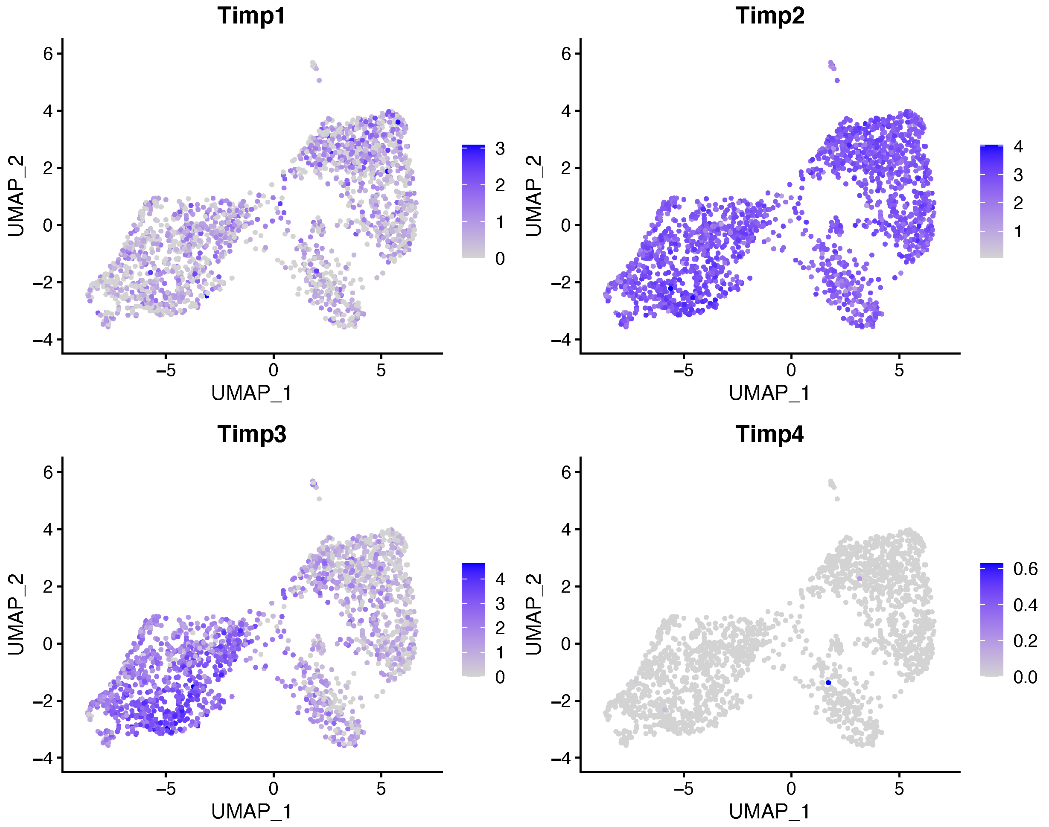

Bone Marrow Granulocytes_UMAP Cluster (PC = 40)

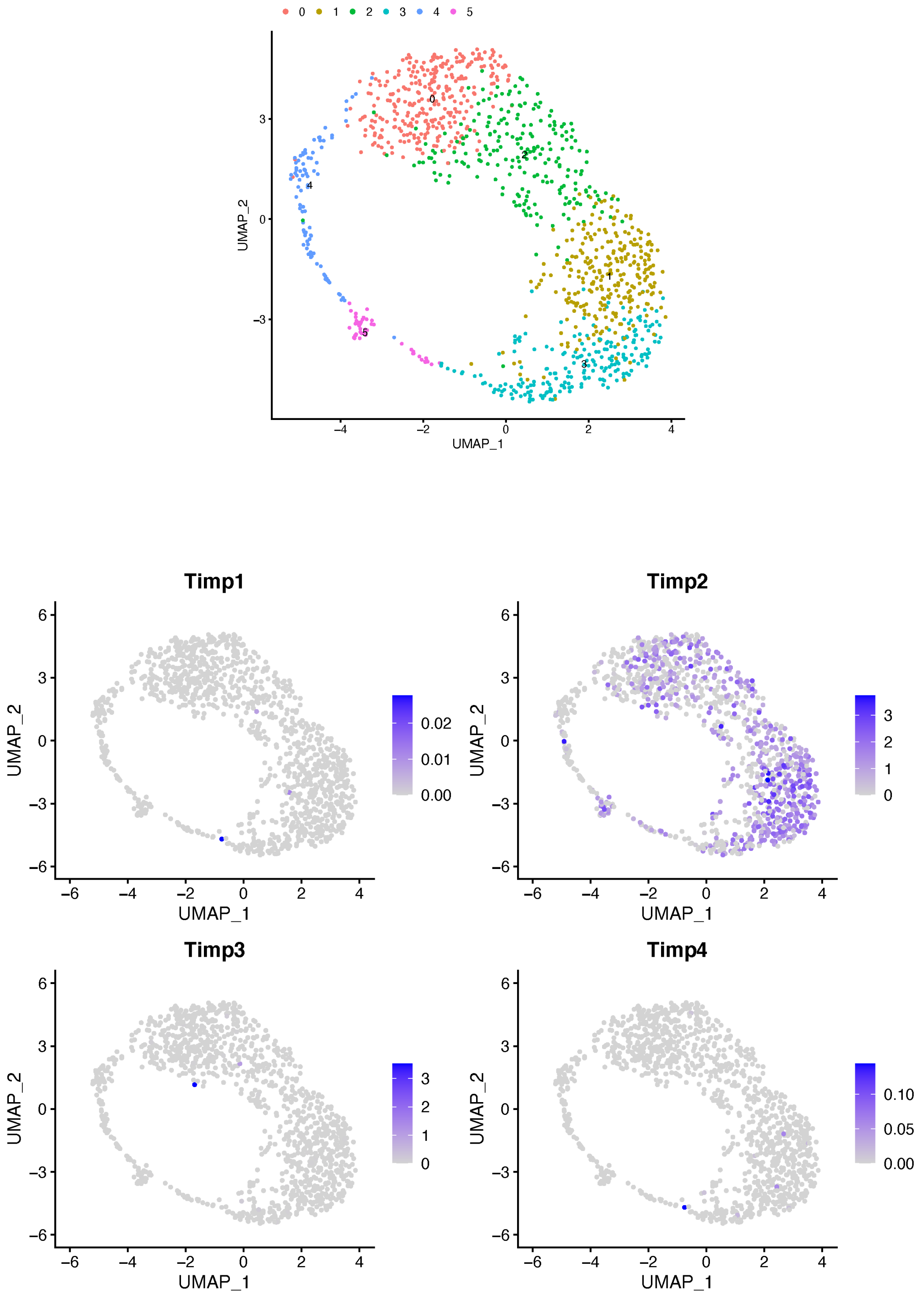

### Brain Astrocytes and Bergmann Glial Cells_UMAP Cluster (PC = 40)

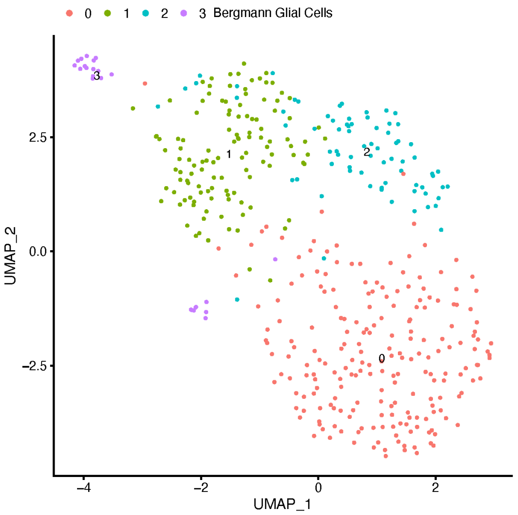


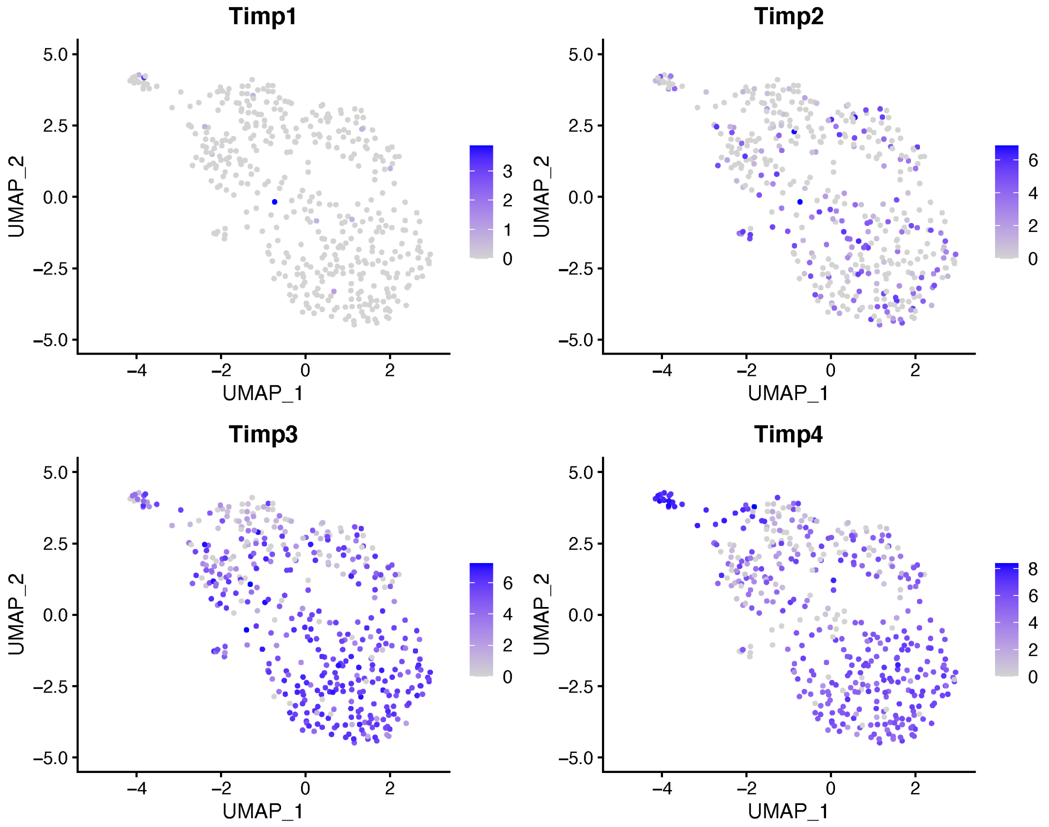

### Brain Endothelial Cells_UMAP Cluster (PC = 40)

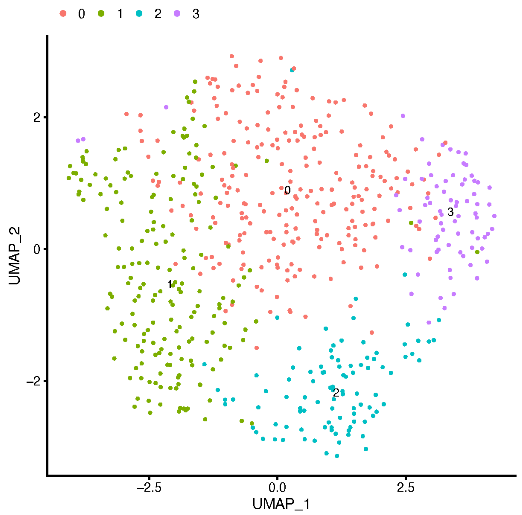


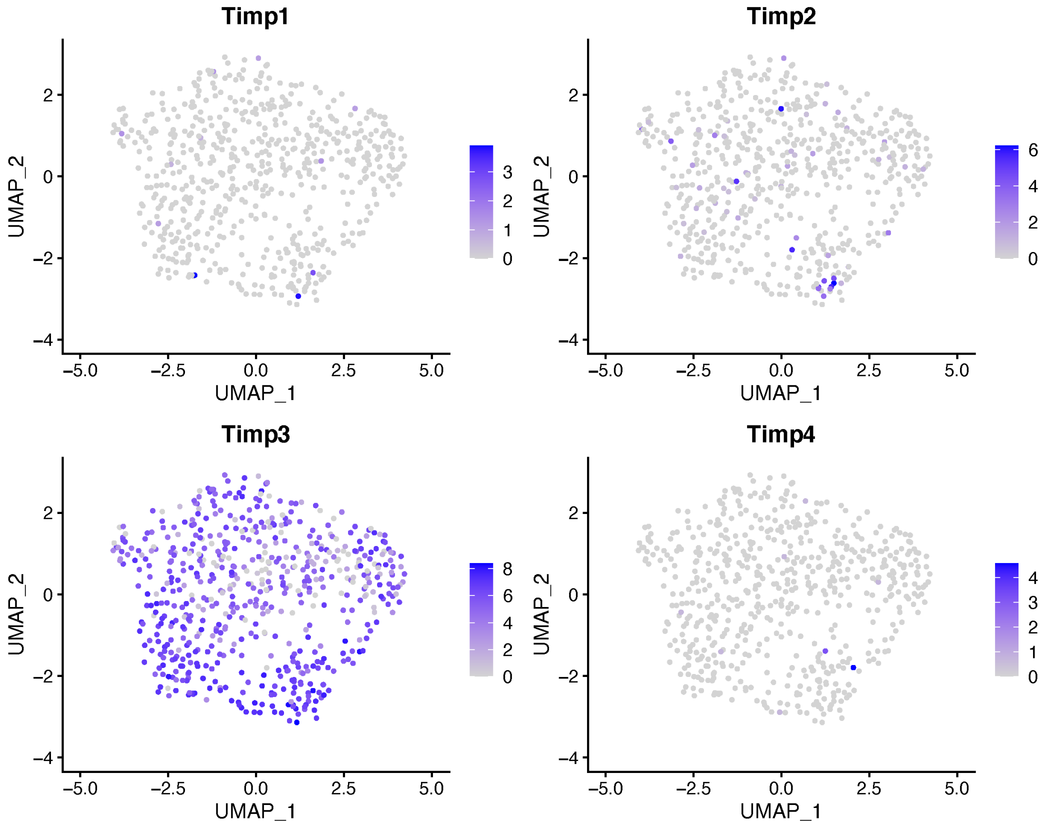

### Heart Endothelial Cells_UMAP Cluster (PC = 40)

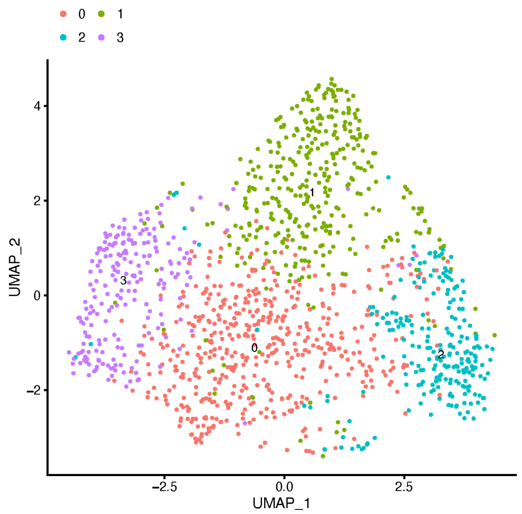


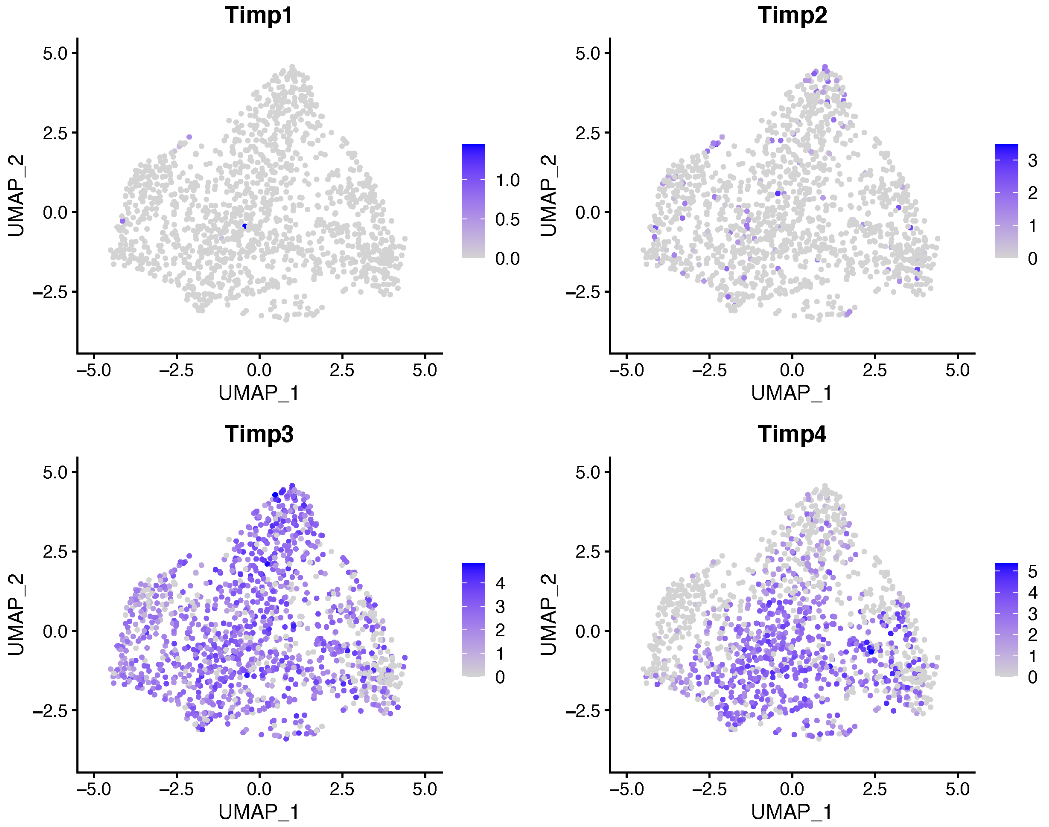

### Heart Fibroblasts_UMAP Cluster (PC = 40)

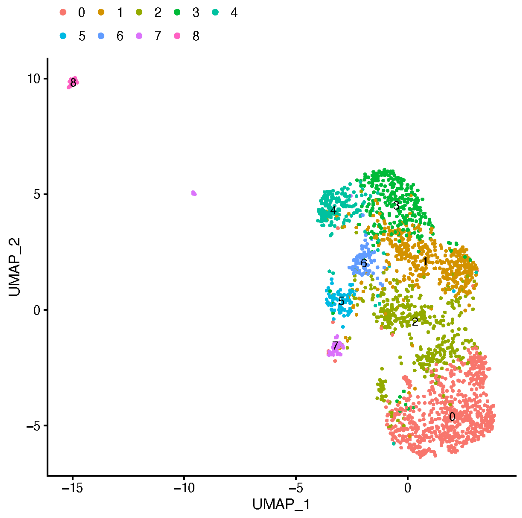


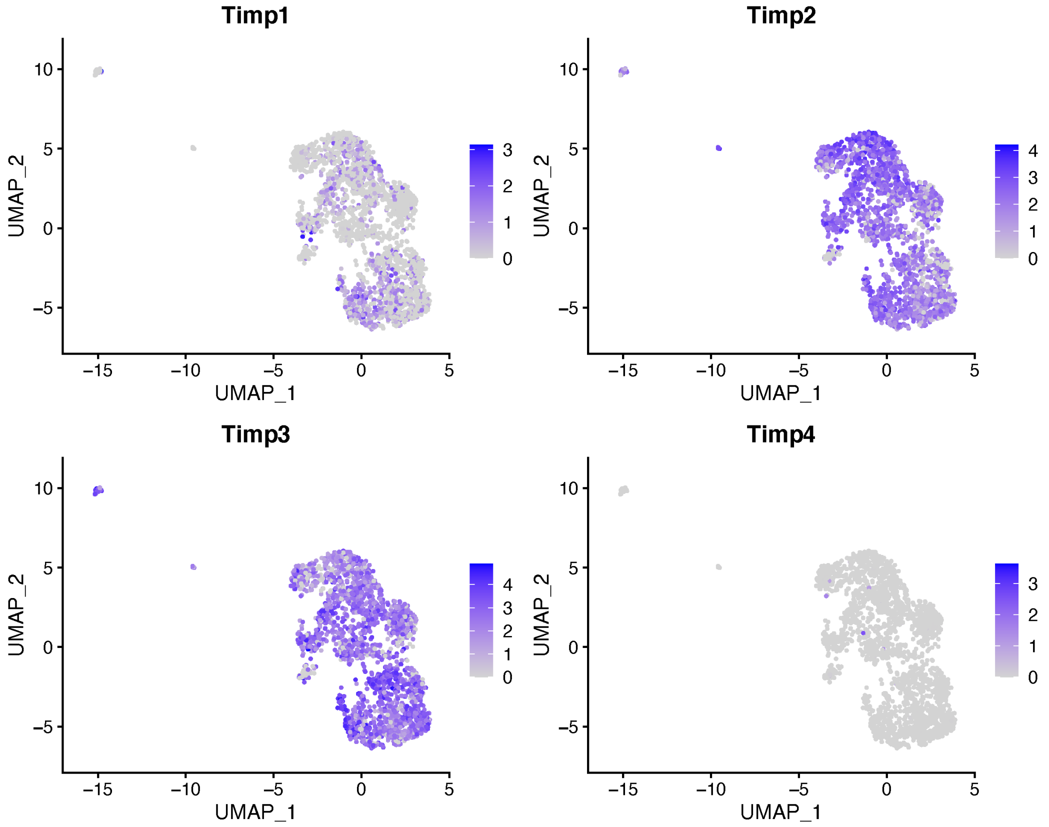

Kidney Tubule Cells_UMAP Cluster (PC = 30)

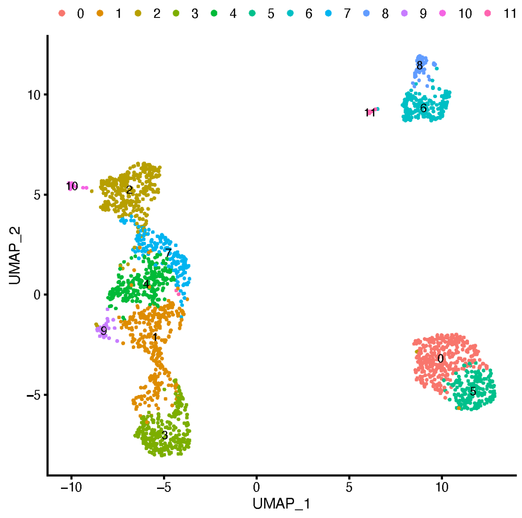


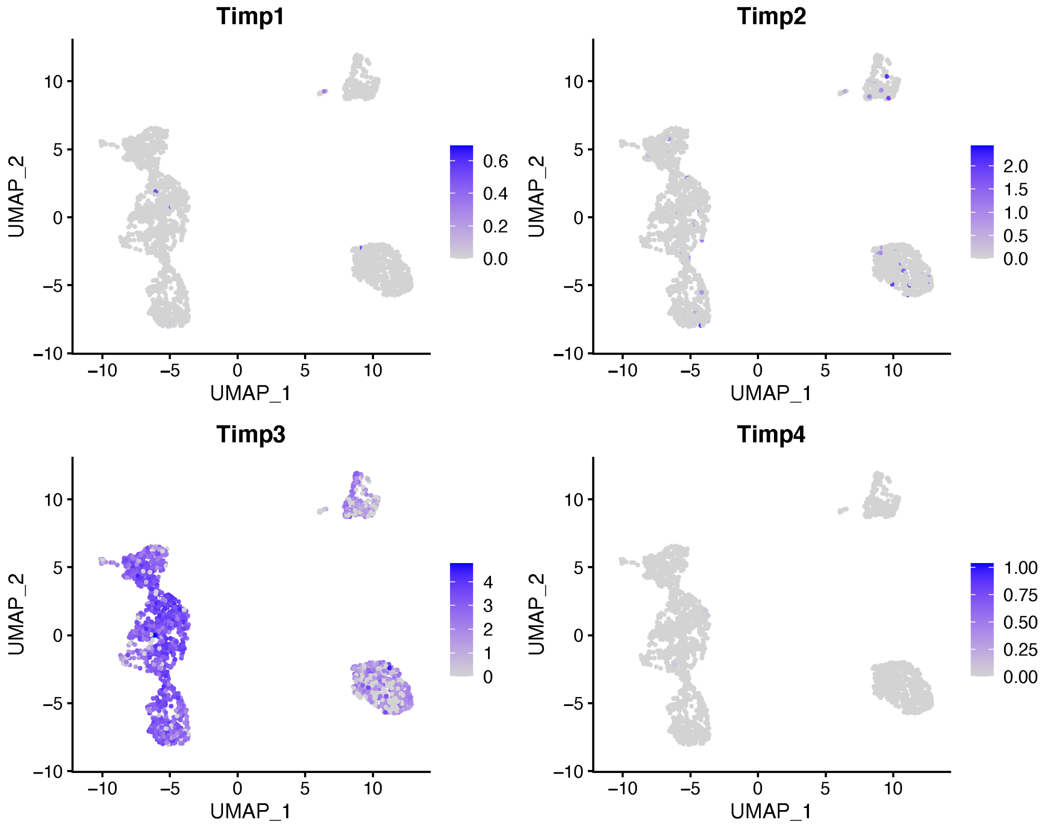

Lung Stromal Cells_UMAP Cluster (PC = 30)

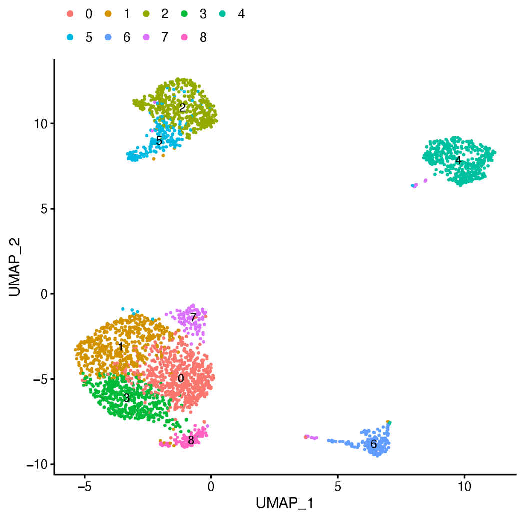


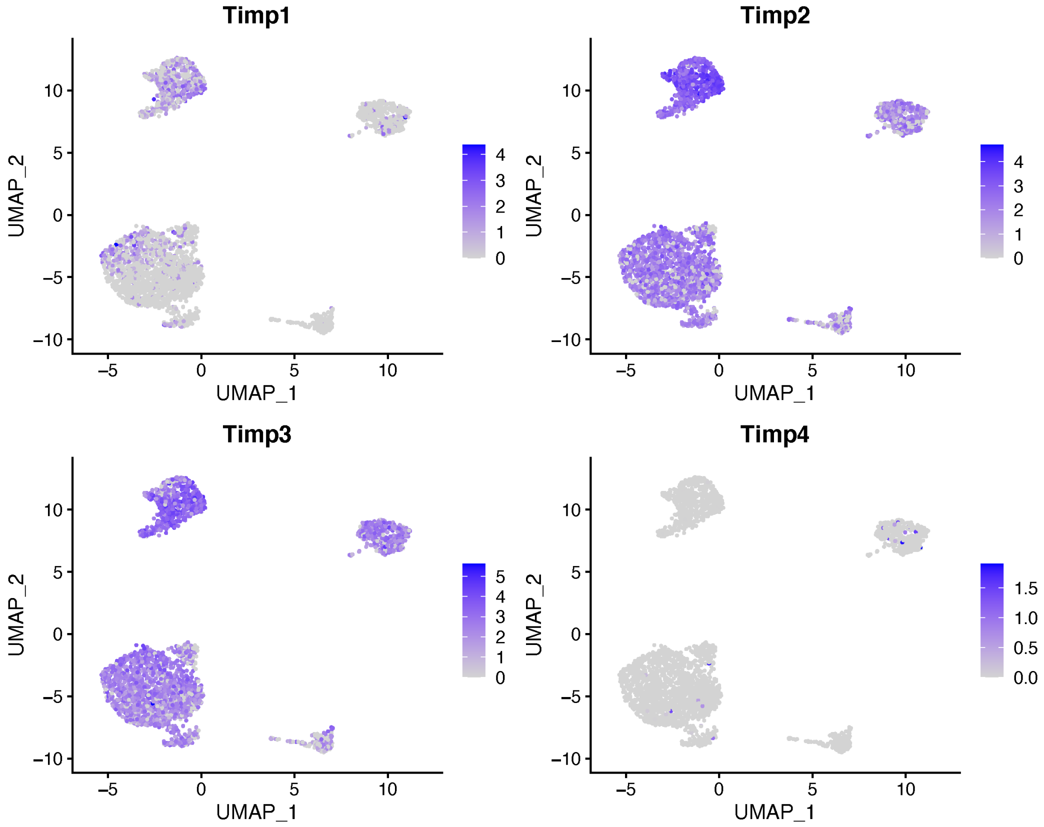

### Mammary Stromal Cells_UMAP Cluster (PC = 40)

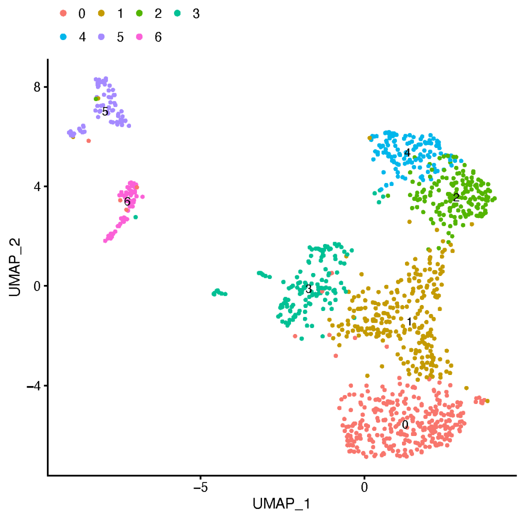


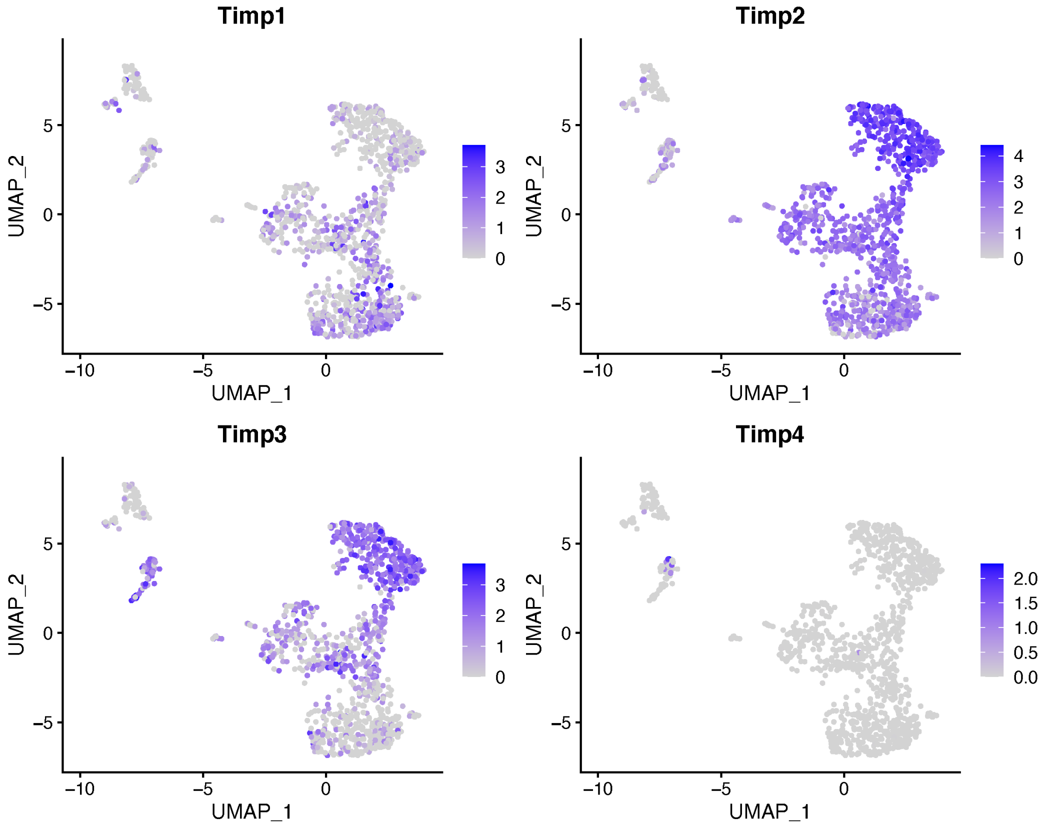

Skeletal Muscle Endothlelial Cells_UMAP Cluster (PC = 40)

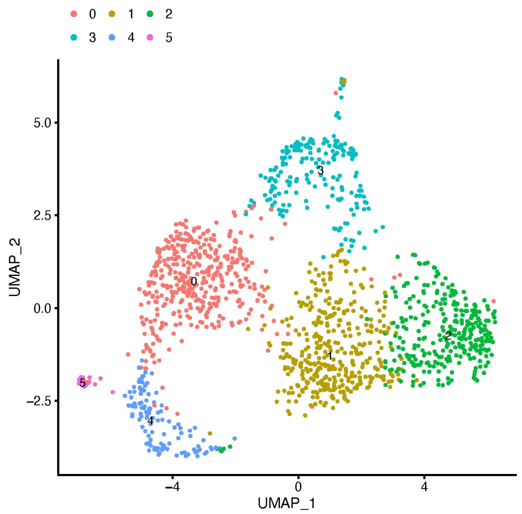


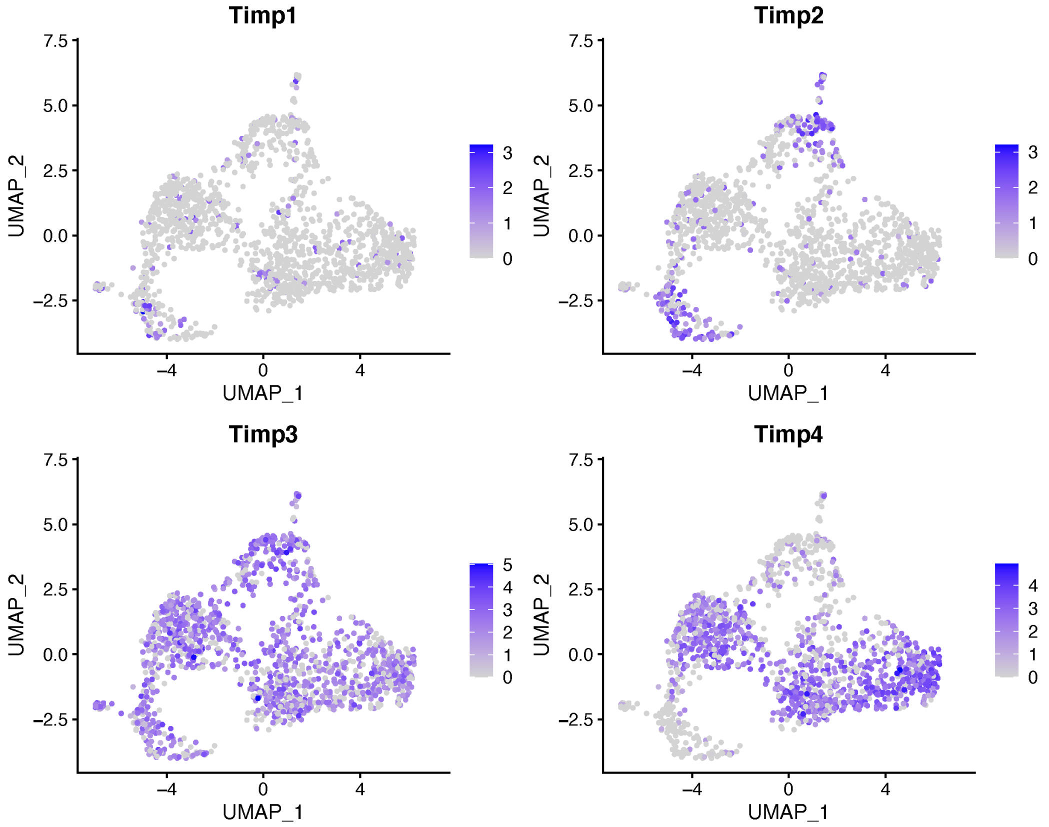

Skeletal Muscle Chondroblasts_UMAP Cluster (PC = 40)

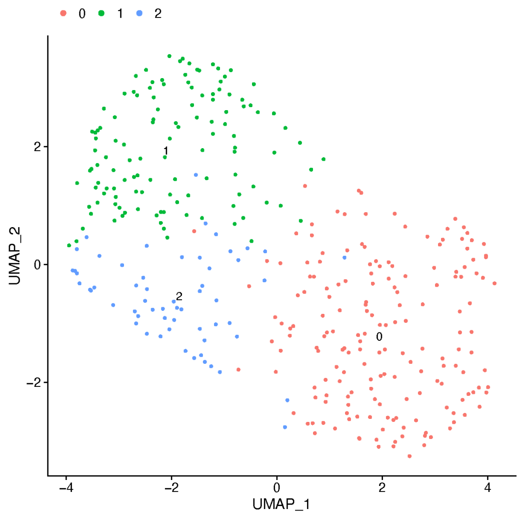


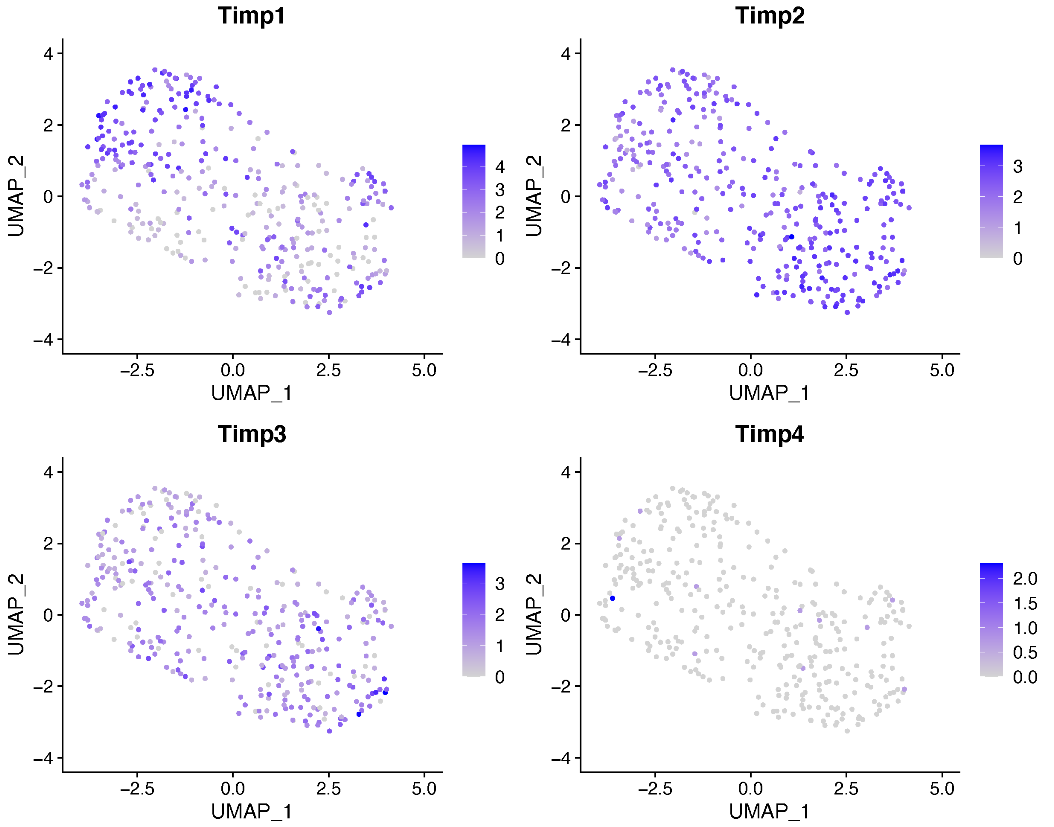

Skeletal Muscle Macrophages_UMAP Cluster (PC = 40)

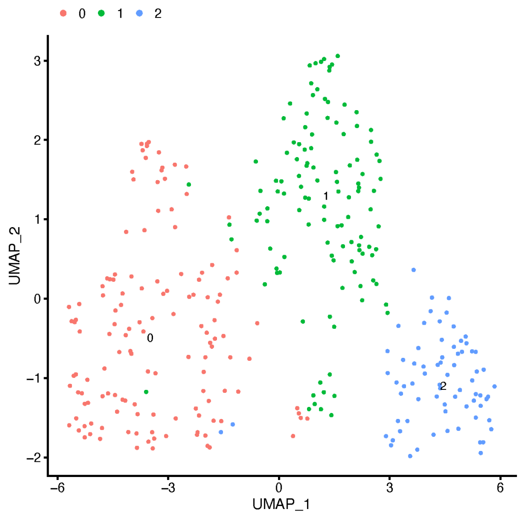


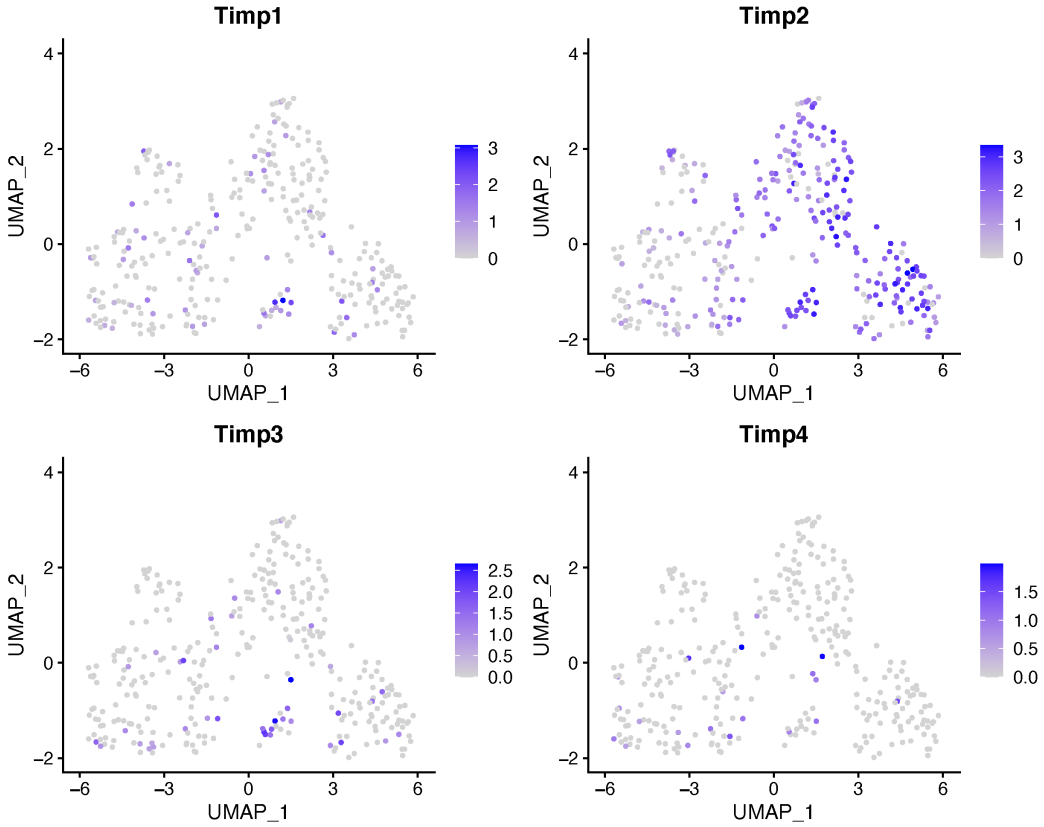

### Trachea Endothelial Cells_UMAP Cluster (PC = 40)

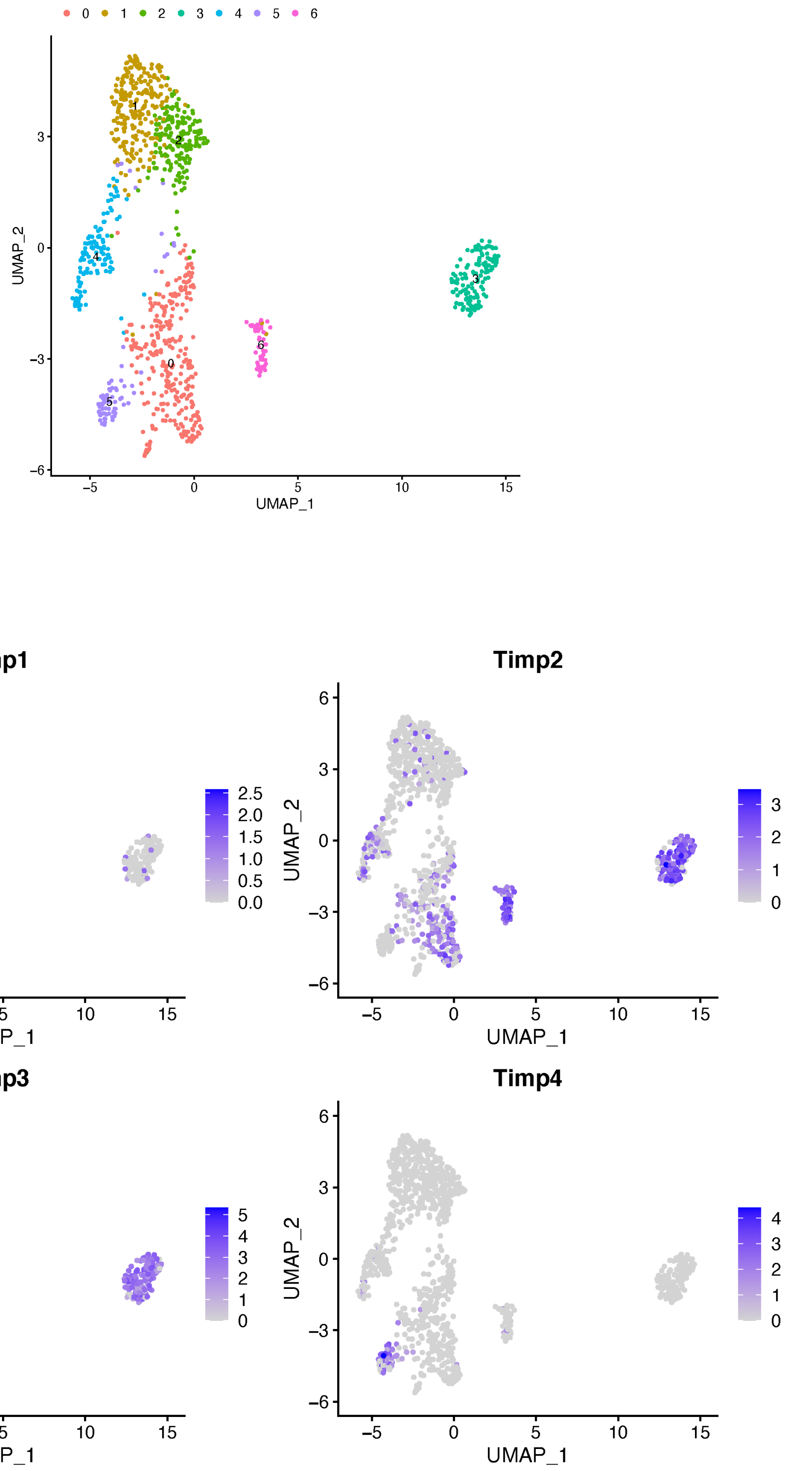

### Trachea Stromal Cells_UMAP Cluster (PC = 40)

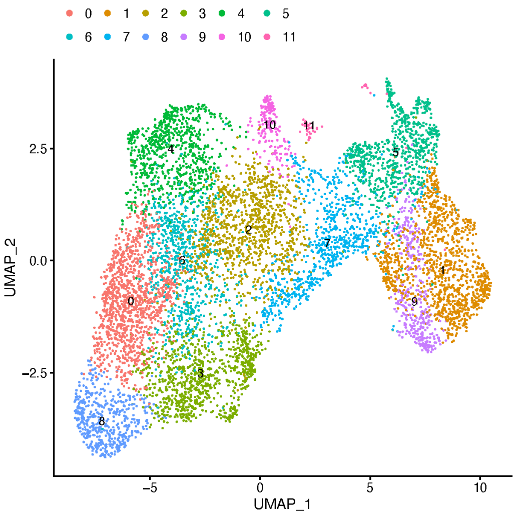


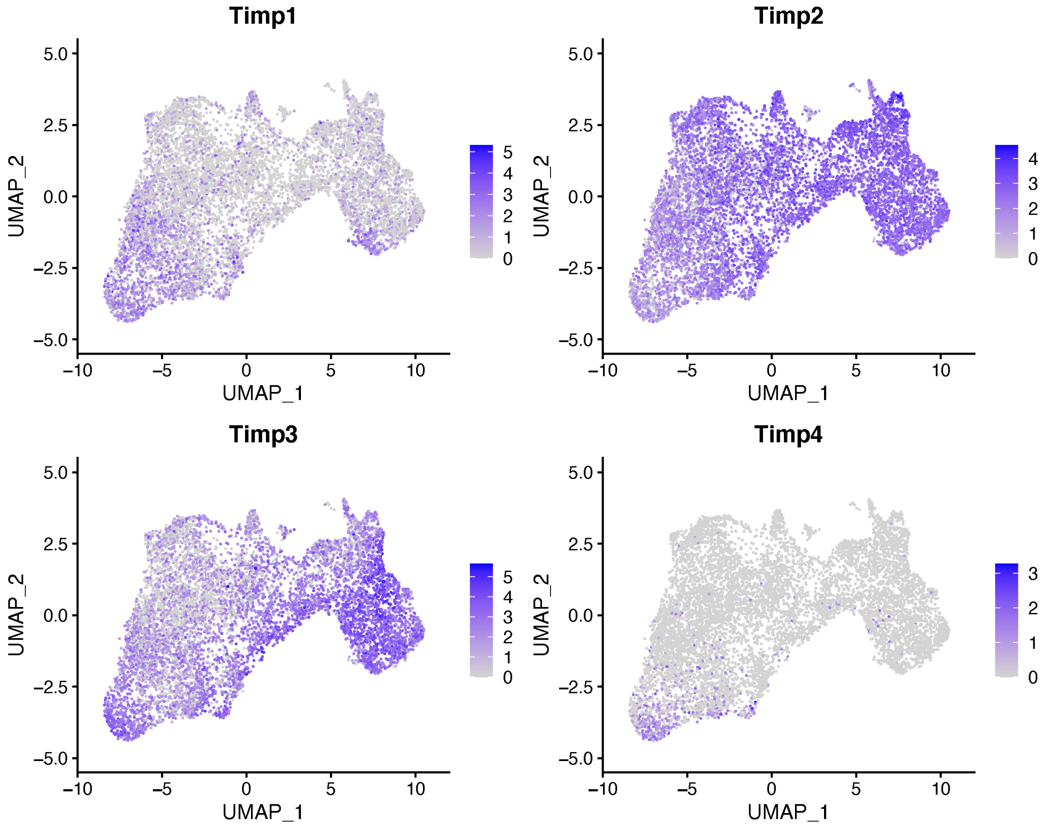
