## Supplemental Figure 4 for "Whole Organism Profiling of the Timp Gene Family"

**Figure S4. RNA ISH of mammary tissue depicting Timp2 expression throughout the fibrous connective tissue.**

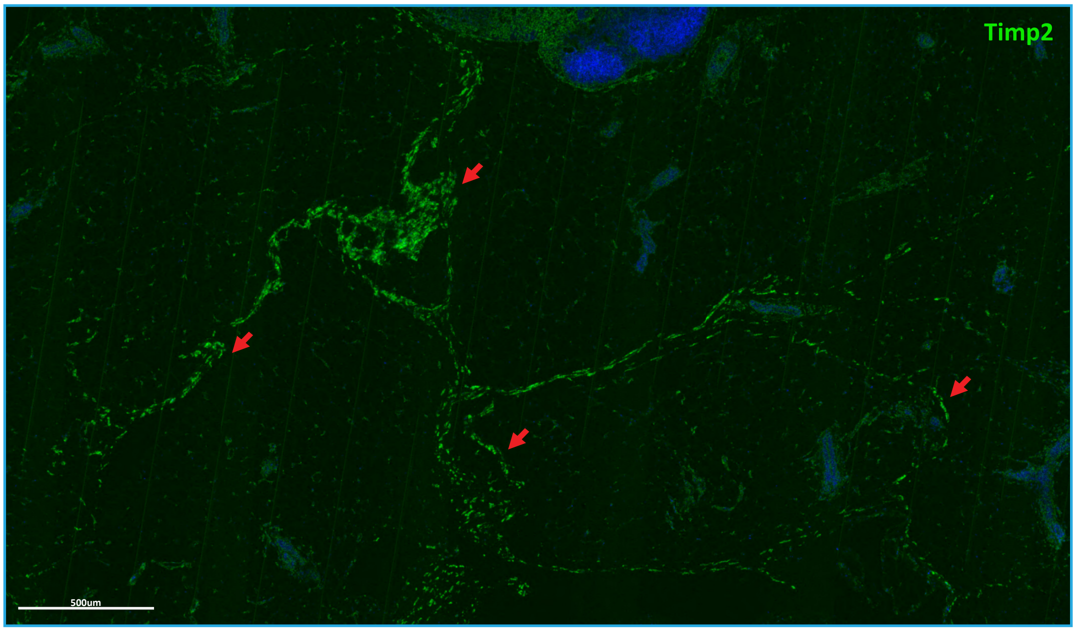


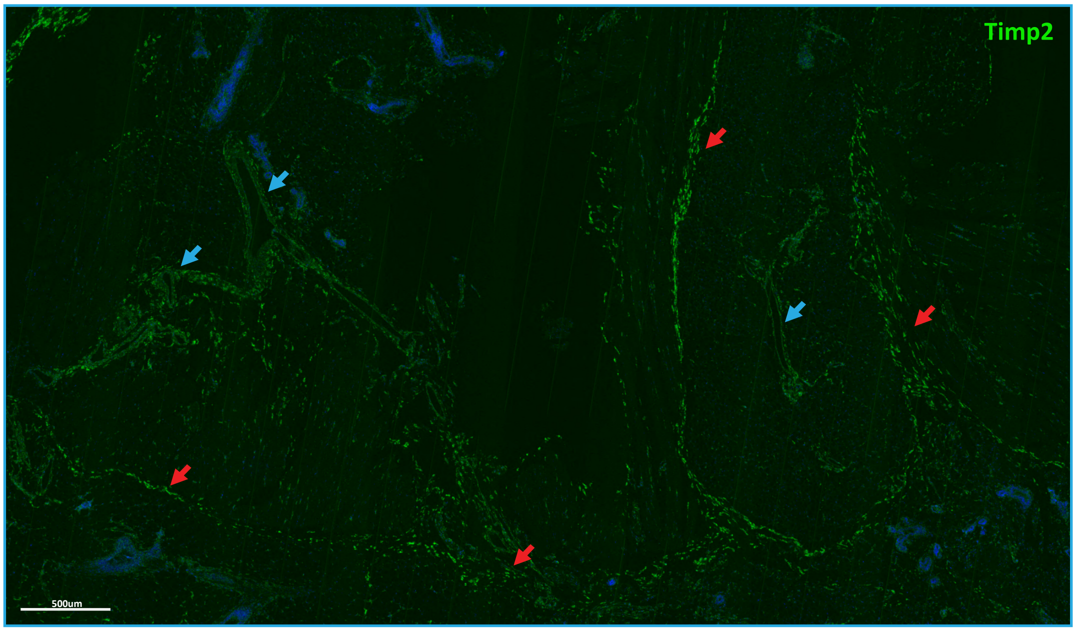
