## Supplemental Figure 5 for "Whole Organism Profiling of the Timp Gene Family"

**Figure S5. RNA ISH of mammary tissue depicting Timp2/3 expression throughout a peripheral nerve and skeletal muscle co-harvested with the mammary tissue.**

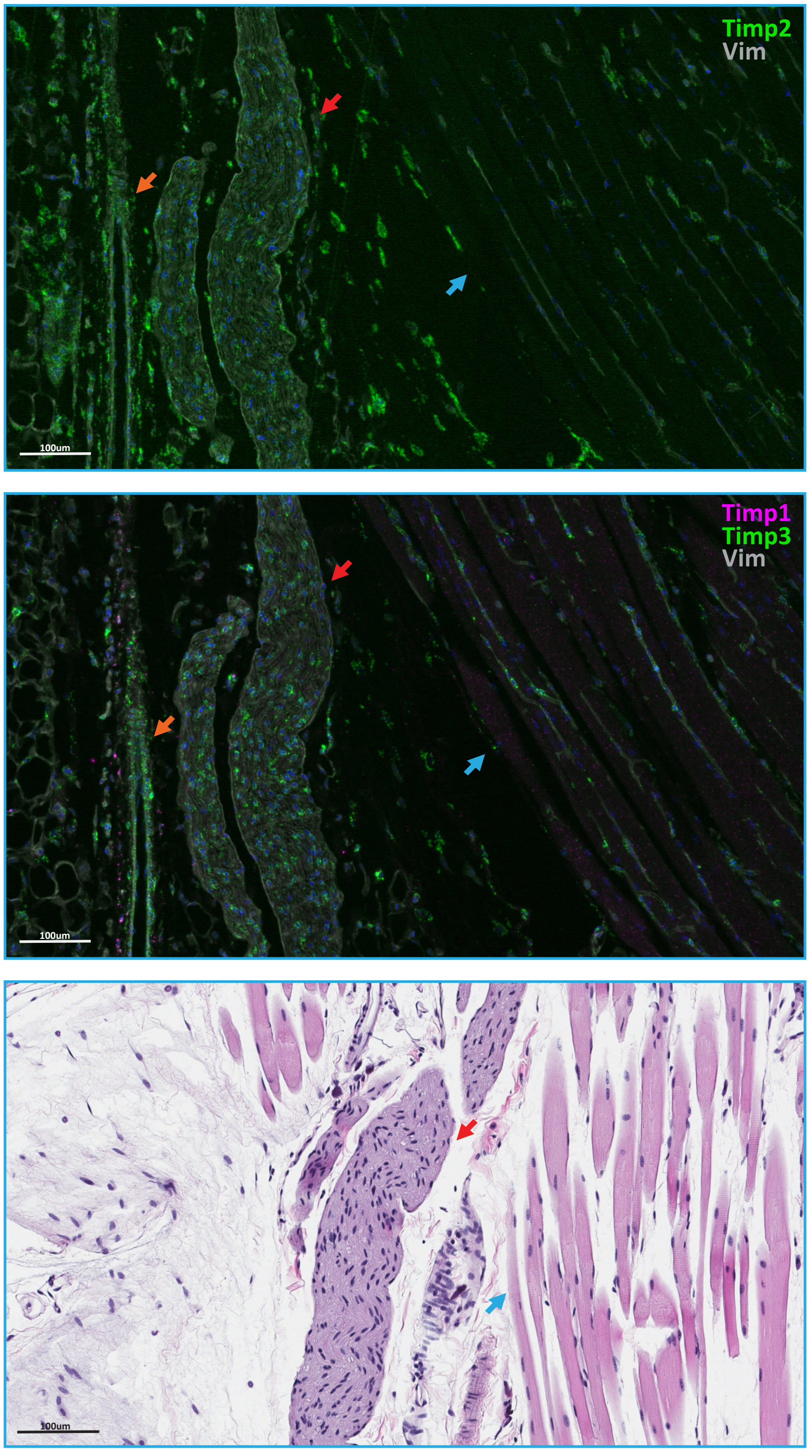
