## Supplementary figures and images for "Whole Organism Profiling of the Timp Gene Family"

### Supplemental Figure 6

**Figure S6. Inflammation of the murine lungs.**

A

B

C

D

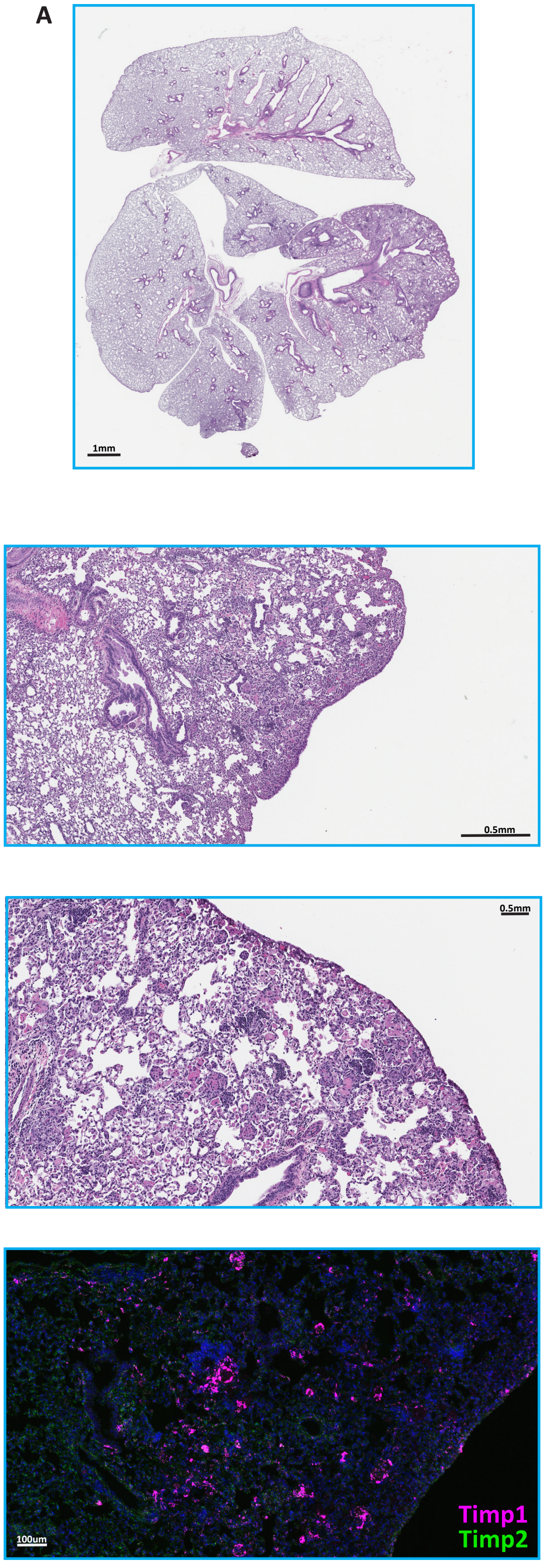

### Supplemental Figure 7

**RNA ISH and H+E the cardiac vessel within the interventricular septum.**

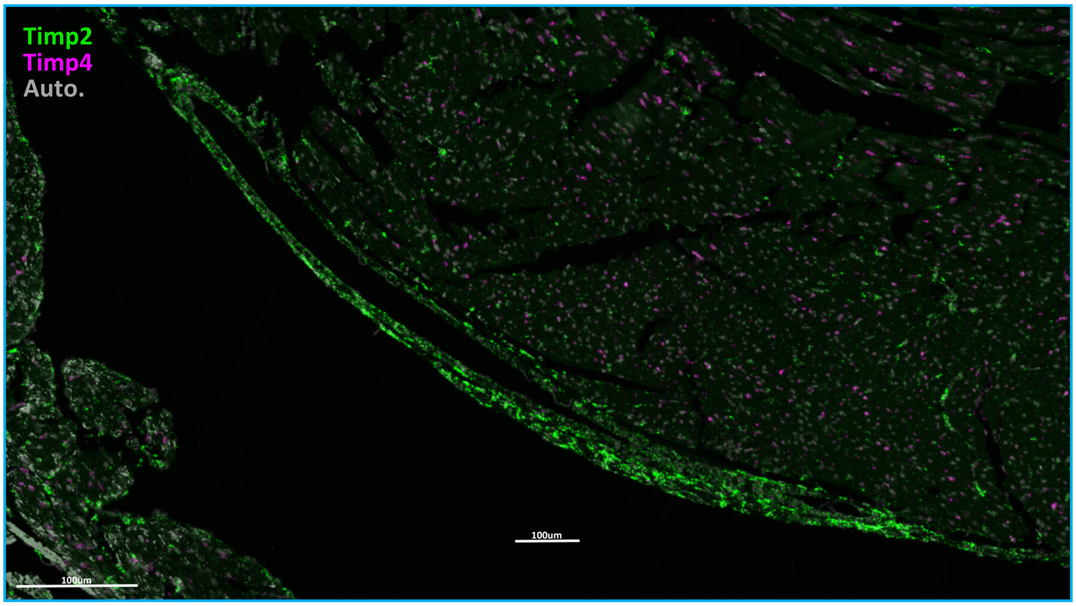


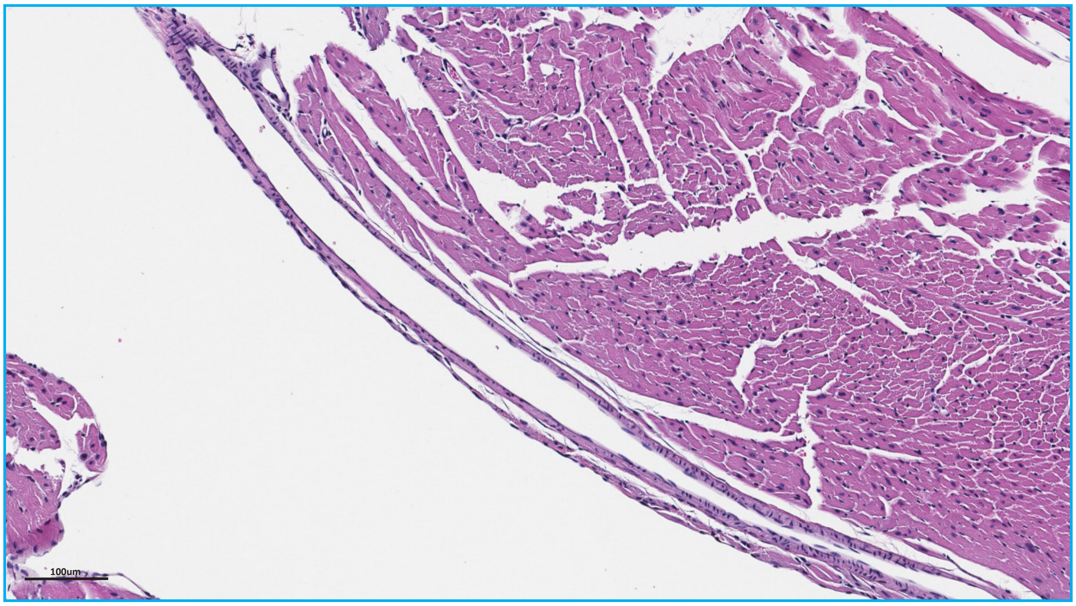
