## Supplemental Figure 8 for "Whole Organism Profiling of the Timp Gene Family"

**Figure S8. RNA ISH/IHC negative control images**

Mammary

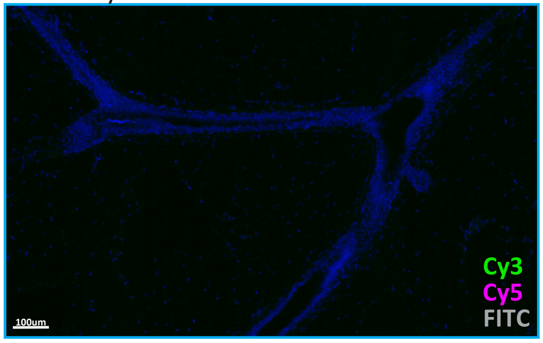


Lung

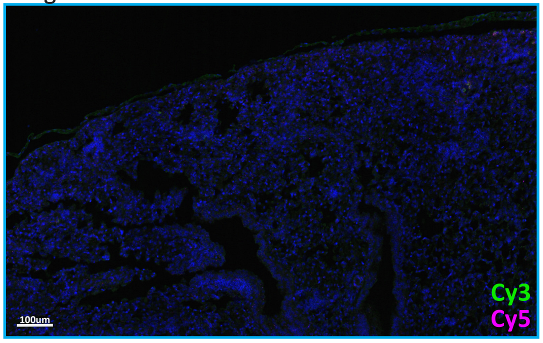


Heart

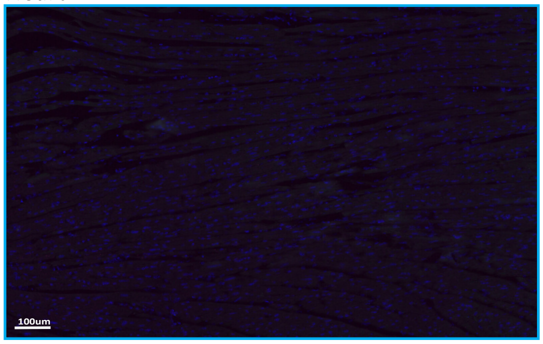


Additional heart

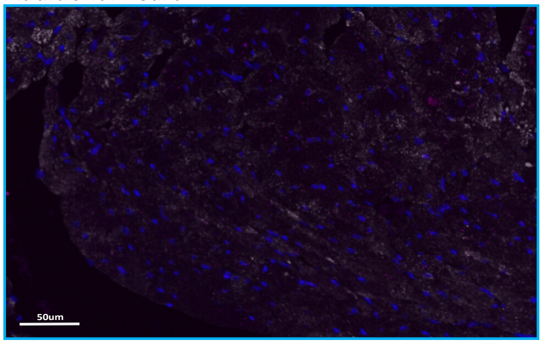


Skeletal Muscle

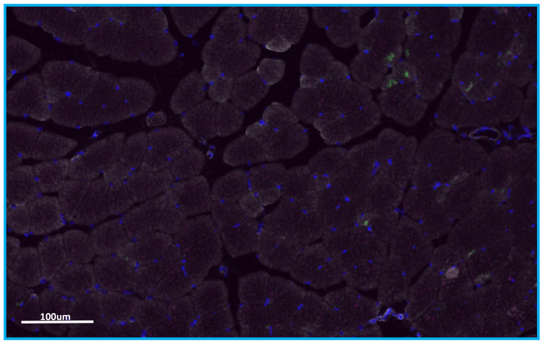
