## Supplementary material for "Whole Organism Profiling of the Timp Gene Family": Supplementary File 1_Example code (1).r

```
## Supplementary File 1_Example code for lung tissue

## Analyze Tabula Muris Lung tissue data
library(Seurat)
library(ggplot2)
library(patchwork)
options(future.globals.maxSize = 4000 * 1024^2)
## read in FACS data
raw_counts_FACS <- read.csv("./FACS/Lung-counts.csv", row.name=1)
dim(raw_counts_FACS)
[1] 23433  1923
FACS_Lung <- CreateSeuratObject(counts = raw_counts_FACS, project = "FACS_Lung")
FACS_Lung
An object of class Seurat 
23433 features across 1923 samples within 1 assay 
Active assay: RNA (23433 features, 0 variable features)
## read in droplet data
droplet_L1.data <- Read10X(data.dir = "./droplet/Lung-10X_P7_8/")
droplet_L2.data <- Read10X(data.dir = "./droplet/Lung-10X_P7_9/")
droplet_L3.data <- Read10X(data.dir = "./droplet/Lung-10X_P8_12/")
droplet_L4.data <- Read10X(data.dir = "./droplet/Lung-10X_P8_13/")
droplet_L1 <- CreateSeuratObject(counts = droplet_L1.data, project = "droplet_L1")
droplet_L2 <- CreateSeuratObject(counts = droplet_L2.data, project = "droplet_L2")
droplet_L3 <- CreateSeuratObject(counts = droplet_L3.data, project = "droplet_L3")
droplet_L4 <- CreateSeuratObject(counts = droplet_L4.data, project = "droplet_L4")
droplet_L1
An object of class Seurat 
23433 features across 625 samples within 1 assay 
Active assay: RNA (23433 features, 0 variable features)
droplet_L2
An object of class Seurat 
23433 features across 1525 samples within 1 assay 
Active assay: RNA (23433 features, 0 variable features)
droplet_L3
An object of class Seurat 
23433 features across 737280 samples within 1 assay 
Active assay: RNA (23433 features, 0 variable features)
droplet_L4
An object of class Seurat 
23433 features across 737280 samples within 1 assay 
Active assay: RNA (23433 features, 0 variable features)

## Note: there is no mitochondrial annotations/reads

## Filter low-quality cells
droplet_Lung <- subset(droplet_1234, subset = nCount_RNA > 1000 & nCount_RNA < 15000 & nFeature_RNA > 500 & nFeature_RNA < 3500)
FACS_Lung <- subset(FACS_Lung, subset = nCount_RNA > 50000 & nCount_RNA < 2000000 & nFeature_RNA > 500 & nFeature_RNA < 4500)

## Normalization
droplet_Lung <- SCTransform(droplet_Lung)
FACS_Lung <- SCTransform(FACS_Lung)
## Integration
object.list <- list(droplet_Lung, FACS_Lung)
Lung.features <- SelectIntegrationFeatures(object.list = object.list, nfeatures = 3000)
Lung.list <- PrepSCTIntegration(object.list = object.list, anchor.features = Lung.features)
Lung.anchors <- FindIntegrationAnchors(object.list = Lung.list, normalization.method = "SCT", anchor.features = Lung.features)
Lung.integrated <- IntegrateData(anchorset = Lung.anchors, normalization.method = "SCT")

identical(Lung.integrated[["RNA"]]@data, Lung.integrated[["RNA"]]@counts)
[1] TRUE
Lung.integrated <- NormalizeData(Lung.integrated, assay="RNA", verbose = FALSE)

## Clustering
Lung.integrated <- RunPCA(Lung.integrated, verbose = FALSE)
pdf("ElbowPlot_Lung.integrated.pdf")
ElbowPlot(Lung.integrated, ndims = 50)
dev.off()
Lung.integrated <- RunUMAP(Lung.integrated, dims = 1:30)
Lung.integrated <- FindNeighbors(Lung.integrated, dims = 1:30, verbose = FALSE)
Lung.integrated <- FindClusters(Lung.integrated, verbose = FALSE)

## Add batch information
batch1 <- c("droplet_L1", "droplet_L2", "droplet_L3", "droplet_L4")
batch2 <- c("FACS")
bat <- c()
for (s in Lung.integrated@meta.data$orig.ident) {
if (s %in% batch1) {bat <- c(bat, "droplet")} else {bat <- c(bat, "FACS")}
}
Lung.integrated@meta.data[, "batch.info"] <- bat
head(Lung.integrated@meta.data)

## UMAP
pdf("Umap_Lung.integrated_pc30.pdf", 10, 10);
plots <- DimPlot(Lung.integrated, label = TRUE)
plots & theme(legend.position = "top") & guides(color = guide_legend(nrow = 2, byrow = TRUE, override.aes = list(size = 2)))
dev.off()
## Feature plot
DefaultAssay(Lung.integrated) <- "RNA"
pdf("FeaturePlot_Lung.integrated_pc30_Timps.pdf", 12, 10);
FeaturePlot(Lung.integrated, features = c("Timp1", "Timp2", "Timp3", "Timp4"), ncol = 2)
dev.off()

MetaData <- Lung.integrated@meta.data
i.droplet <- which(MetaData$batch.info=="droplet")
cells.droplet <- strsplit(rownames(MetaData)[i.droplet], '-')
cells.t <- do.call(rbind, cells.droplet)
cell.ids <- cells.t[,1]
rownames(MetaData)[i.droplet] <- cell.ids

## Add cell type annotations
anno_droplet <- read.csv("annotations_droplets.csv", row.name=1)
anno_FACS <- read.csv("annotations_FACS.csv", row.name=1)
Anno <- rbind(anno_droplet, anno_FACS)[,c(1,2,4)]

Lung.integrated@meta.data[, "cell.type"] <- Anno[rownames(MetaData),2]

## Show cell type annotations on UMAP
pdf("Umap_Lung.integrated_pc30_celltype.pdf", 12, 12);
plots <- DimPlot(Lung.integrated, group.by = "cell.type", label = TRUE)
plots & theme(legend.position = "top") & guides(color = guide_legend(nrow = 3, byrow = TRUE, override.aes = list(size = 2)))
dev.off()

saveRDS(Lung.integrated, file = "Lung.integrated.rds")
```
