## Supplementary material for "Whole Organism Profiling of the Timp Gene Family": Supplementary File 2_Example code (2).r

```
## Supplementary File 2_Example code for lung tissue stromal cells

## Analyze stromal cells from Lung tissue
library(Seurat)
library(ggplot2)
library(patchwork)
options(future.globals.maxSize = 4000 * 1024^2)
## Read in integrated data
Lung <- readRDS(file = "Lung.integrated.rds")
raw_counts <- Lung[['RNA']]@counts
dim(raw_counts)
[1] 23433  6928
MetaData <- Lung@meta.data
identical(rownames(MetaData), colnames(raw_counts))
[1] TRUE
unique(MetaData$cell.type)
 [1] "macrophage"          "B cell"              "natural killer cell"
 [4] "T cell"              "endothelial cell"    "stromal cell"       
 [7] "dendritic cell"      "leukocyte"           "monocyte"           
[10] "unknown"             "type II pneumocyte"  "mast cell"          
[13] "mesothelial cell"    "ciliated cell"       "Clara cell"         
[16] NA                    "epithelial cell"     "type I pneumocyte"            NA     
i.ct <- which(MetaData$cell.type=="stromal cell")
length(i.ct)
[1] 2870
i.facs <- which(MetaData$batch.info=="FACS")
i.droplet <- which(MetaData$batch.info=="droplet")
raw_counts_facs <- raw_counts[,intersect(i.ct, i.facs)]
raw_counts_droplet <- raw_counts[,intersect(i.ct, i.droplet)]

Lung_SC_facs <- CreateSeuratObject(counts = raw_counts_facs, project = "Lung_SC_facs")
Lung_SC_droplet <- CreateSeuratObject(counts = raw_counts_droplet, project = "Lung_SC_droplet")

## Normalization
Lung_SC_droplet <- SCTransform(Lung_SC_droplet)
Lung_SC_facs <- SCTransform(Lung_SC_facs)

## Integration
object.list <- list(Lung_SC_droplet, Lung_SC_facs)
Lung.features <- SelectIntegrationFeatures(object.list = object.list, nfeatures = 3000)
Lung.list <- PrepSCTIntegration(object.list = object.list, anchor.features = Lung.features)
Lung.anchors <- FindIntegrationAnchors(object.list = Lung.list, normalization.method = "SCT", anchor.features = Lung.features)
Lung_SC.integrated <- IntegrateData(anchorset = Lung.anchors, normalization.method = "SCT")

identical(Lung_SC.integrated[["RNA"]]@data, Lung_SC.integrated[["RNA"]]@counts)
[1] TRUE
Lung_SC.integrated <- NormalizeData(Lung_SC.integrated, assay="RNA", verbose = FALSE)

## Clustering
Lung_SC.integrated <- RunPCA(Lung_SC.integrated, verbose = FALSE)
pdf("ElbowPlot_Lung_SC.integrated.pdf")
ElbowPlot(Lung_SC.integrated, ndims = 50)
dev.off()
Lung_SC.integrated <- FindNeighbors(Lung_SC.integrated, dims = 1:30)
Lung_SC.integrated <- FindClusters(Lung_SC.integrated)
Lung_SC.integrated <- RunUMAP(Lung_SC.integrated, dims = 1:30)

## Add batch and cell type information
Lung_SC.integrated@meta.data[, "batch.info"] <- MetaData$batch.info[c(intersect(i.ct, i.droplet), intersect(i.ct, i.facs))]
head(Lung_SC.integrated@meta.data)
Lung_SC.integrated@meta.data[, "cell.type"] <- MetaData$cell.type[c(intersect(i.ct, i.droplet), intersect(i.ct, i.facs))]

saveRDS(Lung_SC.integrated, file = "Lung_SC.integrated.rds")
Lung_SC.integrated <- readRDS(file = "Lung_SC.integrated.rds")

## UMAP
pdf("Umap_Lung_SC.integrated_pc30.pdf");
plots <- DimPlot(Lung_SC.integrated, label = TRUE)
plots & theme(legend.position = "top") & guides(color = guide_legend(nrow = 2, byrow = TRUE, override.aes = list(size = 2)))
dev.off()

## Feature plot
DefaultAssay(Lung_SC.integrated) <- "RNA"
pdf("FeaturePlot_Lung_SC.integrated_Timps.pdf", 10, 8);
FeaturePlot(Lung_SC.integrated, features = c("Timp1", "Timp2", "Timp3", "Timp4"), ncol = 2)
dev.off()

## Perform differential expression analysis
markers <- FindAllMarkers(Lung_SC.integrated, min.pct = 0.1, assay="RNA", slot="data", test.use = "LR", latent.vars="batch.info")
write.csv(markers, "FindAllMarkers_Lung_SC.integrated.csv", row.names=T, quote=F);
markers.f <- markers[which(markers$p_val_adj <0.05),]
write.csv(markers.f, "FindAllMarkers_Lung_SC.integrated_fdr05.csv", row.names=T, quote=F);

table(Lung_SC.integrated@meta.data$seurat_clusters)
  0   1   2   3   4   5   6   7   8 
591 494 422 408 388 189 162 109 107

markers <- FindMarkers(Lung_SC.integrated, ident.1=6, min.pct = 0.1, assay="RNA", slot="data", test.use = "LR", latent.vars="batch.info")
write.csv(markers, "FindMarkers_Lung_SC.integrated_clu6.csv", row.names=T, quote=F);
markers.f <- markers[which(markers$p_val_adj <0.05),]
write.csv(markers.f, "FindMarkers_Lung_SC.integrated_clu6_fdr05.csv", row.names=T, quote=F);

markers <- FindMarkers(Lung_SC.integrated, ident.1=c(2,5), min.pct = 0.1, assay="RNA", slot="data", test.use = "LR", latent.vars="batch.info")
write.csv(markers, "FindMarkers_Lung_SC.integrated_clu25.csv", row.names=T, quote=F);
markers.f <- markers[which(markers$p_val_adj <0.05),]
write.csv(markers.f, "FindMarkers_Lung_SC.integrated_clu25_fdr05.csv", row.names=T, quote=F);
```
